# A responsivity theory of call timing patterns in bat echolocation

**DOI:** 10.1101/2025.06.16.659925

**Authors:** Ravi Umadi, Uwe Firzlaff

## Abstract

**Background:** The temporal patterning of bat echolocation is extensively characterised, but a general mathematical account linking call timing to acoustic propagation, relative motion, and behaviourally selected information remains incomplete.

**Results:** We present the responsivity framework, which represents each call event as a delayed active-sensing cycle comprising an acoustic acquisition interval and a subsequent behavioural interval. Their proportional relation is described by the responsivity coefficient, *k*_*r*_, connecting call timing to effective anchor distance, relative velocity, call duration, and the selected echo window. Analytical derivation, call-by-call simulation, internal validation, and comparison with field recordings show how the framework generates hyperbolic call-rate–distance relationships, shifts approach timing with echo-window selection, and places the terminal buzz at the near-target limit of the same timing process. Responsivity and kinematics have separable terminal effects: *k*_*r*_ constrains terminal update density and attainable call rate, whereas closing velocity determines the time available within the terminal region. A buzz-readiness state precedes the complete expression of buzz, and transient sonar strobe groups emerge under multi-target anchor allocation without an explicit grouping rule. Field recordings show broad correspondence with simulations in the organisation of movement, call timing and call duration, and illustrate conditional evaluation of acquisition-window and anchor assumptions.

**Conclusions:** Responsivity theory provides a process-based descriptor of echolocation timing and a diagnostic framework for hypothesis generation and empirical testing. The framework identifies the measurements needed to test how selected acoustic information governs subsequent call timing.

## 1 BACKGROUND

Dynamic behavioural adaptations depend on detecting relevant information, determining when another observation is needed, and deciding when the information already acquired is sufficient to support action (Schroeder et al., 2010; Yang et al., 2016). In active sensing, this time-domain problem is especially clear because each sampling act measures the environment, and influences the conditions under which the next sample will be obtained. Echolocating bats provide a remarkable example: each vocal emission initiates an acoustic sampling event, and the emission times provide directly observable markers of the sampling sequence (Griffin, 1944, 2001; Griffin and Galambos, 1941; Griffin et al., 1960). A call sequence is therefore more than an acoustic description of behaviour; it is a record of how an animal allocates opportunities to acquire new sensory information.

The temporal patterning of bat calls has been extensively measured and described across search, approach, tracking, and interception (Kalko and Schnitzler, 1989; Moss and Surlykke, 2001; Simmons et al., 1979). Bats alter call rate, duration, intensity, spectral structure, beam direction, and flight behaviour with target range, clutter, prey motion, interference, and task difficulty (Falk et al., 2014, 2015; Geberl et al., 2015; Jakobsen and Surlykke, 2010; Moss and Surlykke, 2010; Surlykke et al., 2009; Wohlgemuth and Moss, 2013). Organised sonar strobe groups have also been observed in call sequences during demanding tasks (Hulgard and Ratcliffe, 2016; Kothari et al., 2014, 2018), while bats can accumulate information across successive acoustic snapshots and coordinate acoustic gaze with flight and target selection (Aihara et al., 2013; Fujioka et al., 2016; Ghose et al., 2006; Ghose and Moss, 2006; Nishiumi et al., 2024; Salles et al., 2020; Sumiya et al., 2017). These observations establish that sonar timing is adaptive, but a temporal pattern does not by itself identify the control process that generated it. Similar sequences could reflect deliberate call grouping, changing acoustic delay, movement through the scene, selection among competing echoes, or interactions among these processes. Thus, a timing law that connects such behavioural organisation to acoustic propagation, relative motion, and the finite opportunity for sensory updating remains to be formalised.

At the centre of this unresolved timing law is a long-standing question in echolocation research: which event in the returning echo stream governs the timing of the next call? A natural scene does not return a single isolated echo, but a cascade from prey, vegetation, ground, conspecifics, and other structures. The behaviourally relevant event might be the first return, the strongest or most expected echo, or the point at which a selected portion of the stream has supplied sufficient information. Neural sensitivity to early-arriving echoes is compatible with a first-arrival rule (Beetz et al., 2016, 2017, 2026; Greiter and Firzlaff, 2017), whereas behavioural studies show that bats can direct acoustic gaze and distribute sampling among several objects (Fujioka et al., 2016; Sumiya et al., 2017; Surlykke et al., 2009). Echo precedence, perceptual selection, and the echo that controls the next interpulse interval are therefore related but distinct. A useful timing theory should not require that this effective acoustic anchor be known in advance; it should make alternative anchors and information-window endpoints empirically testable.

Distance is only one component of this timing problem. Relative closing velocity determines how acoustic geometry changes between calls and how long an animal remains within a spatial region. Speed and reaction time influence interception more generally (Wilson et al., 2013), while velocity-dependent acoustic control has been studied most directly through Doppler-related behaviour in constant-frequency echolocators (Bell and Fenton, 1984; Perrine, 1944; Schoeppler et al., 2023; Trappe and Schnitzler, 1982; Yoshida et al., 2024). Its role in the temporal organisation of frequency-modulated approach sequences has received less attention, although flight velocity is increasingly recognised as an important driver of aerial-hawking behaviour (Jakobsen et al., 2025) and information-update rate is emerging as a broader problem across sensory systems (Ma et al., 2025; Taub et al., 2023). The terminal buzz makes these coupled constraints conspicuous: echo delay and call duration contract as the bat closes on a target, call rate rises, and the time remaining for further updates collapses. Yet closing velocity alone need not set the attainable call rate. A complete account must distinguish the density at which updates are generated from the time available to express them.

Here, we introduce the *responsivity framework*, a new theory in which *responsivity* denotes the coupling between the time required to acquire behaviourally relevant acoustic information and the timing of the subsequent call. Each call is treated as a time marker in a delayed active-sensing loop. The interval preceding the next call is separated into an acoustic acquisition interval, *T*_*a*_, which ends when the selected acoustic information becomes available, and a behavioural interval, *T*_*b*_, representing the additional temporal allocation before the next emission. Their proportional relation is described by the dimensionless responsivity coefficient, *k*_*r*_. This coefficient is an effective behavioural scaling term rather than a specified neural or motor latency. Sound propagation, relative motion, call duration, and the selected echo-window endpoint determine the physical opportunity for acquisition; responsivity determines how that opportunity is temporally allocated. Once *k*_*r*_ is specified, an observed interpulse interval, or its reciprocal call rate, can therefore be partitioned directly into *T*_*a*_ and *T*_*b*_, making the latent acoustic and behavioural components of each interval analytically accessible. The resulting identities define callrate–distance and call-rate–velocity relationships; under specified information-window and kinematic assumptions, they can also be rearranged to estimate effective anchor distance or radial closing velocity and to derive terminal call-rate ceilings, update budgets, and buzz-readiness metrics. Conversely, the same relations provide a route to quantifying *k*_*r*_ from behavioural data, either by fitting call-resolved timing to measured anchor distance and radial closing velocity under a specified information-window rule, or from an attained terminal call-rate ceiling when stop distance and closing velocity are known.

The framework leads to a set of linked predictions. First, changing the endpoint of acoustic acquisition should shift interpulse intervals, behavioural intervals, and call-duration trajectories even when responsivity is unchanged. Second, *k*_*r*_ and radial closing velocity should have separable terminal effects: responsivity should strongly constrain update density and the attainable terminal rate, whereas velocity should primarily determine how rapidly the terminal region is traversed and hence how long a buzz can persist. Third, the approach to the terminal ceiling should define a state of buzz readiness before a complete buzz is expressed. Finally, if categoric temporal patterns and phases are observable organisations of a common call-to-call rule rather than mechanisms in themselves, intermittent call-grouping should be able to emerge when calls are redistributed among several acoustic anchors without requiring a separate group-specific timing programme.

We evaluate these predictions through analytical derivation, call-by-call simulation, internal validation, and comparison with field recordings from free-flying bats. Parameterised simulations isolate the effects of responsivity, relative velocity, echo-window choice, call duration, and multi-target anchor allocation; the validation tests whether the implemented event sequence preserves the derived identities. The field recordings provide an empirical comparison in a shared inter-call-displacement coordinate. We first compare directly observed timing, movement, velocity, and call duration, then match the field and simulated velocity distributions to determine whether kinematics explain their remaining timing differences. Finally, under explicitly fixed responsivity and geometric assumptions, we profile candidate echo-window fractions to show how the framework can diagnose the latent conditions required to reconcile field and model distributions. Together, these analyses ask whether the familiar terminal buzz, its preceding readiness state, and transient call groups can be understood as different expressions of a common delayed active-sensing process. More broadly, they shift the unit of explanation from the name assigned to a call pattern to the event structure that generates each successive sensory update.

## 2 METHODS

The framework treats the rise in call rate during target approach—which is approximately described by an inverse-distance relation—as the observable consequence of successive closed-loop sampling cycles, not as a trajectory that bats are assumed to follow. We formalise those cycles by defining the geometry between the caller and an effective acoustic anchor, decomposing each interpulse interval into an acoustic acquisition interval, *T*_*a*_, and a behavioural interval, *T*_*b*_, and deriving the corresponding timing, kinematic, and terminal-state identities. The Supplementary Information at the end of this document provides a consolidated notation glossary and an application-oriented reference for the framework’s parametric interdependencies, conditional inferences, and underlying assumptions (Supplementary Tables S1–S4).

### 2.1 Spatial geometry of an approach

At the onset of call *n*, let the bat position be x_*b,n*_ and the position of the effective acoustic anchor be x_*q,n*_. The acoustic anchor may be a prey item, clutter object, target component, first-arriving reflector, expected echo feature, or selected portion of a composite echo stream. The displacement vector from the bat to this anchor is

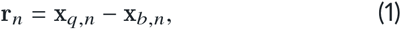

and the corresponding range is

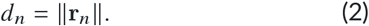

The unit vector pointing from the bat to the anchor is

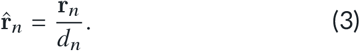

Let the bat and anchor velocities at call onset be v_*b,n*_ and v_*q,n*_, respectively. The relative velocity of the anchor with respect to the bat is

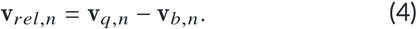

We define radial closing velocity as positive when range decreases,

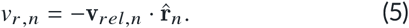

Thus, positive *v*_*r,n*_ denotes closing range, zero denotes no radial closure, and negative values denote increasing range. In continuous notation, the corresponding range dynamics are

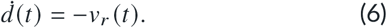

This spatial setup is useful because the same variables describe movement during the call, during echo propagation, during behavioural processing, and across the full interpulse interval.

### 2.2 The call-event model

The event clock is tied to call onset. We denote the onset of call *n* as 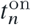, its duration as *T*_*c,n*_, and its offset as

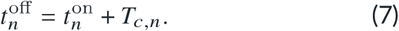

The next call begins at 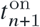. The interpulse interval (IPI) following call *n* is therefore the onset-to-onset interval

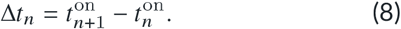

The offset-to-onset silent interval is

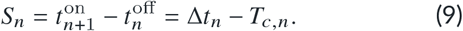

This distinction is important because measured IPI usually refers to call onset-to-onset timing, whereas the period after call offset is the interval in which self-emission no longer occupies the receiver.

Let the first relevant echo from the acoustic anchor reach the bat at 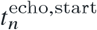, and let the behaviourally relevant echo-information window end at 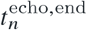. We define the acoustic acquisition interval as the time from call onset to the end of the echo information used to govern the next call,

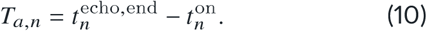

The behavioural interval is the remaining time between acquisition of the relevant acoustic information and the onset of the next call,

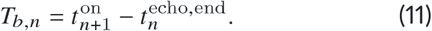

Combining Equations 8, 10, and 11 gives the event-level decomposition

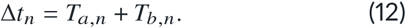

Because the bat and anchor can move during this interval, the distance associated with the next call is not identical to the distance that initiated the current sensorimotor cycle. From the range dynamics in Equation 6, the range at the next call onset is

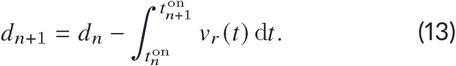

Under a locally constant radial closing velocity across the call cycle, this reduces to

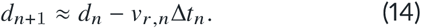

Equation 13 can be partitioned over the call duration, acoustic acquisition interval, and behavioural interval if the movement within each event segment is of interest. Here it serves to make explicit that the call-event model is embedded in a moving trajectory: the same IPI that is generated by acoustic and behavioural timing also determines how far the bat-anchor geometry advances before the next call. The interval between calls *n* and *n*+ 1 is therefore determined by the acoustic state initiated by call *n*, because call *n* generated the echo stream that constrains the timing of call *n*+ 1. When an observed IPI is related to target position, the appropriate reference is the distance to the effective acoustic anchor at the onset of call *n*, not necessarily the distance at call *n* +1. By the time the next call is emitted, the bat and target may already have moved.

We use the term *effective acoustic anchor* to denote the echo-generating object, object component, or integrated portion of the returning echo stream that governs a given call interval. The model does not require the anchor to be specified a priori as the nearest object, the strongest echo, the first-arriving echo, or the experimentally designated target. All such alternatives enter the model through the effective value of *T*_*a,n*_. The identity of the acoustic anchor is therefore treated as an experimentally testable question rather than as a fixed assumption of the mathematical framework.

The resulting event structure, interval decomposition, and trajectory geometry are summarised in Figure 1.

**Figure 1.**
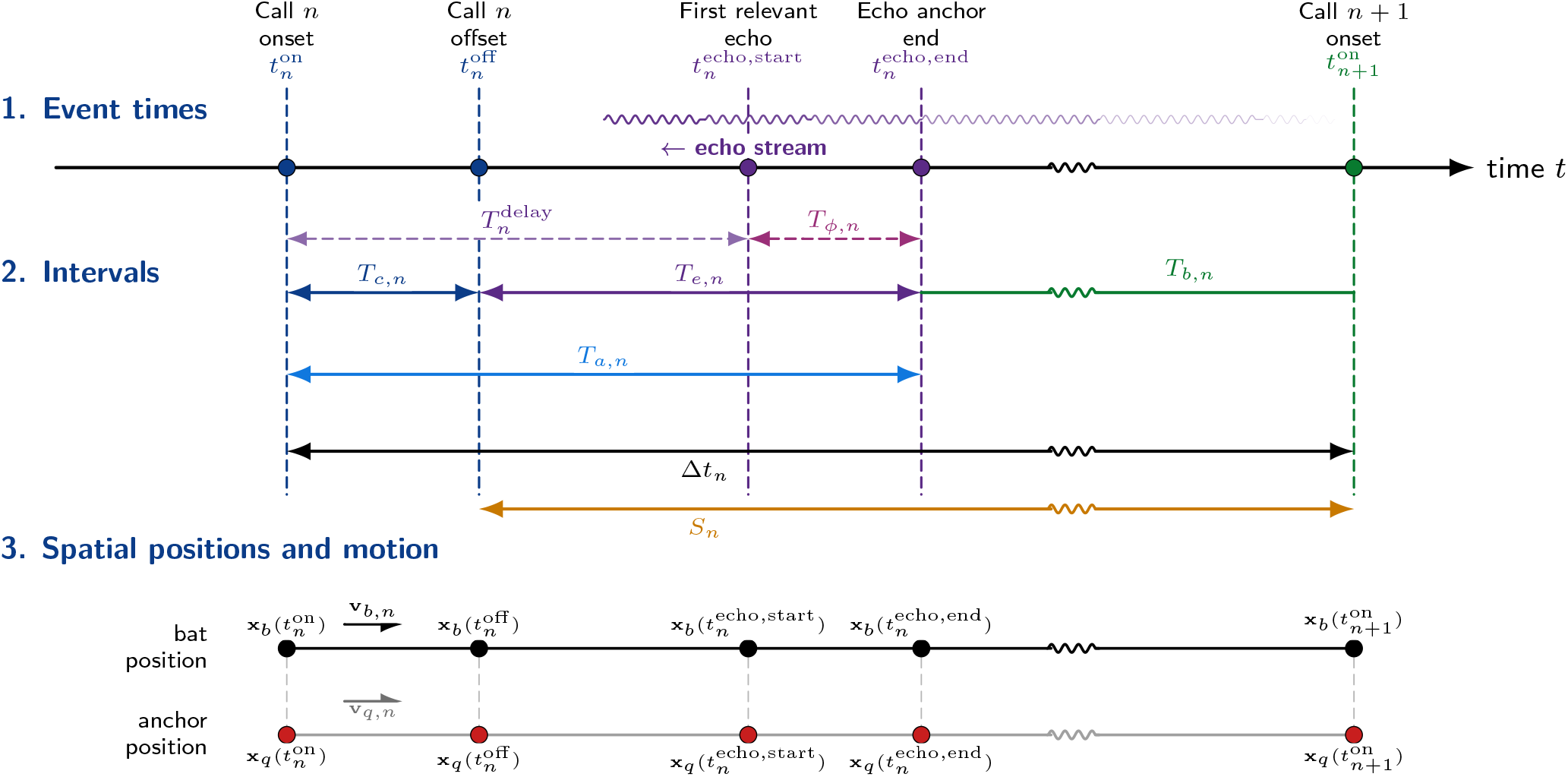
Event timeline, interval decomposition, and spatial geometry for one echolocation cycle. The framework fixes the interpulse interval, Δ*t*_*n*_, as the onset-to-onset interval between call *n* and call *n* + 1. Call duration, *T*_*c,n*_, occupies the first part of this interval, while the returning echo stream defines an acoustic acquisition interval, *T*_*a,n*_, from call onset to the behaviourally selected echo-anchor end. This acquisition interval can include the propagation delay to the first relevant echo, 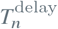, and a selected echo-stream window, *T*_*ϕ,n*_. The remaining time before the next call is the behavioural interval, *T*_*b,n*_, and the offset-to-onset silent interval is *S*_*n*_. The lower spatial schematic shows that bat and anchor positions continue to evolve throughout the same event sequence, so the range associated with call *n* is the range at the beginning of the sensorimotor cycle, whereas the geometry at call *n* + 1 has already advanced along the trajectory.

### 2.3 The acoustic acquisition interval, *T*_*a*_, as a master variable

The acoustic acquisition interval in Equation 10 is a master interval. Its simplest constituent is the delay from call onset to the first relevant echo arrival. If the bat and anchor are separated by distance *d*_*n*_ at call onset and the speed of sound is *c*, then the stationary approximation is

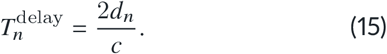

During pursuit, range changes while sound is travelling. Using the radial closing velocity defined in Equation 5, the motion-aware approximation for this delay is

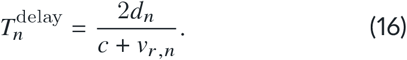

For stationary prey, *v*_*r,n*_ is the bat’s radial flight speed towards the target. For moving prey, it is the radial component of relative velocity between the bat and prey. The condition *v*_*r,n*_ = 0 therefore denotes zero radial closing speed, not necessarily a stationary target.

The first-arrival delay need not exhaust the acoustic acquisition interval. A finite-duration call generates a returning echo packet, and the bat may act on the first informative echo, integrate part of the packet, or wait until a selected portion of the stream has become available. We therefore denote the echo-stream information window used for anchoring as *T*_*ϕ,n*_. A convenient call-duration-linked form is

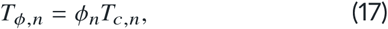

where ϕ_*n*_ is a positive dimensionless echo-window scaling parameter. In the analyses presented here we use ϕ = 0, 0.5, and 1 as convenient reference cases spanning first-arrival anchoring, partial integration of the echo stream, and integration of the full call-defined echo packet, respectively; however, the formulation itself does not require ϕ to be bounded above by one. The master acoustic acquisition interval is then

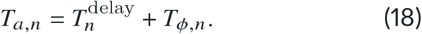

Using Equation 16, this becomes

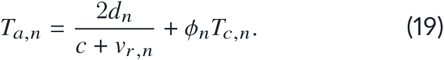

Equation 19 keeps the behavioural definition of the acoustic anchor separate from the propagation delay. The first-arrival model is recovered when the information-window term is negligible. More generally, the same formalism can represent a selected echo feature, a complete call-packet integration window, or a prospectively expected anchor time. Equation 19 is therefore the forward physical form of the acoustic acquisition interval: once anchor distance, radial closing velocity, call duration, and the anchoring rule are specified, *T*_*a*_ can be computed directly from the geometry and event structure of the call cycle.

### 2.4 Call duration as an acoustic information aperture

Call duration, *T*_*c*_, is not treated only as an emitted-sound duration or as a pulse–echo overlap constraint. A call of duration *T*_*c,n*_ occupies a spatial length

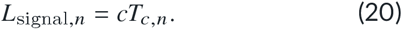

Echoes from reflectors separated in range by less than half of this length overlap in time. The corresponding overlap depth is

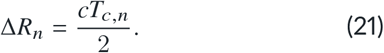

Thus, call duration regulates the spatial depth of the acoustic scene admitted into the returning echo stream. Longer calls generate broader echo packets containing more potentially overlapping reflections; shorter calls restrict the packet and may reduce perceptual load. This provides a mechanistic reason for call-duration decrement during approach: shortening the call not only reduces pulse–echo overlap risk, but can also narrow the sensory information window represented by *T*_*ϕ,n*_ in Equation 17.

It is sometimes useful to describe the post-call interval from call offset to echo-information completion. We denote this offset-referenced interval as

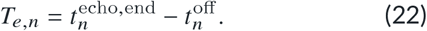

Because *T*_*a*_ is measured from call onset, this quantity is a constituent decomposition rather than a separate definition:

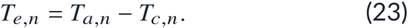

This distinction prevents double-counting. If *T*_*a*_ is onset-referenced, call duration is already part of the event timeline. It enters the timing model through the echo-stream information window, not as an automatic additional delay.

### 2.5 The behavioural interval, *T*_*b*_

The behavioural interval, *T*_*b*_, should not be interpreted as a simple reaction time. It is an aggregate biological interval that may include auditory transduction, afferent transmission, sensory integration, echo selection, decision-making, motor planning, vocal preparation, and optional behavioural lag. We write this conceptually as

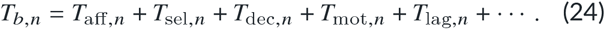

The individual terms denote afferent, selection or integration, decision, motor, and voluntary or stochastic lag components. They need not be strictly serial and may overlap physiologically. Equation 24 is therefore not intended as a complete neural model; it is a statement that the postacquisition portion of the IPI is biological, composite, and potentially behaviourally regulated.

During target approach, bats progressively shorten their interpulse intervals and call durations as target distance decreases (Kalko, 1995; Kalko and Schnitzler, 1989). Bats can also initiate an approach with distance-appropriate call durations and intervals, indicating coordinated organisation of acoustic sampling with target distance (Amichai and Yovel, 2017). These observations motivate a simple effective description of the relationship between acoustic acquisition and the subsequent behavioural interval. Within the behavioural state or approach segment being analysed, we approximate the aggregate interval allocated to processing newly acquired information and preparing the next emission as proportional to the acoustic acquisition interval:

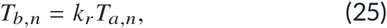

where *k*_*r*_ is a dimensionless responsivity coefficient. It describes the behavioural interval allocated per unit acoustic acquisition time.

This approximation describes the aggregate temporal allocation, including any voluntary or stochastic lag, rather than requiring each constituent process in Equation 24 to scale individually with acoustic acquisition time. Inverse-distance scaling is expected under Equation 25 when the acquisition interval scales approximately with distance over the same segment and *k*_*r*_ is approximately constant. The coefficient need not remain constant across changes in behavioural state.

Substituting the motion-aware acquisition interval from Equation 19 gives

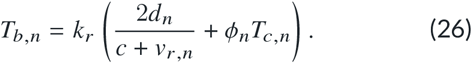

When the information-window term is negligible, this reduces to the first-arrival motion-aware form

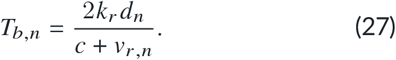

This proportionality also means that once *k*_*r*_ is specified, *T*_*b*_ can be written either in motion-aware geometric form by substituting Equation 19 into Equation 25, or recovered directly from observed IPI or call rate through the identities derived below.

This proportionality should not be read as implying that all physiological processes contributing to *T*_*b*_ are constant. Instead, *k*_*r*_ is treated as an effective systems parameter. One possible mechanistic interpretation is that bats regulate the quantity of sensory information entering the processing loop. As calls shorten during approach, Equations 20 and 21 imply that the spatial depth and duration of the returning echo stream decrease. If processing time scales with the amount of information acquired, then the aggregate behavioural interval can remain approximately proportional to *T*_*a*_ even though the underlying neural and motor components vary from call to call.

Coordinated regulation of emission behaviour and the information admitted for processing could therefore provide a biological basis for the effective proportionality in Equation 25. This information-regulation account is a proposed mechanism rather than a demonstration of the quantitative relation. The coefficient *k*_*r*_ summarises the behavioural interval allocated per unit acquisition time without identifying the contributions of individual neural, motor or scheduling processes.

### 2.6 Interpulse interval and call-rate identity

Substituting Equation 25 into Equation 12 gives

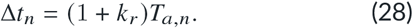

Using the call-duration-aware acquisition interval in Equation 19,

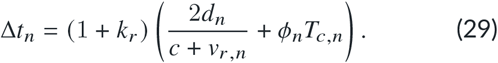

The call rate, *C*_*r,n*_, is the reciprocal of the interpulse interval,

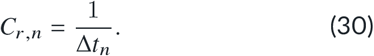

Combining Equations 28 and 30 yields the general callrate identity,

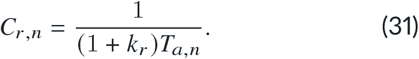

Using Equation 19,

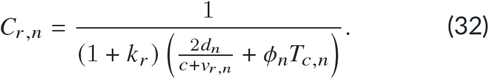

The same proportional structure also allows the acoustic and behavioural components of the IPI to be recovered directly from observed timing once *k*_*r*_ is known or estimated. Rearranging Equation 28 gives

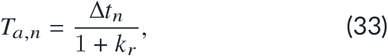

and substituting Equation 33 into Equation 25 gives

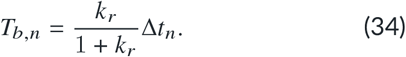

Using Equation 30, these can be written equivalently in call-rate form as

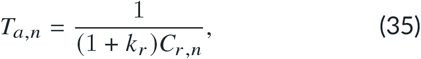

and

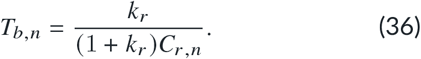

These inverse forms do not replace the motion-aware definition in Equation 19. Rather, they provide the complementary timing identities that partition any observed IPI into its implied acoustic and behavioural components under a specified responsivity coefficient.

When the information-window term is negligible, Equation 32 reduces to the first-arrival form

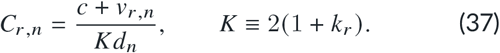

Equation 37 jointly constrains call rate, effective anchor distance, radial closing velocity, and the responsivity coefficient at each call event under first-arrival anchoring. In the far-field limit, where *v*_*r*_ ≪ *c*,

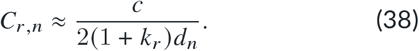

The far-field approximation also clarifies the direct role of closing velocity. At fixed distance and responsivity coefficient, the first-arrival solution gives

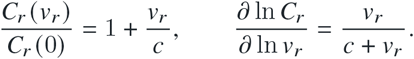

Thus, call rate is not directly proportional to *v*_*r*_ : its fractional sensitivity to velocity is only of order *v*_*r*_ /*c*. For radial closing velocities of 2–10 m s^−1^, the direct increase relative to the stationary-anchor value is approximately 0.6–2.9%. A non-zero selected echo window in Equation 32 reduces this sensitivity further because part of *T*_*a*_ is then independent of propagation delay. Closing velocity nevertheless remains important dynamically because it determines how rapidly distance changes and, therefore, how rapidly an approach traverses the distance-dependent call-rate trajectory.

The predicted inverse-distance relationship between call rate and distance therefore follows from the interval decomposition together with Equation 25 and the assumed distance dependence of acoustic acquisition. The hyperbola is not imposed as a curve-fitting function; it is the observable form predicted by these timing assumptions. In the more general model, the same inverse-distance form is preserved whenever the effective acquisition interval in Equation 19 scales approximately with distance and *k*_*r*_ is approximately constant over the analysed segment. The analytical consequences of these timing rules for call-rate scaling and call-duration contraction are summarised in Figure 2.

**Figure 2.**
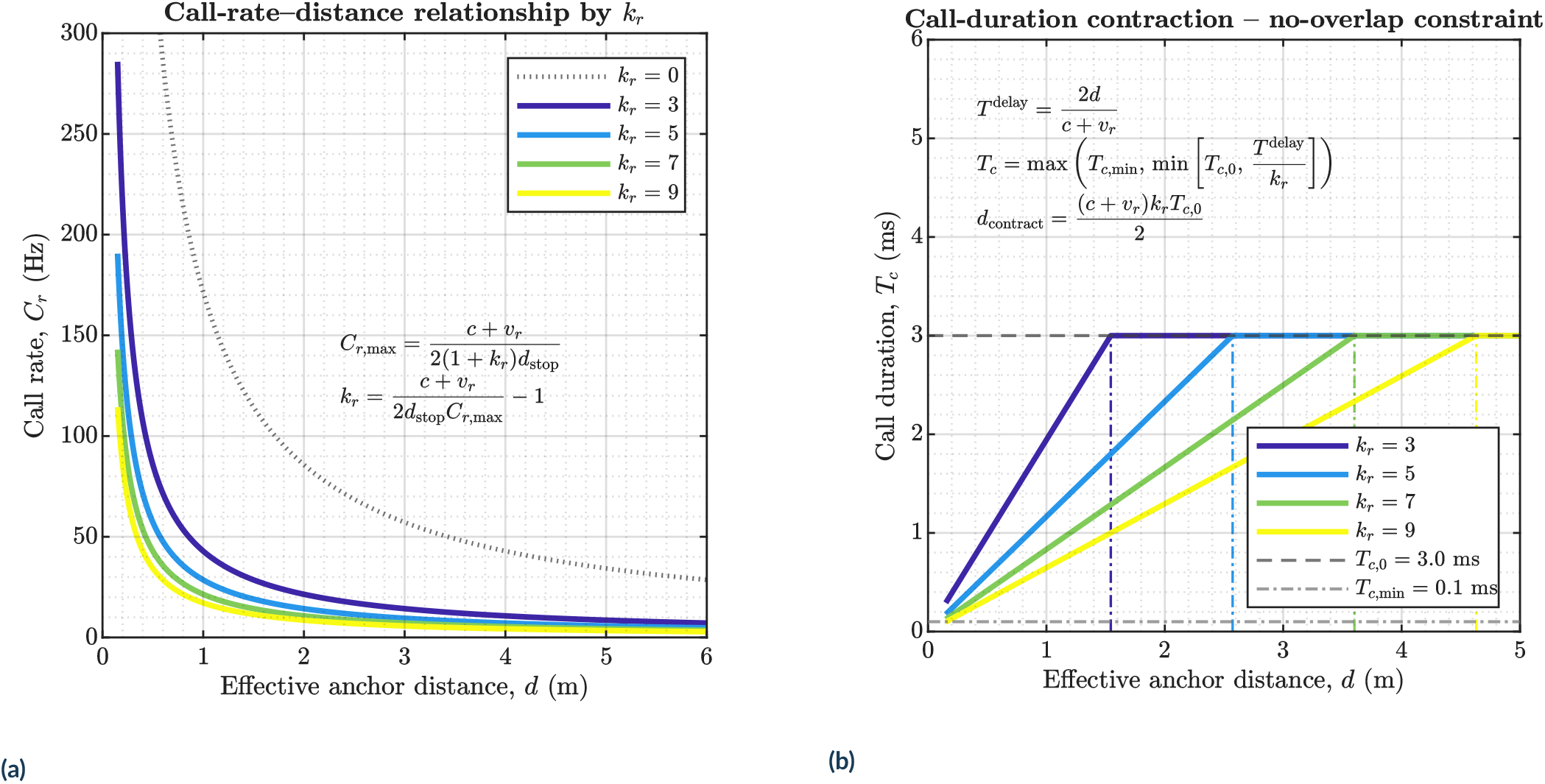
Analytical consequences of the responsivity framework for call rate and call duration. **a**, Under first-arrival anchoring, Equation 37 generates a family of hyperbolic call-rate trajectories across distance. Lower responsivity coefficients allow higher call-rate regimes at a given effective anchor distance, while higher responsivity coefficients shift the trajectory downward and lower the terminal ceiling. The accompanying rearrangement shows how *k*_*r*_ can be inferred from a specified stop distance, radial closing velocity, and terminal call rate. **b**, The no-overlap call-duration rule links emitted duration to the first relevant echo delay, such that call duration remains at its initial value until the available delay becomes too short, then contracts linearly with decreasing distance until the minimum permitted duration is reached. Together, the panels show that the framework constrains both the tempo of sensory updates and the signal duration through which those updates are acquired.

### 2.7 State identities and trajectory dynamics

The general identity in Equation 31 constrains call rate through the effective acquisition interval. When the anchor is approximated by first-arrival delay, Equation 37 can be rearranged to express either effective anchor distance or radial closing velocity in terms of the remaining variables. Solving for distance gives

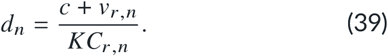

Solving for radial closing velocity gives

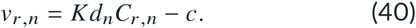

These identities are instantaneous algebraic constraints under first-arrival anchoring. They do not describe how distance or velocity evolves through time; rather, they specify the combinations of distance, velocity, and call rate that are mutually compatible under that reduced form of the responsivity model. If *k*_*r*_ is known or estimated, call rate can be used to infer the effective echo-generating distance when relative velocity is known. Conversely, if distance is known, call timing can be used to infer radial closing velocity.

If the information-window term in Equation 19 is not negligible, distance estimates must be corrected by the anchoring rule. In that case, the first step is to infer *T*_*a*_ from Equation 31, then map the inferred acquisition interval to distance using Equation 19.

To describe how call rate changes during an approach, we differentiate Equation 37 with respect to time. Since *d* = −*v*_*r*_ for positive closing velocity,

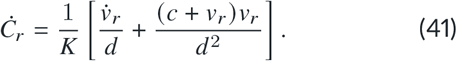

Rearranging for the dynamics of closing velocity gives

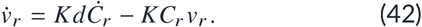

Under an approximately constant-velocity approach,

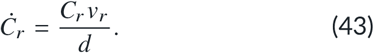

Combining this with the far-field distance identity yields a timing-based velocity estimator,

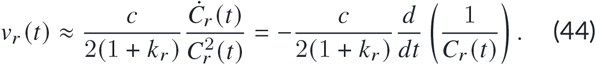

This result shows that relative velocity is encoded not simply in instantaneous call rate, but in the rate at which call rate changes during approach. Pooled call-rate–velocity data should therefore be interpreted as an accessible region generated by many approach trajectories rather than as a single within-sequence regression.

### 2.8 Maximum call rate and terminal update budget

Within the responsivity framework, the terminal buzz is interpreted as the high-call-rate regime that emerges when decreasing target distance drives the acoustic acquisition interval and the behavioural interval to very small values. Because the IPI is given by Equation 12, increasing call rate necessarily reflects the shrinkage of both *T*_*a*_ and *T*_*b*_, with the latter set by the responsivity coefficient *k*_*r*_ through Equation 25. The vocal-motor system therefore does not require a separate buzz-specific timing rule in the model; rather, the buzz emerges as the near-target limit of the same closed-loop responsivity law that governs the rest of the approach.

Under first-arrival anchoring, Equations 37 and 27 already show that call rate rises hyperbolically and that the behavioural interval shrinks with decreasing anchor distance. If the terminal limit is defined at a stop or interception distance *d*_stop_, the maximum call rate implied by the responsivity law is

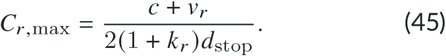

Equation 45 makes explicit that *k*_*r*_ sets the scale of the terminal ceiling: lower *k*_*r*_ permits a higher maximum call rate, whereas higher *k*_*r*_ lowers the ceiling. At the same stop distance, Equations 16, 27, and 28 define the corresponding minimum acoustic acquisition interval, minimum behavioural interval, and minimum IPI. Together, these relations show how the terminal buzz emerges mathematically. As the bat enters the final approach to the target, shrinking anchor distance drives call rate upward. The responsivity coefficient *k*_*r*_ determines how small the behavioural interval can become relative to the acoustic interval, and thus determines both the maximum achievable call rate and the density of terminal sensory updates.

To quantify the terminal high-rate regime, we define the final update window as the interval between a chosen fraction *f* of the maximum call rate and the stop distance. Setting *C*_*r*_ (*d*) = *f C*_*r*,max_ and substituting Equation 45 gives

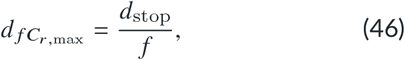

so that the corresponding final spatial interval is

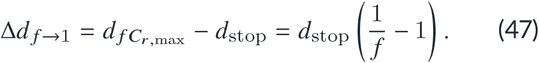

For a terminal window defined from 0.75*C*_*r*,max_ to *C*_*r*,max_, Equation 47 reduces to a short near-target interval that is set primarily by stop-distance geometry. Variation in buzz length then arises not because this terminal spatial interval changes strongly, but because radial closing velocity determines how long the bat remains within it. A first-order estimate of the time available inside the final interval is

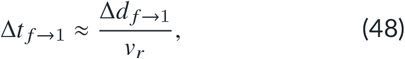

and the corresponding number of calls that fit into this final interval is

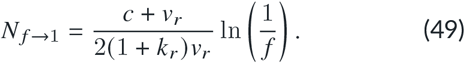

Equation 48 shows that radial closing velocity is the principal kinematic determinant of terminal buzz duration: slower terminal approaches prolong the time available in the final near-target regime, whereas faster approaches compress it. Equation 49 shows that *k*_*r*_ sets the density of updates within that same interval. Thus, *k*_*r*_ primarily determines the structure of terminal call timing, whereas *v*_*r*_ determines how long the bat remains in the regime where those short intervals can occur.

The terminal buzz can therefore be viewed as a finite update budget generated by the interaction between stop-distance geometry, responsivity coefficient, and radial closing velocity. Once the ceiling implied by Equation 45 is reached, further shortening of IPI is unavailable; remaining behavioural adjustment must occur through velocity, trajectory, capture manoeuvre, sensory selection, or anchor switching rather than further call-rate acceleration.

The constant-velocity estimate in Equation 49 provides a lower-level timing constraint, but real approaches may include deceleration. Simulated approach profiles illustrate how deceleration modifies the terminal update budget. The comparison shows that buzz-length variation can arise from the interaction between initial approach velocity and late-stage deceleration: high initial velocity compresses the available update window, whereas deceleration can recover additional near-target time and thereby extend the number of terminal calls.

### 2.9 Call duration and echo-overlap constraints

Call duration changes systematically during target approach. The hard pulse–echo overlap condition is set by the delay to the first relevant echo, not by the full acquisition interval. Overlap begins when the echo-start delay in Equation 16 becomes shorter than the emitted call duration. Thus, the no-overlap condition is

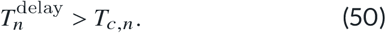

Solving by equality gives the overlap boundary distance

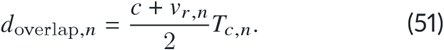

In the far-field limit, Equation 51 becomes

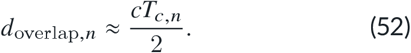

Call shortening should therefore begin before pulse– echo overlap compromises the behaviourally relevant echo stream. Because call duration also controls the spatial depth of the emitted signal through Equation 20, call shortening may reduce both overlap risk and perceptual load. The two constraints are related but not identical: pulse–echo overlap concerns whether outgoing and returning sounds physically overlap, whereas *T*_*ϕ,n*_ concerns how much of the call-defined echo stream contributes to the behavioural anchor.

### 2.10 Responsivity curve and buzz readiness

The coefficient *k*_*r*_ is the central responsivity parameter of the framework. We additionally define a derived timing-adjustment metric to quantify how sharply the IPI changes from one call interval to the next. For successive interpulse intervals,

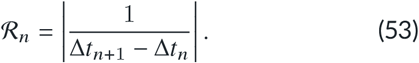

Large values of ℛ_*n*_ occur when successive IPI changes are small, indicating fine temporal adjustment near the terminal high-gain region of an approach. To compare this quantity with the physiological call-rate ceiling, we normalise it as

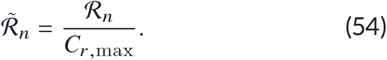

We define the buzz-readiness index, *n*^∗^, as the call index at which this timing-adjustment metric approaches the call-rate ceiling most closely,

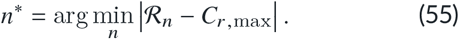

The associated IPI step is

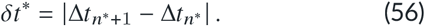

These quantities are used as trajectory diagnostics rather than as the primary definition of responsivity. The primary theoretical parameter remains *k*_*r*_ ; ℛ_*n*_ and *δt*^∗^ describe the realised sharpness of timing adjustment in a simulated or observed sequence. Under a single, stable acoustic anchor and orderly closing dynamics, the resulting responsivity curve is expected to follow a smooth, approximately hyperbolic trajectory. This single-anchor case provides the diagnostic baseline: departures from smoothness are not treated as model failure, but as potentially informative signatures of anchor switching, abrupt changes in relative motion, clutter interactions, or other transient instabilities in the sensorimotor loop.

### 2.11 Simulation framework

To evaluate how the foregoing derivations translate into observable timing structure, we implemented a custom call-by-call responsivity simulator in MATLAB. The simulator was designed to instantiate the event model across complete synthetic approach trajectories, beginning from an initial detection distance and ending either in interception or in one of several predefined stopping states. Beyond illustrating the analytical equations, its purpose was to encompass a range of ecologically plausible behavioural settings in which bats may regulate echolocation timing during prey pursuit, target approach, and near-terminal interception.

For each simulated sequence, the initial bat position, target position, initial bat speed, initial call duration, speed of sound, responsivity coefficient, stop or interception distance, maximum anchor distance, maximum sequence duration, and maximum number of calls were specified. The simulator can operate in either one-dimensional or three-dimensional geometry. In one-dimensional simulations, bat and target positions lie along a single approach axis. In three-dimensional simulations, positions and velocities are represented as vectors, and radial closing velocity is computed from the instantaneous bat–anchor line of sight using Equation 5. For the acoustic timing calculation, we defined a timing-constrained radial velocity,

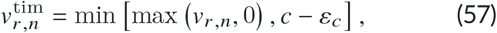

where ε_*c*_ > 0 is a small numerical margin that keeps the timing velocity below the speed of sound. Receding or stationary configurations therefore have 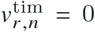 and were evaluated using the stationary-delay limit, whereas positive closing velocities entered the motion-aware delay directly. In the simulation, 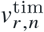 was substituted for *v*_*r,n*_ in Equation 16; the signed *v*_*r,n*_ was retained separately to describe the actual relative motion.

The simulator supports one or more possible targets. For two-target or multi-target simulations, the anchor can be selected as the nearest target, as a random target, as a fixed designated target, or by a hysteretic rule that preserves the previous anchor unless a switch occurs. These options allow the same event-level timing model to represent strict prey locking, nearest-object anchoring, intermittent environmental sampling, or temporary persistence on one object. Target motion can be disabled, held constant, or sampled stochastically. In stochastic motion mode, target speed is drawn around a specified fraction of bat speed and target direction is sampled in one or three dimensions according to the selected geometry. Bat speed can likewise be fixed, decelerating, or jittered around the specified mean approach speed, permitting the same simulator to represent steady approaches, variable-velocity pursuits, and terminal slowing before interception.

At each call event, the simulator first selects the acoustic anchor and computes anchor distance and radial closing velocity. The first relevant echo delay is then computed either with the stationary approximation in Equation 15 or with the motion-aware form in Equation 16. Unless otherwise stated, simulations used the motion-aware form. The information-window term *T*_*ϕ,n*_ was set as in Equation 17. The parameter ϕ_*n*_ can be held constant, prescribed by a sequence, or supplied by a user-defined function. The first-arrival special case is obtained when ϕ_*n*_ = 0. The acoustic acquisition interval is then obtained from Equation 18, and the behavioural interval is computed from Equation 25, optionally subject to a minimum response floor,

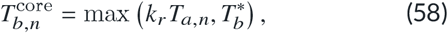

where 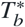 denotes an irreducible behavioural interval when this constraint is enabled.

The emitted IPI was generated as

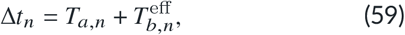

where 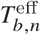 is the realised post-acquisition interval. In deterministic simulations, 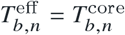. When caller-side timing variability was enabled, an additional non-negative lag term was added to the behavioural interval,

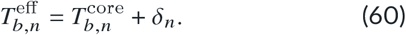

The optional lag δ_*n*_ was implemented as a bounded autoregressive process. Its mean was specified as a fraction of 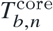,

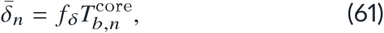

with stochastic deviations

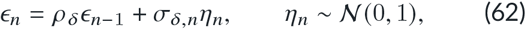

and

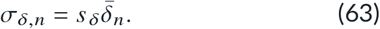

Here, *f* _δ_ is the mean-lag fraction, ϵ_*n*_ is the autoregressive deviation from that mean, ρ _δ_ is its lag-one autocorrelation coefficient, and η_*n*_ is an independent standard-normal innovation. The quantity σ_δ,*n*_ is the call-specific innovation standard deviation, and *s* _δ_ sets its magnitude as a fraction of the mean lag. The realised lag was

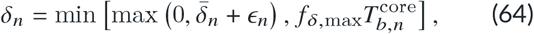

where *f* _δ,max_ is the maximum permitted lag as a fraction of the core behavioural interval. Thus, the realised lag remained non-negative and bounded above. This term represents caller-side motor or scheduling variability, not uncertainty in echo arrival.

The locally predicted displacement between calls was computed as

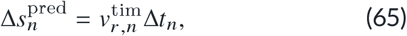

for diagnostic comparison with the local approximation in Equation 14. This timing-based prediction is distinct from the realised geometric displacement, which can retain receding motion. In vector simulations, the actual next-state geometry was obtained by updating bat and target positions over the same onset-to-onset interval,

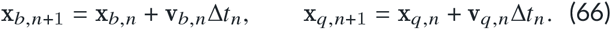

The next anchor distance was then recomputed from the updated positions. Simulations continued until the bat reached the specified interception distance, exceeded the maximum elapsed time, exceeded the maximum number of calls, or the selected anchor moved beyond a specified maximum anchor distance.

#### 2.11.1 Simulation scenarios and behavioural conditions

The same simulation core was used for the complementary analyses summarised in Table 1. Single-target trajectories and parameter sweeps traced the development of interval structure and terminal organisation, whereas multi-target simulations tested whether transient call groupings could emerge from anchor allocation without a grouping-specific timing rule. These are systematic configurations of one event model, not independent models.

**Table 1.**
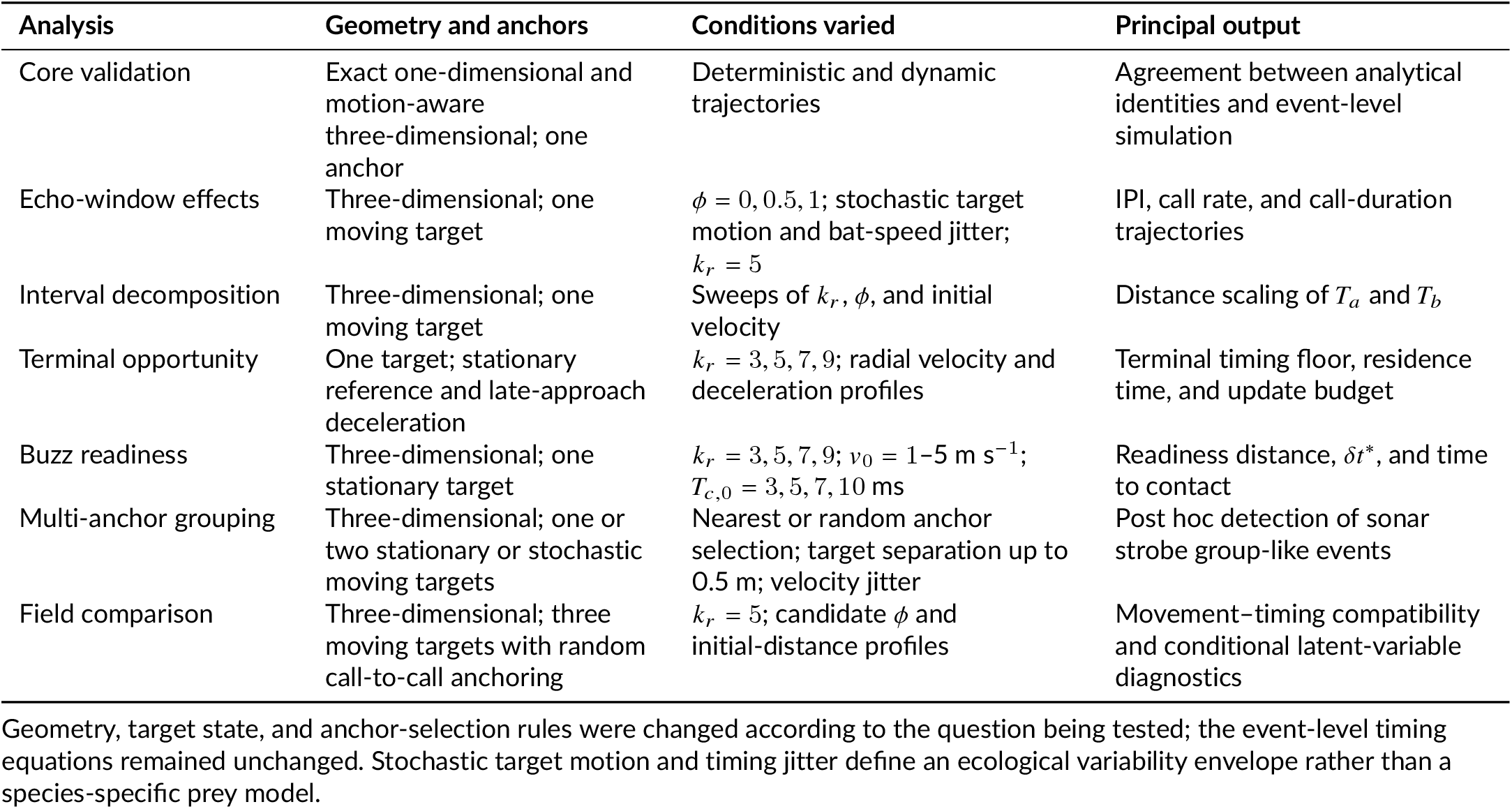
Simulation configurations and their analytical roles.

#### 2.11.2 Call duration and call-duration jitter

Call duration was treated as a state variable that must be determined before computing *T*_*ϕ,n*_. The simulator includes several call-duration modes. In the constant mode, *T*_*c,n*_ = *T*_*c*,0_, where *T*_*c*,0_ is the specified initial and maximum call duration. In the causal decrement mode, call duration for call *n* is based on the acoustic acquisition interval available from the previous call cycle,

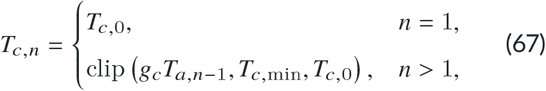

where *g*_*c*_ is the duration gain, *T*_*c*,min_ is the imposed minimum call duration, and clip (*x, a, b*) = min [max (*x, a*), *b*] constrains *x* to the interval [*a, b*]. This form avoids using future echo information from the call whose duration is being set. A threshold mode was also implemented to reproduce pulse–echo-overlap contraction rules, in which call shortening begins once the current first-echo delay approaches the emitted call duration, consistent with Equation 50.

To represent biological variability in duration control, an optional jitter term was added after the deterministic duration was computed. Additive, proportional, and combined jitter modes were implemented. In the additive mode,

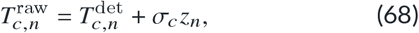

where 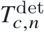 is the deterministic duration, σ_*c*_ is the additive duration-jitter standard deviation, and *z*_*n*_ is a standardised jitter state. In the proportional mode,

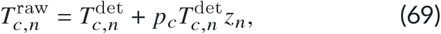

where *p*_*c*_ is the proportional jitter magnitude. In the combined mode,

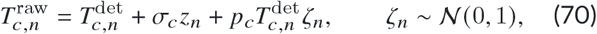

where ζ_*n*_ is an independent standard-normal variate. Correlated duration variability in the additive component was generated with an optional autoregressive state,

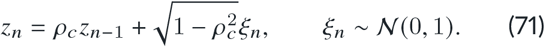

Here, ρ_*c*_ is the lag-one autocorrelation coefficient and ξ_*n*_ is an independent standard-normal innovation. The factor 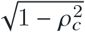 preserves unit marginal variance in *z*_*n*_; when autocorrelation was disabled, *z*_*n*_ was sampled independently from *N* (0, 1). The realised duration was clipped between *T*_*c*,min_ and *T*_*c*,0_. The output table stores both the deterministic and realised duration, allowing downstream analyses to separate the intended contraction rule from biological perturbation.

Optional wingbeat-state and waveform metadata could also be stored for subsequent analyses, but these variables did not enter the event-level timing calculations or the analyses reported here.

#### 2.11.3 Simulation output and downstream analysis

The simulator returned a long-format call table containing event times, anchor and geometric states, bat and target kinematics, acoustic and behavioural interval components, realised lag, call timing and duration, and displacement diagnostics. Parameter snapshots and random-number seeds were retained to support reproducible simulation sweeps.

All summary quantities were derived from these event-level outputs rather than imposed as additional simulation rules. These included time to contact, stop-distance-defined call-rate limits, terminal update budget, call-duration contraction, event-order relationships, and the trajectories of *T*_*a*_ and *T*_*b*_. For the multi-target simulations, candidate sonar strobe groups were identified subsequently from the emitted IPI sequence using within-group stability and flanking-interval criteria. Call grouping was therefore analysed as an emergent property of simulated anchor-allocation behaviour rather than encoded directly in the event model.

### 2.12 Field recordings, empirical processing, and model comparison

#### 2.12.1 Field recording setup

To evaluate the fit of the responsivity framework with natural bat behaviour, we analysed field recordings collected with a four-channel microphone array in a free-foraging setting at Weihenstephan, Freising, Germany, during August–September 2021. The array was designed for portable deployment in open field conditions and consisted of four custom-built microphones mounted on three wooden arms joined by a triangular central junction (Figure 3). The junction was mounted on a ball joint to allow angular adjustment of the array when required, and all exposed surfaces were covered with absorptive foam to reduce local reflections.

**Figure 3.**
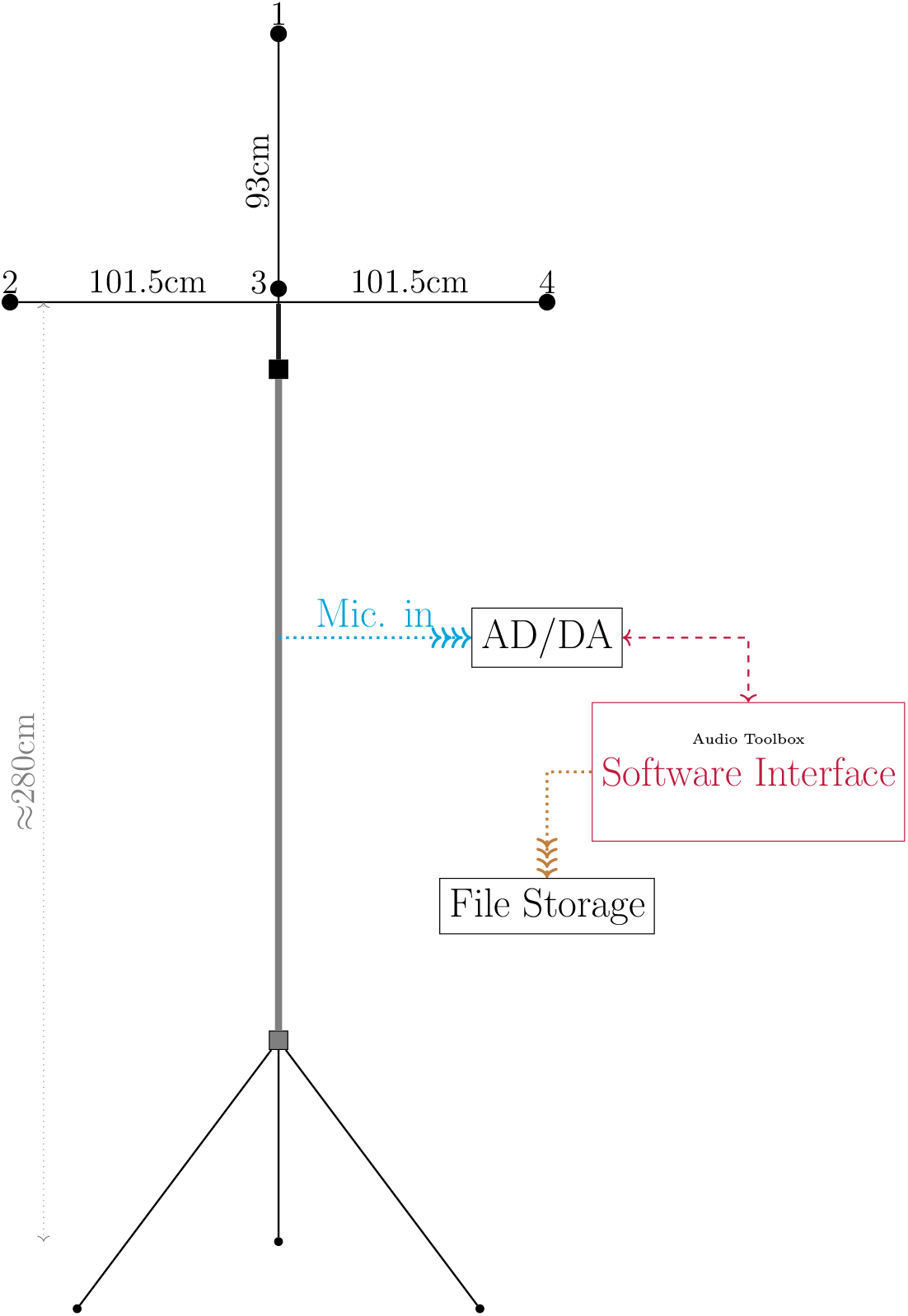
Four-channel field recording array used for empirical validation. The microphone array was designed for portable deployment in open foraging habitat and for reconstructing call-by-call flight trajectories from time-difference-of-arrival measurements. AD, analogue-to-digital; DA, digital-to-analogue.

The array was mounted on a studio tripod with sufficient ground clearance to record bats flying through the foraging space. The microphones were based on Knowles SPU0410LR5H-QB elements and were calibrated against a 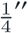 Bruel & Kjaer reference microphone. During acquisition, the incoming signal was convolved with a calibration filter in the frequency domain before being written as uncompressed .wav files. The recording system was powered in the field by 12 V car batteries coupled to a direct-current to alternating-current (DC–AC) inverter supplying the computer and audio interface. Trigger-based recording was used, with automatic extension when call activity occurred near the end of the nominal recording window.

#### 2.12.2 Trajectory reconstruction and call-level extraction

Call localisation and trajectory reconstruction were performed in MATLAB using time-difference-of-arrival estimation as described in our earlier methodological work on array localisation and spatial tracking (Umadi, 2025a,b). The processed dataset consisted of 6360 call rows distributed across 496 sequences. For each call, the exported table stored the call timestamp, interpulse interval, call rate, emitted call duration, call level, localisation-derived position, displacement between successive localised call positions, and velocity estimated from displacement divided by the corresponding timestamp difference.

Only call sequences with adequate signal-to-noise ratio were retained for downstream analysis, and extracted calls were manually checked to remove obvious ground reflections and low-quality detections. Because localisation uncertainty increases at longer range and in weak-signal segments, the raw processed table was further filtered before model comparison.

We therefore applied a two-stage quality-control procedure to the processed call table. First, rows were flagged if they violated basic physical constraints, including non-positive call rate, non-positive IPI, negative velocity, non-positive call duration, or undefined timestamp. Second, robust within-sequence filtering was applied using median absolute deviation (MAD) criteria to call rate, IPI, velocity, call duration, and call-to-call duration difference, together with MAD-based jump filtering on the same variables to identify abrupt discontinuities inconsistent with smooth within-sequence behaviour. Rows exceeding a conservative velocity cap of 10 m s^−1^ were also excluded from the model-comparison subset. Under this combined procedure, 1453 of 6360 rows (22.85%) were flagged, and only the retained subset was used for empirical comparisons with the model.

#### 2.12.3 Inter-call displacement as the shared empirical coordinate

Since the field data corresponded to freely foraing bats, direct bat–target distance was not available in the field recordings. The localisation pipeline reconstructs bat position relative to the array, not the absolute position of the prey item being pursued. Therefore, we used the displacement between successive calls as the shared kinematic coordinate for empirical–simulation comparison.

This choice follows the logic of the responsivity framework itself. The model is fundamentally concerned with how much movement occurs between successive sensory updates. In the field data, inter-call displacement is the directly observable outcome of that update cycle. In the simulations, the same quantity can be derived from the realised bat motion between successive call onsets. Intercall displacement therefore provides the most defensible common spatial variable across empirical and synthetic datasets, even though it is not equivalent to the remaining bat–target distance. The empirical validation was thus performed in displacement space, asking whether observed call timing occupied the same movement–timing relationships predicted by the responsivity framework.

#### 2.12.4 Matched field–simulation comparison and conditional parameter diagnostics

The primary field–simulation comparison used only quantities available in both datasets without assuming a value for an unobserved anchor: IPI, call rate, inter-call displacement, velocity, and emitted call duration. The retained field rows were additionally analysed with a fixed responsivity coefficient of *k*_*r*_ = 5 for conditional framework diagnostics. Under this assumption, observed IPI can be partitioned into inferred acquisition and behavioural components using Equations 28, 33, and 34. These inferred quantities are deterministic transformations of IPI at the specified *k*_*r*_, rather than independent field measurements, and were used only to test the consequences of candidate echo-window and anchor assumptions.

The field-validation simulations used the same simulator core described above, but were configured in a three-dimensional, motion-aware mode with three available targets and random anchor selection. This setting was used not to reproduce any one observed pursuit geometry literally, but to generate a broad behavioural envelope against which the field observables could be compared in shared displacement space. Target motion was stochastic, with target speed scaled to a fraction of bat speed and perturbed by modest jitter. Bat speed was also jittered from call to call, and the initial speed of each synthetic sequence was sampled across the retained field velocity range up to 10 m s^−1^. Simulations were generated across ϕ = 0, 0.5, and 1, with 100 sequences per ϕ value.

For the present comparison, the available simulation speed range was extended across the retained field range and simulated calls were sampled within 0.5 m s^−1^ velocity bins to reproduce the call-level field velocity distribution up to 10 m s^−1^. This matching prevents a different mixture of slow and fast calls from being mistaken for a difference in timing control. Call timing followed the motion-aware delay model, and call duration was updated causally from the previous call’s acquisition interval (using the previousTa mode, see code), with additional additive duration jitter and weak temporal autocorrelation. A maximum call-rate ceiling was enforced for each sequence using the stop-distance-based derivation defined in the framework, so that the simulated terminal regime remained consistent with the same closed-loop timing constraints used elsewhere in the study. For each simulated call, inter-call displacement was calculated from the realised onset-to-onset bat motion, and only rows with finite values and bat speeds ≤10 m s^−1^ were retained. We compared call rate and IPI against inter-call displacement, IPI within shared velocity bins, and call duration against call rate. Binned medians and interquartile ranges were used to display the central relationships while retaining the spread of each dataset.

Compatibility with the distance-dependent call-duration envelope was evaluated across 20 independently seeded simulation ensembles. Each ensemble contained the same parameter grid and number of sequences, but used a predetermined seed for initial speeds and stochastic trajectory and duration variation. For each ensemble, the simulated 5–95% duration band was calculated across anchor distance and the percentage of field calls falling within that band was recorded. We report the median and interquartile range of this percentage across ensembles; Figure 8aE displays the ensemble whose compatibility percentage was closest to the across-seed median.

We then asked whether the residual timing difference after velocity matching had the direction expected from a change in the selected acoustic acquisition window. Holding *k*_*r*_ = 5 and the default initial simulation distance *d*_0_ = 5 m fixed, we profiled candidate echo-window fractions from ϕ = 0 to 2 in increments of 0.05, generating 100 sequences per candidate value. To provide common kinematic support across the full profile, field and simulated calls were sampled within 1 m s^−1^ bins over 2–9 m s^−1^, yielding 894 matched calls for every value of ϕ. At each candidate value, the field IPI was converted conditionally into *T*_*a*_, *T*_*b*_, and inferred anchor distance using the motion-aware identities, while the corresponding simulated quantities were taken from the known event state. Combining Equations 33 and 19, the inferred propagation delay was *T*_*a*_ −*ϕT*_*c*_ = Δ*t* / (1 +*k*_*r*_) −*ϕT*_*c*_. Because a negative propagation delay is physically inadmissible, this term was constrained to be non-negative before conversion to distance: 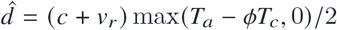. Thus, calls for which *T*_*a*_−ϕ*T*_*c*_ ≤0 were assigned an inferred distance of zero. Agreement was quantified as the mean absolute difference between corresponding percentiles from the 1st to the 99th percentile. The distance and *T*_*b*_ profiles were therefore not independent estimates of ϕ; they were complementary diagnostics of how the assumed acquisition window altered two framework-derived distributions.

Finally, we repeated the profile over initial distances *d*_0_ = 2–8 m and *ϕ* = 0–2 to test whether the selected default geometry alone generated the inferred optimum. This sensitivity analysis used common random-number schedules and common velocity support across parameter combinations. The resulting profile was used as a robustness check and was summarised numerically rather than treated as an additional empirical measurement.

### 2.13 Validation of analytical identities

To test internal consistency, we used the simulator output to reconstruct the quantities that generated each call interval. The validation asks whether the discrete event simulation preserves the algebraic relationships derived from the responsivity framework.

The validation was performed in the first-arrival special case, with *ϕ*_*n*_ = 0, no caller-side lag, no response floor, and no call-rate ceiling. Under these conditions, Equations 28 and 16 imply the first-arrival call-rate identity in Equation 37. We therefore reconstructed anchor distance using Equation 39 and radial closing velocity using Equation 40, where *K* = 2 (1 +*k*_*r*_). We also checked that the radial closing velocity stored for timing was recovered from the local step relation, 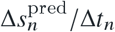, and compared this with the position-derived displacement across the same interval.

Three validation modes were implemented. The exact mode used a deterministic one-dimensional approach with fixed radial velocity and therefore provides a strict machine-precision check of the identities. A variable-velocity mode introduced bat-speed jitter to produce a changing *v*_*r*_ profile while preserving the local timing identities. A moving-target mode introduced stochastic target motion and tested whether the local anchor distance, radial closing velocity, and call rate remained mutually consistent when the target state changed from call to call. Hard numerical assertions were applied only to the exact deterministic mode; the variable-velocity and moving-target modes were used to visualise the same internal relationships under more dynamic trajectories.

Across all three validation modes, the recovered quantities remained effectively identical to the simulated state variables within machine precision (Figure 4). In the exact deterministic mode, the mean absolute difference between the stored radial closing-velocity profile and the value recovered from the predicted local step was exactly zero, the median absolute anchor-distance reconstruction error was of order 10^−18^ m, the median difference between step-recovered and identity-recovered velocity was exactly zero, and the median difference between observed and predicted stepwise velocity was of order 10^−14^ m s^−1^. In the variable-velocity mode, the corresponding discrepancies were of order 10^−17^ m s^−1^ for the profile-versus-step velocity comparison, 10^−17^ m for anchor distance, and 10^−14^ m s^−1^ for both velocity-identity comparisons. In the moving-target mode, these quantities remained of the same order: 10^−16^ m s^−1^ for the profile-versus-step velocity comparison, 10^−17^ m for anchor distance, and 10^−14^ m s^−1^ for the velocity-identity comparisons. Thus, in both the strict deterministic case and the more exploratory dynamic cases, the simulator preserved the analytical identities to essentially zero error, with residual differences attributable only to floating-point representation rather than to inconsistency in the event model.

**Figure 4.**
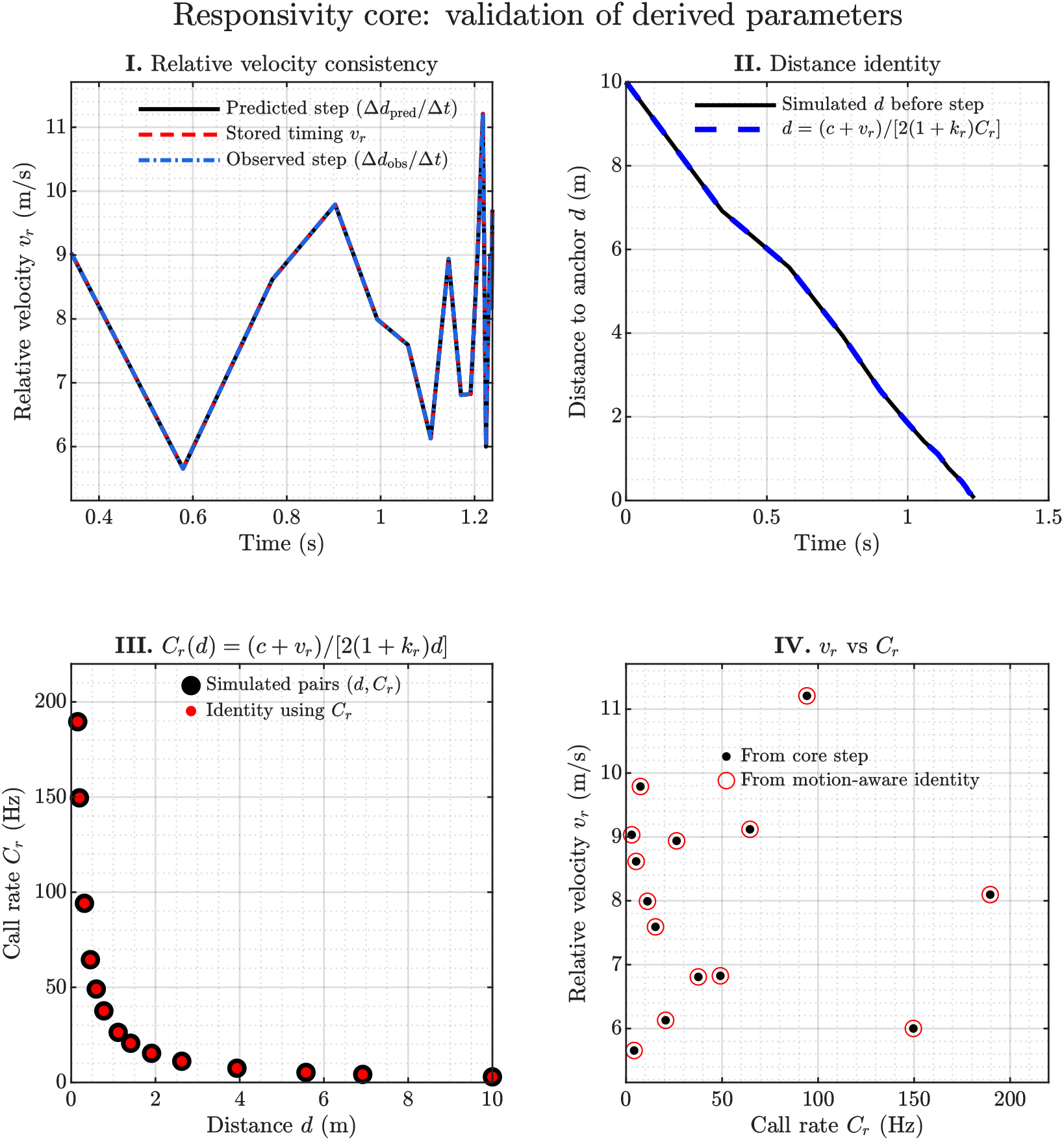
Validation of analytical identities in the responsivity framework. Simulated trajectories were used to validate the derived relationships among call rate, effective anchor distance, and radial closing velocity. The example shown uses the variable-velocity validation mode. The stored radial closing velocity matches the velocity recovered from the local displacement step, reconstructed anchor distances match the simulated distances, and velocities inferred from the analytical identities match those computed from the simulated timing variables. This confirms that the discrete event simulation preserves the timing constraints derived from the responsivity framework before the same simulator is used for parameter sweeps and downstream analyses.

The validated simulator outputs were then used throughout the study for parameter sweeps across *k*_*r*_, *ϕ*, call duration, and velocity, for empirical comparison in inter-call-displacement space, and for the analysis of emergent temporal patterns.

## 3 RESULTS

The responsivity framework advances a simple but consequential prediction: if call timing is governed by the coupled evolution of acoustic acquisition, behavioural interval, and relative motion, then a broad range of echolocation patterns should emerge from the same closed-loop rule rather than from separate behavioural programmes. The results that follow evaluate this prediction in progressively more complex settings. We first examine the analytically expected timing consequences of echo-window choice, responsivity, and approach kinematics, then test how these same variables shape terminal organisation, buzz structure, and buzz readiness, and finally ask whether the framework can also account for multi-target grouping patterns and the structure of empirical field recordings. Together, these analyses assess whether responsivity provides a unified description of how bats organise call timing during prey approach and interception.

### 3.1 Echo-window choice shifts distance-dependent timing under motile-target conditions

To isolate how echo-window choice reshapes call timing under pursuit-like variability, we analysed 4039 calls from three-dimensional simulations with one stochastically moving target, call-to-call bat-speed jitter, fixed *k*_*r*_ = 5, and *ϕ* = 0, 0.5, or 1 (Table 1). We fitted distance-by-*ϕ* interaction models for IPI and call duration, tested whether pairwise curve separation narrowed near the target, and examined whether *ϕ* altered call-rate spread within the 0.3–1.5 m bend region. Complete fitted coefficients and secondary tests are reported in Supplementary Table S5. IPI showed a strong and systematic dependence on echo-window choice (Figure 5). Distance, *ϕ*, and their interaction were all significant (*p* ≈0; *R*^2^ = 0.997), and fitted IPI slopes increased from 34.36 to 37.62 and 40.42 ms m^−1^ across *ϕ* = 0, 0.5, and 1. Increasing *ϕ* therefore shifted the IPI–distance relation upward and made it steeper. Pairwise separation was smallest near the target and increased with distance; the *ϕ* = 0 versus 1 difference increased at 6.14 ms m^−1^ (*p* ≈0, *R*^2^ = 0.840). Echo-window choice thus leaves a clear signature on onset-to-onset timing, but that signature compresses at short range.

**Figure 5.**
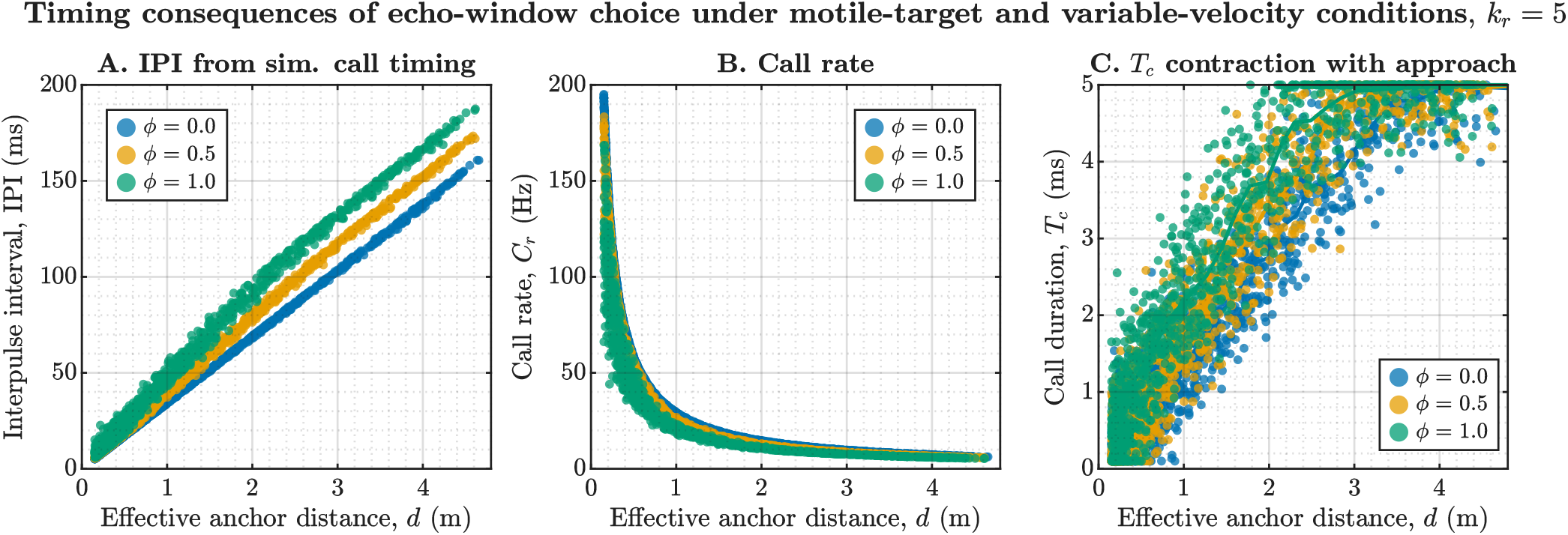
Echo-window choice reshapes simulated timing under motile-target and variable-velocity conditions. Simulations were run with one moving target, three-dimensional geometry, stochastic target motion, call-to-call velocity jitter, and fixed responsivity *k*_*r*_ = 5, while varying echo-window fraction *ϕ* across 0, 0.5, and 1. **A**, Increasing *ϕ* shifted the IPI–distance relation upward and made it steeper, indicating that a larger selected echo window lengthened the composite onset-to-onset timing interval at a given anchor distance. **B**, The corresponding call-rate curves were displaced downward, but the spread around the bend region was not strongly attributable to *ϕ* itself, instead reflecting stochastic motile-target and velocity-jitter dynamics shared across conditions. **C**, Call durations contracted with decreasing distance in all conditions, while *ϕ* mainly shifted the fitted duration trajectories upward rather than strongly altering their slopes. Together, the panels show that echo-window choice leaves its clearest signature on IPI, whereas call-duration contraction remains primarily governed by the imposed distance-dependent shortening rule.

Call duration shifted more weakly. Fitted slopes were similar across *ϕ* (1.09, 1.08, and 1.01 ms m^−1^), whereas intercepts rose from 0.33 to 0.55 and 0.94 ms; none of the pairwise convergence slopes was significant (*p* ≥0.054). Thus, *ϕ* displaced the expected duration envelope mainly as an offset while target-distance-dependent shortening remained dominant. The call-rate spread analysis likewise found no effect of *ϕ* or its interaction with distance in the bend region (*p* = 0.890 and *p* = 0.955). Its visible breadth therefore arose chiefly from the kinematic variability shared across conditions rather than from echo-window choice itself.

Taken together, these simulations show that *ϕ* has a strong and statistically robust effect on composite timing under the conditions examined, especially on IPI, while exerting a weaker and more offset-like influence on call duration. The strongest and most interpretable signature of echo-window choice is therefore the upward displacement and distance-dependent separation of the IPI curves. This section sets up the interval-decomposition results that follow, where the same *ϕ*-dependent timing shifts are resolved into their acoustic acquisition and behavioural components.

### 3.2 Distance dependence of acoustic acquisition and behavioural intervals

We next quantified how *T*_*a*_ and *T*_*b*_ scaled with effective anchor distance in separate sweeps of responsivity (*k*_*r*_ = 3, 5, 7, 9), echo-window fraction (*ϕ* = 0, 0.5, 1), and initial speed (*v*_0_ = 2, 4, 6, 8, 10 m s^−1^; Table 1). Per-condition regressions described interval–distance scaling, nested interaction models tested changes in level and slope, and binned pairwise differences assessed short-range convergence across *ϕ*. Complete coefficients and tests are given in Supplementary Table S6.

The *k*_*r*_ sweep confirmed that *T*_*a*_ was effectively independent of the responsivity coefficient in practical terms (Figure 6). Across *k*_*r*_ = 3, 5, 7, 9, its fitted slopes were nearly identical (5.99–6.05 ms m^−1^), despite formally significant effects in the large dataset. By contrast, *T*_*b*_ slopes increased from 17.96 to 30.23, 42.38, and 54.43 ms m^−1^. Responsivity therefore acted primarily on the behavioural interval and the achievable call-rate regime at a given distance.

**Figure 6.**
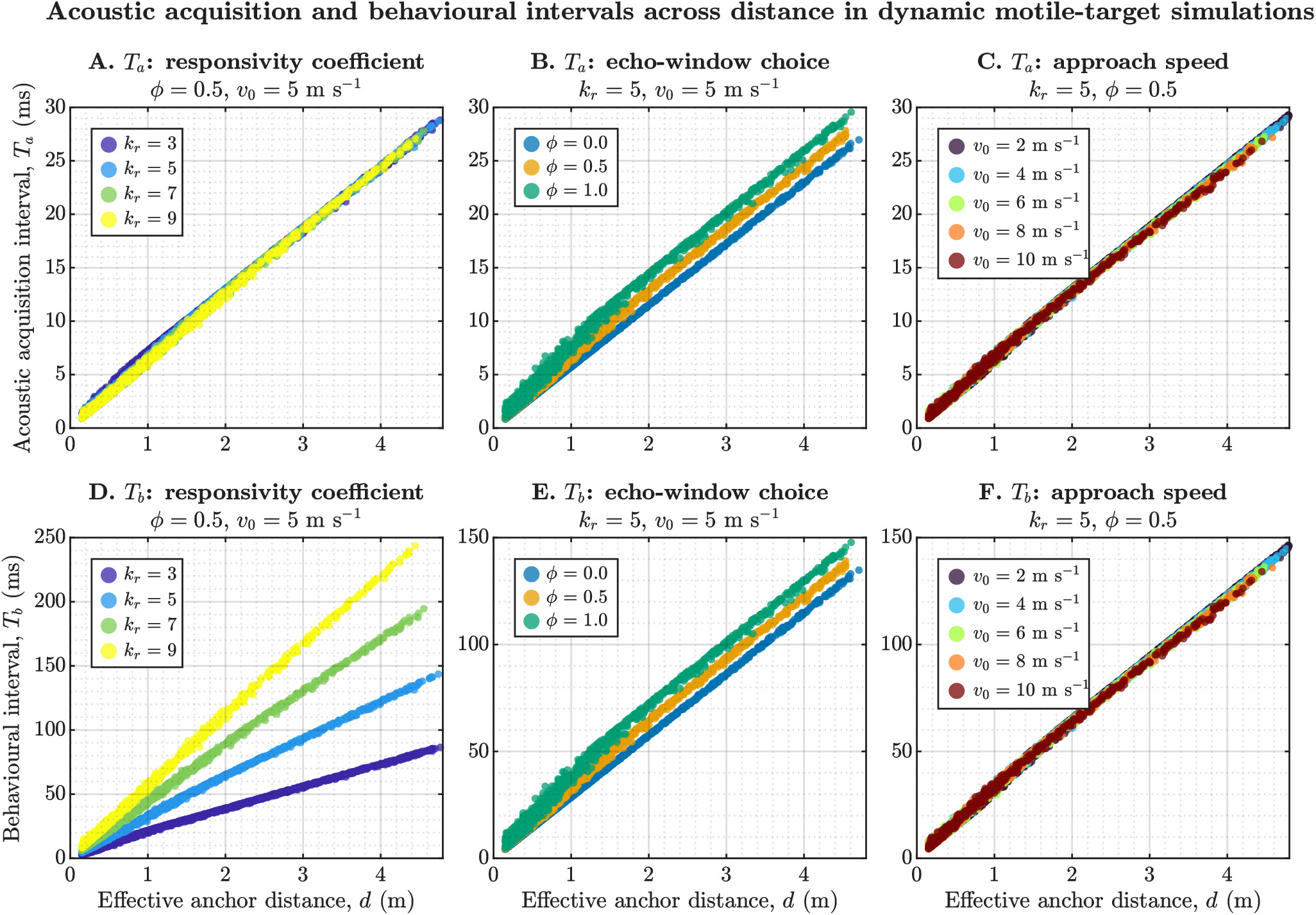
Distance dependence of acoustic acquisition and behavioural intervals in dynamic motile-target simulations. The upper row shows the acoustic acquisition interval, *T*_*a*_, as a function of effective anchor distance, and the lower row shows the behavioural interval, *T*_*b*_, for the same simulations. Panels A and D sweep responsivity coefficient (*k*_*r*_ = 3, 5, 7, 9) while holding *ϕ* = 0.5 and *v*_0_ = 5 m s^−1^ constant; panels B and E sweep echo-window fraction (*ϕ* = 0, 0.5, 1) at fixed *k*_*r*_ = 5 and *v*_0_ = 5 m s^−1^; panels C and F sweep initial approach speed (*v*_0_ = 2, 4, 6, 8, 10 m s^−1^) at fixed *k*_*r*_ = 5 and *ϕ* = 0.5. Quantitative comparisons were made using per-condition ordinary least-squares fits and nested linear models testing main effects of distance, condition, and their interaction. *T*_*a*_ remained nearly invariant across *k*_*r*_, whereas *T*_*b*_ changed strongly with *k*_*r*_, indicating that responsivity primarily reshapes the behavioural interval. Varying *ϕ* shifted both *T*_*a*_ and *T*_*b*_, but pairwise curve separation increased with distance and narrowed near the target, consistent with convergence of anchoring modes at short range. Velocity produced comparatively small changes in either interval over the plotted range.

The *ϕ* sweep shifted both intervals upward, but the curves converged towards shorter anchor distances. From *ϕ* = 0 to 1, fitted slopes increased from 5.74 to 6.29 ms m^−1^ for *T*_*a*_, and from 28.72 to 31.44 ms m^−1^ for *T*_*b*_. The separation between the *ϕ* = 0 and 1 curves increased with distance at 0.48 and 2.38 ms m^−1^, respectively (*p* < 0.001). Echo-window choice therefore had its strongest effect farther from the anchor, while near-target timing became increasingly compressed across conditions.

Approach speed had comparatively weak effects. Across *v*_0_ = 2–10 m s^−1^, fitted slopes changed only from 6.13 to 5.91 ms m^−1^ for *T*_*a*_, and from 30.64 to 29.54 ms m^−1^ for *T*_*b*_. This is expected from Equation 37: at a given distance, velocity enters the first-arrival relation only through the small correction 1+ *v*_*r*_ /*c*. Velocity therefore has little influence on instantaneous call rate at fixed distance, although it strongly determines how quickly an approach traverses the distance-dependent timing curve.

Taken together, these simulations reveal a clear functional separation within the responsivity framework. The acoustic acquisition interval is governed mainly by anchor geometry and echo-window definition, whereas the behavioural interval is where responsivity exerts its dominant effect. This result foreshadows the terminal-buzz analyses developed below, because it shows that the responsivity-dependent expansion or compression of *T*_*b*_ is the principal route by which the framework modulates the call-rate regime available near interception.

### 3.3 Terminal buzz length emerges from the interaction between responsivity and closing velocity

We next used the responsivity framework to ask why terminal buzzes can vary so widely in length, even when bats operate under a common closed-loop timing rule. The analysis was built in two steps. First, we calculated the terminal timing floor at *d*_stop_ = 0.15 m for *k*_*r*_ = 3, 5, 7, 9 using the stationary-anchor reference (*v*_*r*_ = 0) so that the implied maximum call rate could be defined for each responsivity condition. We selected 0.15 m as a representative late-interception boundary because it lies within the approximately 10–22 cm range at which *Myotis daubentonii* enters the final tail-down capture stage and agrees with the 15–20 cm distance at which *Noctilio albiventris* was reported to cease signal emission before prey capture (Kalko and Schnitzler, 1989; Kalko et al., 1998). European pipistrelles were reported to continue emitting closer to the target, stopping at 6.7± 2.0 cm (range 3.5–10 cm), showing that the selected boundary is a biologically plausible reference rather than a universal stopping distance (Kalko, 1995). Second, we analysed the final interval from 75% of *C*_*r*,max_ to *d*_stop_ across a radial closing-velocity sweep of 0.1–3.0 m s^−1^, and then added a set of illustrative late-approach deceleration profiles for *k*_*r*_ = 5 to show how time and call number accumulate when closing speed drops before interception.

The stationary-anchor reference revealed a clear responsivity-dependent ceiling (Figure 7A). At *d*_stop_, the minimum acoustic acquisition interval remained constant across conditions (*T*_*a*_ = 0.875 ms), but the shortest behavioural interval increased from 2.62 to 4.37, 6.12, and 7.87 ms for *k*_*r*_ = 3, 5, 7, 9, respectively. The corresponding minimum onset-to-onset intervals increased from 3.50 to 5.25, 7.00, and 8.75 ms, implying decreasing terminal call-rate ceilings of 285.83, 190.56, 142.92, and 114.33 Hz. This range brackets reported terminal regimes of approximately 155–210 Hz in *M. daubentonii*, 180–200 Hz in European pipistrelles, and 160–180 Hz in *N. albiventris* (Kalko, 1995; Kalko and Schnitzler, 1989; Kalko et al., 1998). The correspondence places the analytical ceilings within an observed biological range, but does not identify the responsivity coefficient underlying any particular field sequence. Thus, the responsivity coefficient determines the density of terminal updates that is theoretically available at the stopping distance.

**Figure 7.**
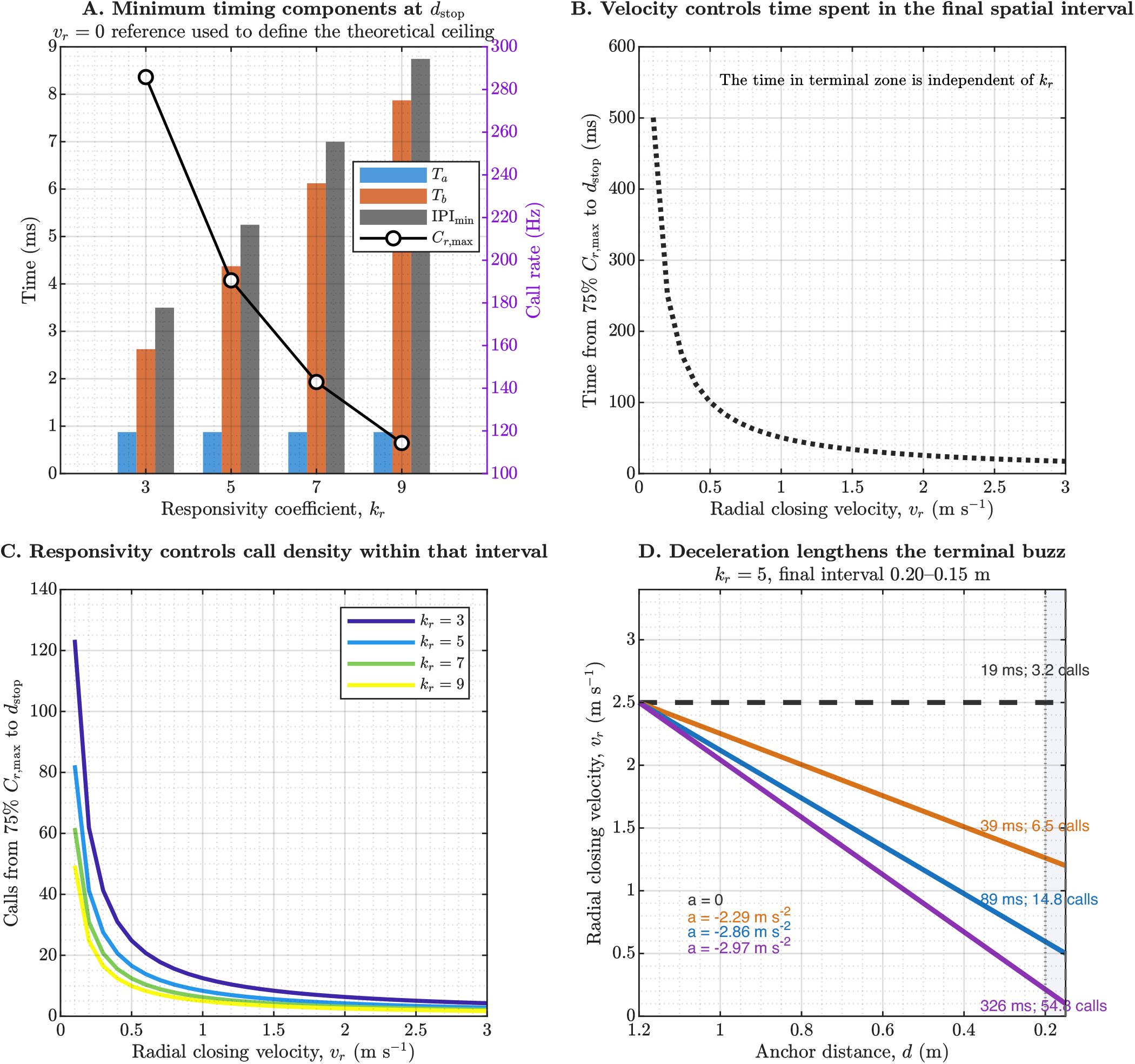
Responsivity principally sets terminal call density, whereas closing velocity sets terminal buzz duration. Panel A shows the stationary-anchor reference at *d*_stop_ = 0.15 m, used to define the theoretical terminal timing floor and the implied *C*_*r*,max_ for each responsivity condition. *T*_*a*_ remains fixed at the stopping distance, whereas *T*_*b*_ and the minimum onset-to-onset interval increase with *k*_*r*_, lowering the achievable terminal call-rate ceiling. Panel B shows that the time available between 75% of *C*_*r*,max_ and *d*_stop_ is identical across *k*_*r*_ and depends only on radial closing velocity. Panel C shows that, within that same spatial interval, lower *k*_*r*_ allows more calls to be emitted because the terminal call density is higher. Panel D illustrates the consequence for buzz length: under fixed *k*_*r*_ = 5, progressive late-approach deceleration lengthens the time spent in the 0.20–0.15 m terminal interval and thereby increases the number of calls emitted there. Variable terminal buzz lengths therefore emerge naturally from the interaction between a responsivity-defined call-density ceiling and kinematically determined time spent near the target.

By contrast, the duration of the final spatial interval was controlled entirely by radial closing velocity (Figure 7B). Because the interval from 75% of *C*_*r*,max_ to *d*_stop_ spans the same geometric region for all *k*_*r*_, the time available in that zone was identical across responsivity conditions and varied only with speed. Across the explored range, time spent in the terminal zone increased from 0.017 to 0.501 s as *v*_*r*_ decreased from 3.0 to 0.1 m s^−1^. This result is important because it shows that the overall duration of the buzz-like terminal phase is not set by responsivity alone.

Responsivity instead controlled how many calls could be packed into that available time (Figure 7C). Across the same 75%–100% terminal interval, the number of calls ranged from 1.71 to 123.50 depending on the combination of *k*_*r*_ and *v*_*r*_, with lower *k*_*r*_ yielding systematically denser terminal call sequences. The framework therefore separates two quantities that are often conflated behaviourally: closing velocity determines how long the bat remains in the terminal zone, whereas responsivity principally determines how densely calls can be emitted while the bat is in that zone. The small direct increase in call rate with *v*_*r*_ is minor relative to the inverse dependence of terminal residence time on *v*_*r*_ .

This separation becomes especially clear in the deceleration examples (Figure 7D). For *k*_*r*_ = 5, with the final interval defined as 0.20–0.15 m, a non-decelerating approach at 2.5 m s^−1^ yielded only 19 ms and 3.2 calls in the terminal zone. Slowing to 1.2 m s^−1^ increased this to 39 ms and 6.5 calls, slowing further to 0.5 m s^−1^ increased it to 89 ms and 14.8 calls, and a very strong deceleration to 0.1 m s^−1^ extended the same spatial interval to 326 ms and 54.8 calls. These simulated speeds overlap the late-pursuit kinematics reported in natural foraging: European pipistrelles slowed from approximately 4–7 to 1.5–3.5 m s^−1^ and sometimes reached 0.25–0.5 m s^−1^ during high pursuits, while aerially capturing *N. albiventris* slowed to 1–2 m s^−1^ and could become nearly stationary (Kalko, 1995; Kalko et al., 1998). The reported values are total flight speeds rather than reconstructed radial closing velocities, but demonstrate that pronounced late-pursuit deceleration occurs naturally. The interpretation is that variable buzz lengths arise not because the underlying responsivity rule changes, but because the time available inside the final spatial interval expands when radial closing velocity falls. Under the same *k*_*r*_, deceleration converts the same terminal geometry into a much longer and more call-rich buzz. In active foraging conditions, no two interception scenes are likely to be exactly alike: wind, prey type, manoeuvrability, flight behaviour, and the bat’s own approach dynamics all reshape relative motion near the target. When those conditions keep bat and prey in very close proximity under low or near-constant relative motion, the terminal buzz can lengthen substantially; conversely, relative motion that increases radial closing velocity can compress the available terminal interval and reduce the opportunity for a longer buzz.

### 3.4 Field timing shares the predicted movement structure and reveals conditionally interpretable differences

We next asked how the responsivity framework compared with call timing measured from free-flying bats. Because prey position was not available in the field recordings, the comparison could not begin from bat–target distance. We instead used inter-call displacement, the distance travelled by a bat between successive emissions, as a shared and directly observable coordinate. The field dataset comprised 6360 calls from 496 reconstructed sequences recorded from unidentified aerial-hawking bats belonging to the genera *Pipistrellus* and *Myotis*. After robust outlier flagging and a retained-row velocity cap of 10 m s^−1^, 4427 calls remained for IPI-based comparisons and 4399 for call-duration fits. We compared these observations with three-dimensional, motion-aware simulations containing three moving targets and random call-to-call anchor selection, generated at *k*_*r*_ = 5 across *ϕ* = 0, 0.5, and 1.

We first tested whether field and simulated calls occupied the same movement–timing regime. Call rate and inter-call displacement jointly imply an effective velocity, *v*_eff_ = *C*_*r*_ Δ*x*, so we sampled simulated calls within velocity bins to reproduce the field velocity distribution before comparing timing. The resulting distributions were effectively identical: median effective velocity was 6.461 m s^−1^ in the field data (interquartile range, IQR, 5.239–7.593 m s^−1^) and 6.460 m s^−1^ in the simulations (5.209–7.606 m s^−1^), and occupancy across the 2, 4, 6, 8, and 10 m s^−1^ reference bands did not differ (*χ*^2^ = 0.011, *df* = 4, *p* = 0.999985). Against this matched kinematic background, field and simulated calls followed the same broad inverse call-rate–displacement organisation (Figure 8aA–C). In plain terms, both datasets occupied the same range of movement between successive sensory updates.

**Figure 8.**
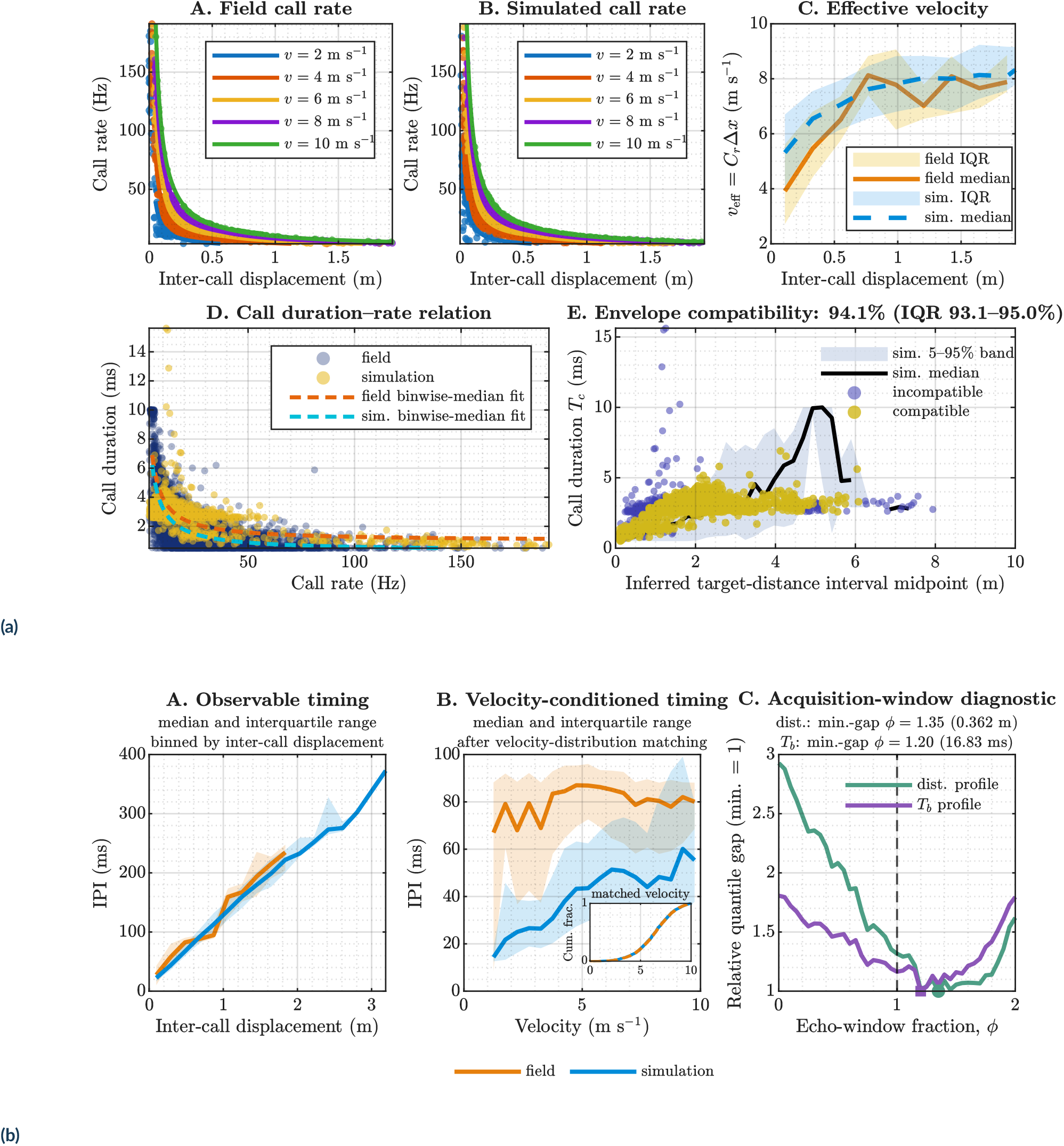
Field and simulated calls share broad movement and timing organisation. **a**, Shared observable timing, and duration compatibility. Panels A and B show the same inverse call-rate–displacement organisation in field and simulated calls; the effective-velocity distributions in panel C are nearly identical, establishing a common kinematic range. Panel D shows a shared reciprocal call-duration– rate relation, and panel E shows that 94.1% of field durations fall within the simulated 5–95% duration envelope under the stated conditional reconstruction. **b**, Timing and conditional diagnostics. Panel A shows the common increase in IPI with inter-call displacement. After matching the velocity distributions shown in the panel B inset, field IPIs remain longer than simulated IPIs, demonstrating that unequal flight-speed sampling does not explain the absolute timing difference. In panel C, distance and *T*_*b*_ discrepancies both decline as candidate *ϕ* extends beyond the original *ϕ* = 0–1 range, but reach different minima and do not disappear. A longer effective acquisition window can therefore move the model towards the field timing, but the available data do not uniquely distinguish this explanation from changes in responsivity or acoustic-anchor geometry. The dashed line marks *ϕ* = 1. Panels aE and bC assume *k*_*r*_ = 5 and reconstruct unobserved quantities from IPI; they are conditional diagnostics rather than independent measurements. Methods and assumptions are given in Sections 2.12.3 and 2.12.4; complete statistics are reported in Section 3.4 and Supplementary Table S7.

The corresponding IPI–displacement trajectories likewise overlapped through much of their common range: calls separated by greater movement were also separated by longer time in both datasets (Figure 8bA). This agreement establishes broad kinematic compatibility, but displacement is itself accumulated during an IPI and therefore cannot independently identify the timing policy. We consequently made a stricter comparison within shared velocity bins. The inset in Figure 8bB shows the empirical cumulative velocity distributions of the field calls and the sampled simulation calls; their near-complete overlap verifies the velocity match. Despite this matching, field IPIs remained systematically longer than simulated IPIs across much of the observed speed range. A combined displacement–velocity model explained 96.9% of IPI variance, but dataset-dependent terms remained significant (*p*≈ 0), confirming that the difference was not simply caused by unequal representation of slow and fast flight. Velocity accounted for the common kinematic support; it did not by itself account for the absolute field timing.

Call duration provided an additional and independent test. Rather than fitting the entire duration cloud, which is strongly broadened by behavioural and biological variation in the field, we compared the central duration–rate structure by fitting hyperbolae to the binwise median call durations as a function of call rate. This approach was used because the model predicts a reciprocal duration–rate dependence and the question of interest was whether the empirical central tendency followed the same structural curve as the simulations. The fitted scale terms were nearly identical between field and simulation (29.57 versus 29.62 ms Hz), while the fitted offsets differed mainly by an upward shift in the field data (0.99 versus 0.29 ms). The corresponding *R*^2^ values were 0.695 for the field medians and 0.977 for the simulated medians. This pattern shows that the field durations follow the same hyperbolic organisation predicted by the model, but are displaced upward and somewhat broadened, consistent with additional biological variability not explicitly encoded in the current simulation (Figure 8aD).

Finally, we tested whether measured field call durations were compatible with the distance-dependent duration envelope generated by the responsivity simulations when target distance was reconstructed from IPI-derived *T*_*a*_. The simulated 5–95% duration band was mapped across inferred distance, and each retained field call was classified according to whether its measured duration fell inside that envelope. Across 20 independently seeded ensembles, the median compatible fraction was 94.1% (IQR 93.1–95.0%; Figure 8aE). This result is especially important because it links the empirical durations not only to call rate, but also to the displacement-based distance structure implied by the framework, while quantifying the modest simulation-to-simulation variation in the estimated compatibility.

We next asked whether a longer effective acquisition window would move the model towards the residual field timing under fixed *k*_*r*_ = 5 and the assumed geometry. Candidate *ϕ* values from 0 to 2 were evaluated after matching 894 field and simulated calls at every value. Profiles compared the full percentile structure of conditionally inferred field distance with known simulated distance, and field-inferred *T*_*b*_ with simulated *T*_*b*_ (Figure 8bC). Both declined beyond the original *ϕ* = 0–1 range, reaching minima at *ϕ* = 1.35 for distance (mean quantile gap 0.362 m) and *ϕ* = 1.20 for *T*_*b*_ (16.83 ms). Under the non-negative-delay reconstruction in Equations 33 and 19, 1.79% of inferred field distances were clipped at zero. A complementary rank-matched diagnostic yielded a median required *ϕ* of 1.76 (IQR 1.05–2.22). Distributional values and the initial-distance sensitivity analysis are given in Supplementary Table S7.

The profiles therefore converged in direction rather than on one uniquely determined value: both indicated that the original simulations supplied too little effective acquisition time to reproduce the absolute field timing, but their optima differed and neither discrepancy vanished. Sensitivity profiling retained the global minimum at the default *d*_0_ = 5 m, although the best *ϕ* changed with assumed initial distance. The longer field IPIs can thus be represented as a conditional change in acquisition-window, responsivity, or anchor assumptions, but the present data cannot identify which variable changed. This is the intended diagnostic use of the model: shared structure establishes compatibility, while structured disagreement identifies the measurements needed to distinguish alternative control policies.

Taken together, the field analyses show that the responsivity framework captures the broad movement–timing and call-duration organisation expressed in natural sequences, while identifying the measurements needed to resolve the remaining latent variables. Such diagnosis would require call-resolved three-dimensional localisation from a task-optimised microphone array (Umadi, 2026d), synchronised tracking of the bat and a prey item of known position or trajectory, and measurement of call duration and flight kinematics. Direct reconstruction of sonar-beam aim would provide the strongest evidence for the effective acoustic anchor; where this is impractical, head direction may provide a useful proxy in oral-emitting frequency-modulated (FM) bats, for which tight head–beam coordination has been demonstrated during obstacle avoidance (Aoki et al., 2026). Simultaneous tracking of pinna orientation could further constrain the receiving acoustic field, because freely flying bats orient their pinnae towards prey and coordinate the emitted and receiving beams during high-speed foraging (Häfele et al., 2026). A controlled interception task using a presented mealworm could therefore provide, for every call, the emission position and time, relative velocity, candidate anchor geometry, and emitted duration. These measurements would allow competing *k*_*r*_, *ϕ*, and anchor-selection hypotheses to be fitted and tested against complete trajectories rather than inferred from IPI alone.

### 3.5 Buzz readiness identifies the onset of terminal organisation before a complete buzz is expressed

We next asked whether the framework could mark the onset of terminal organisation even when a complete buzz was not expressed. The buzz-readiness analysis comprised 1600 single-target, three-dimensional sequences (65,460 retained curve points) spanning *k*_*r*_, starting velocity, and initial call duration under the conditions in Table 1. We summarised readiness distance, sequence position, *δt*^∗^, and time to contact, and tested *δt*^∗^ as a proxy for terminal *T*_*b*,min_. Full grouped summaries and paired comparisons are reported in Supplementary Tables S8 and S9.

Buzz readiness shifted systematically with responsivity coefficient (Figure 9A). The median buzz-readiness distance declined from 1.98 m (IQR 1.43–3.16 m) at *k*_*r*_ = 3 to 1.37 m (1.01–2.00 m) at *k*_*r*_ = 5, 1.02 m (0.70–1.45 m) at *k*_*r*_ = 7, and 0.80 m (0.52–1.19 m) at *k*_*r*_ = 9. At the same time, the median interval *δt*^∗^ increased from 3.52 to 5.43, 6.78, and 8.72 ms. Thus, higher responsivity coefficients moved the onset of terminal organisation closer to the target while simultaneously expanding the characteristic timing interval associated with that readiness state.

**Figure 9.**
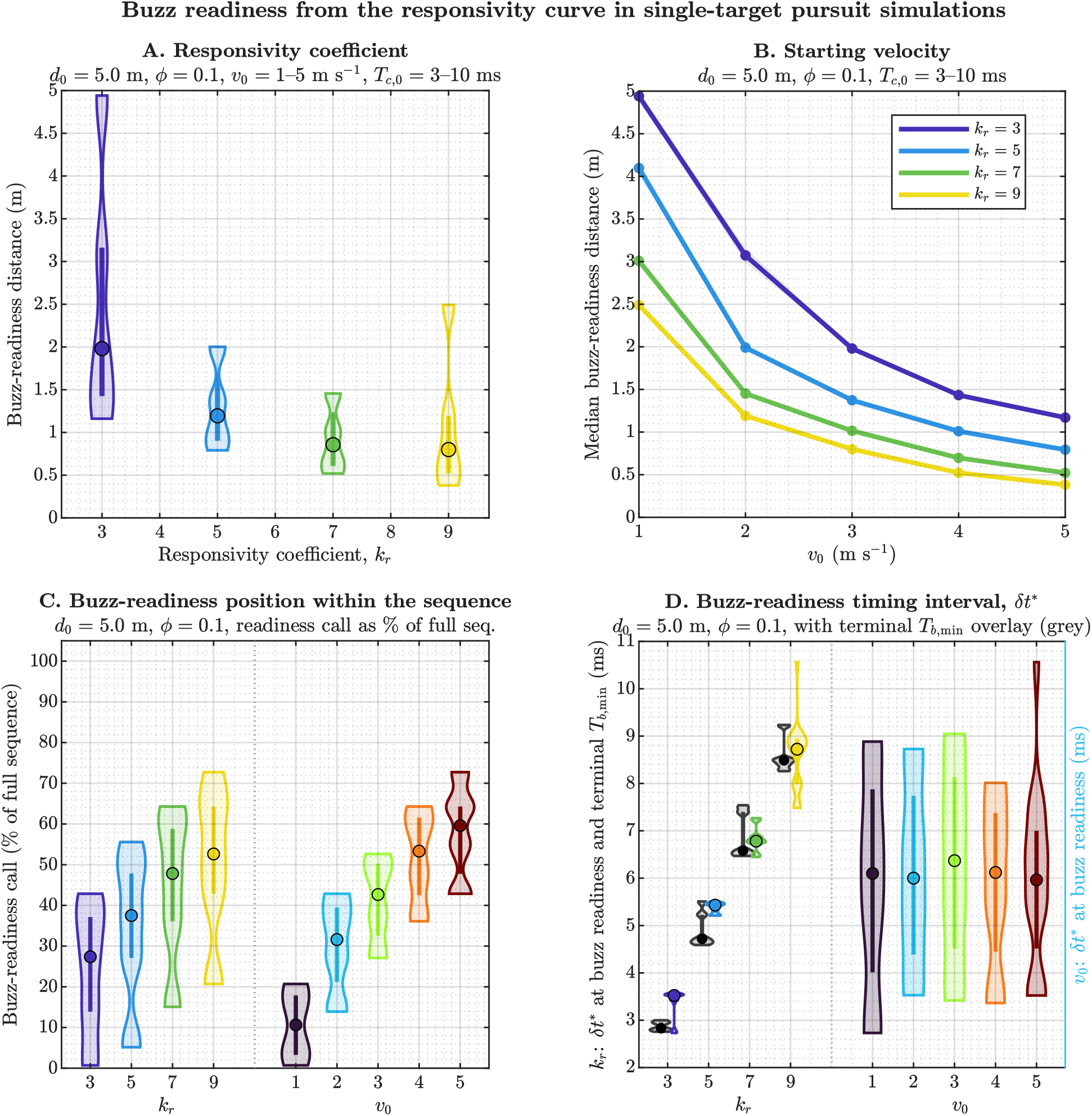
Buzz readiness provides a trajectory-based marker of the onset of terminal organisation. Panel A shows the distribution of buzz-readiness distances across responsivity coefficients, demonstrating that higher *k*_*r*_ shifts the onset of the terminally organised regime closer to the target. Panel B shows the corresponding dependence on starting velocity, with faster approaches reaching buzz readiness at shorter distances. Panel C shows the position of the buzz-readiness call as a percentage of full sequence length, separated by *k*_*r*_ and *v*_0_, indicating how the same readiness state can be reached at different points within an approach sequence depending on responsivity and kinematics. Panel D compares the readiness interval *δt*^*\**^ with the minimum terminal behavioural interval *T*_*b*,min_ for sequences that approached their responsivity-defined call-rate ceiling. The grey violins show realised terminal *T*_*b*,min_ values, whereas the coloured violins show *δt*^*\**^. Their close agreement at moderate to high *k*_*r*_ indicates that *δt*^*\**^ can serve as a practical proxy for the terminal behavioural interval, even when a complete terminal buzz is not expressed.

Starting velocity exerted a similarly strong effect on where readiness occurred, but not on *δt*^∗^ (Figure 9B,D). Median readiness distance fell from 3.55 m at *v*_0_ = 1 m s^−1^ to 0.66 m at 5 m s^−1^, while median *δt*^∗^ remained within 5.96–6.37 ms. Increasing velocity therefore compressed the spatial onset of terminal organisation without materially changing its characteristic timing interval.

Initial call duration had little influence: across *T*_*c*,0_ = 3, 5, 7, 10 ms, median readiness distance remained 1.28–1.29 m and the median sequence length remained 30 calls. The example curves likewise converged once distance-dependent timing was established (Figure 10C), indicating that readiness was governed principally by responsivity and kinematics rather than the duration assigned to the first call.

**Figure 10.**
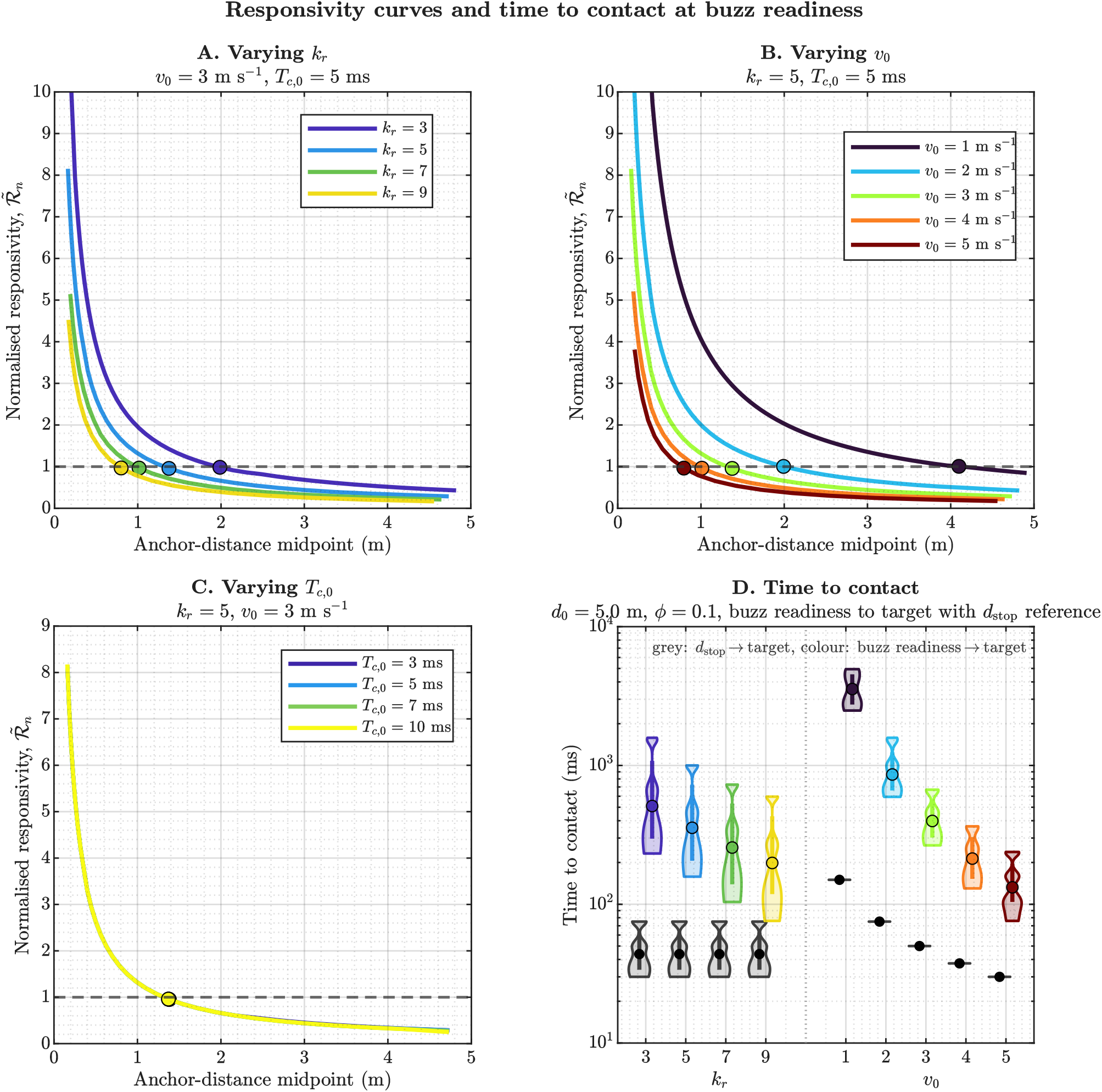
Responsivity curves and time-to-contact structure reveal what buzz readiness diagnoses. Panels A–C show representative normalised responsivity curves under sweeps of *k*_*r*_, starting velocity, and initial call duration. In all cases, buzz readiness is identified as the point where the normalised responsivity curve approaches its characteristic terminal level. The curves show that changing *k*_*r*_ or *v*_0_ strongly alters the distance at which this readiness point is reached, whereas varying initial call duration has relatively little effect once the approach enters closed-loop distance-dependent timing. Panel D compares time remaining from buzz readiness to target with the shorter time remaining from *d*_stop_ to target. The two quantities are clearly separated across both *k*_*r*_ and *v*_0_, showing that buzz readiness marks entry into a pre-terminal state rather than the final stopping boundary itself. This distinction is what allows the responsivity framework to identify terminal organisation even in sequences that do not culminate in a complete, definition-based terminal buzz.

The position of buzz readiness within the full sequence also varied systematically. Lower *k*_*r*_ values tended to reach buzz readiness later in the sequence but at larger distances, whereas higher *k*_*r*_ values reached readiness earlier in spatial terms but with fewer remaining calls before the end of the approach (Figure 9C). Across the starting-velocity sweep, faster approaches shifted the buzz-readiness call to a later percentile of the full sequence, consistent with a shorter and more compressed approach. These distributions show that buzz readiness does not simply reflect an absolute call-rate threshold; rather, it captures the point at which the responsivity curve indicates entry into the terminal high-update regime under the specific kinematic constraints of that approach.

The comparison with realised terminal *T*_*b*,min_ clarified when *δt*^∗^ can act as a practical proxy (Figure 9D). Agreement improved progressively with *k*_*r*_ : at *k*_*r*_ = 7, the median difference was − 0.045 ms (*p* = 0.484), and every sequence lay within 10% of *T*_*b*,min_; at *k*_*r*_ = 9, 80% did so. Thus, *δt*^∗^ is not universally identical to the terminal behavioural interval, but becomes a close proxy in more compressed terminal regimes.

Time to contact separated readiness still more clearly from the stopping boundary (Figure 10D). Across *k*_*r*_ = 3–9, median time remaining from readiness declined from 660 to 266 ms, whereas the corresponding *d*_stop_ reference was 50 ms. Across *v*_0_ = 1–5 m s^−1^, it declined from 3554 to 132 ms, compared with 150 to 30 ms from *d*_stop_. All paired differences were large (*p*≈ 0; Supplementary Table S9). Buzz readiness is therefore not a surrogate for reaching *d*_stop_, but a marker of entry into a pre-terminal state from which the final buzz may or may not unfold.

Taken together, these results show that the responsivity curve provides a useful way to diagnose the onset of terminal organisation without relying on an observed, complete, high-call-rate buzz. The buzz-readiness point moved predictably with both responsivity coefficient and approach speed, *δt*^∗^ captured the characteristic timing scale of this state, and the time-to-contact analysis showed that readiness occurred well before the final stopping boundary. In that sense, the framework replaces a purely threshold-based definition of terminal buzz with a trajectory-based description of when a bat enters the terminally organised regime of prey-capture behaviour.

### 3.6 Sonar strobe groups emerge under multi-target anchor switching

We next asked whether the same timing framework could generate intermittent sonar strobe group-like patterns (SSGs) (Moss and Surlykke, 2001). Six one- and two-target conditions separated target motion from anchor-selection mode (Table 2). Stable doublets, triplets, and quadruplets were detected post hoc when within-group IPI variability was below 5% and both flanking intervals were at least 1.2 times longer than the stable IPI.

**Table 2.** Single- and multi-target conditions and sonar strobe group emergence.

| Code | Scene | Target state | Anchor selection | SSG events | Sequences (%) |
| --- | --- | --- | --- | --- | --- |
| C1 | One target | Stationary | Fixed | 0 | 0 |
| C2 | Two targets | Stationary | Nearest | 0 | 0 |
| C3 | Two targets | Stationary | Random | 78 | 5.2 |
| C4 | One target | Moving | Fixed | 0 | 0 |
| C5 | Two targets | Moving | Nearest | 0 | 0 |
| C6 | Two targets | Moving | Random | 438 | 15.8 |

Conditions are arranged as parallel stationary and moving blocks, each progressing from one fixed anchor to two anchors under nearest or random selection. SSG-like events occurred only when two anchors were available and calls could be redistributed between them by random selection. Percentages give the fraction of simulated sequences containing at least one event.

Among 141,112 simulated calls, 1067 formed 516 SSG-like events. These events occurred exclusively in C3 and C6, yet within those conditions they were distributed across broad portions of the normalised call sequences rather than confined to a common or terminal phase (Figure 11A). The condition specificity rules out target motion and multi-target geometry alone, whereas the broad sequence positions show that the events were not merely a terminal consequence of sequence progression.

**Figure 11.**
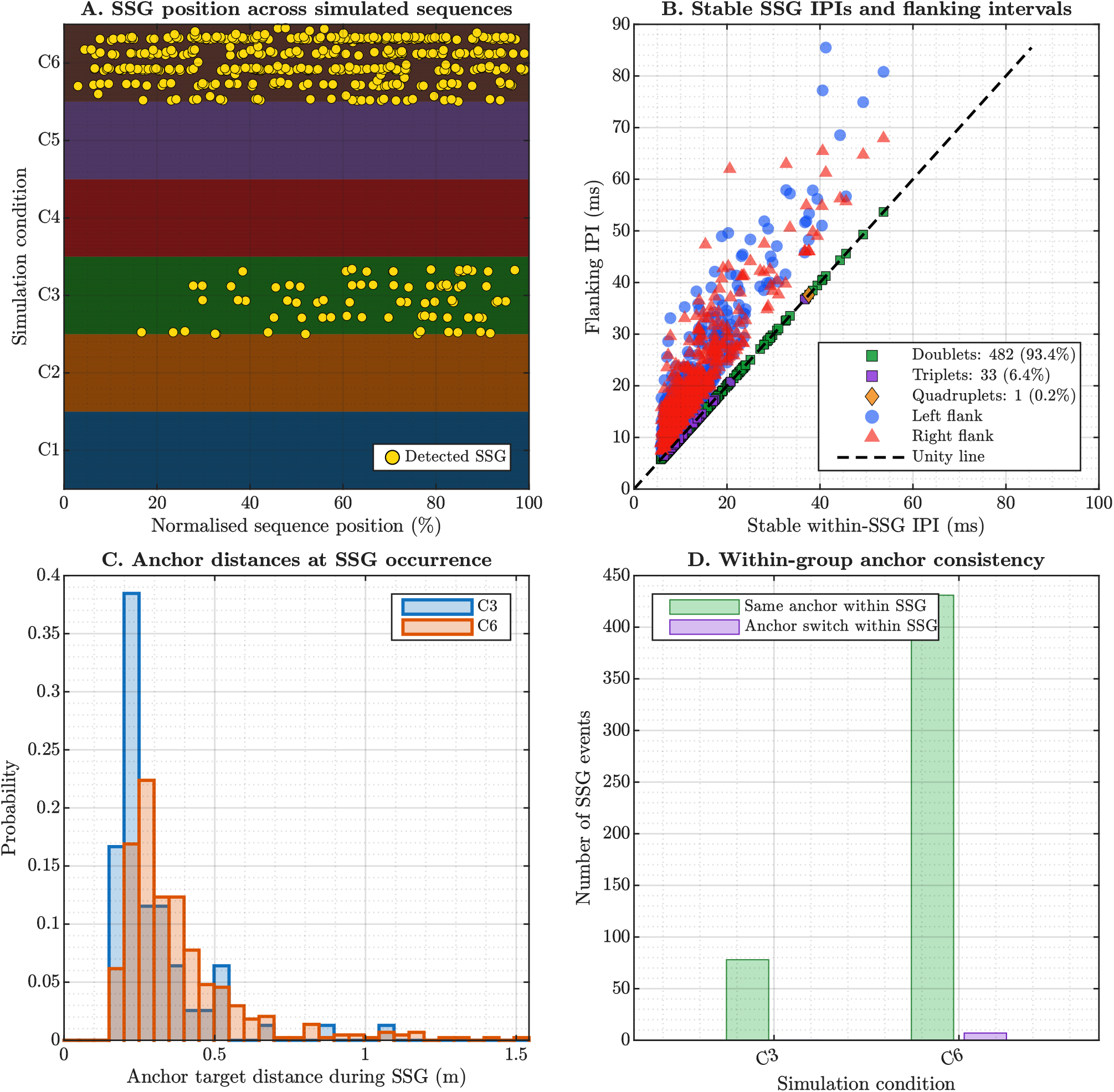
Sonar strobe groups emerge only in switching-rich multi-target conditions. The four panels summarise the SSG analysis from the single- and multi-target responsivity simulations. **A**, SSG events occurred only in the random-anchor two-target conditions, especially C6, but were distributed across broad portions of the normalised sequences rather than restricted to one phase. Their occurrence therefore required both multiple available anchors and intermittent anchor switching and was not simply a terminal-sequence effect. **B**, the within-group stable IPIs were bounded by longer flanking intervals, satisfying the defining temporal signature of an SSG without any explicit grouped-timing rule in the simulator. **C**, grouped events occurred at relatively short anchor distances, with C6 shifted to somewhat larger distances and longer stable IPIs than C3. **D**, once a group formed, calls almost always remained committed to the same anchor within that local event. Together, the panels show that sonar strobe groups can arise naturally when the responsivity framework operates in multi-target conditions that permit intermittent redistribution of calls across anchors.

SSG-like islands emerged only when multiple available anchors were combined with intermittent switching in the surrounding sequence. Short clusters dominated: 482 events were doublets, 33 triplets, and one a quadruplet. C6 events occurred at somewhat longer stable IPIs and greater anchor distances than C3 events (Figure 11B,C; Supplementary Table S10). Despite arising within switching-rich sequences, calls within a group almost always remained committed to one anchor (98.4% in C6; 100% in C3; Figure 11D). The detector therefore recovered brief islands of local regularity and anchor commitment, even though no grouping programme had been specified.

These results show that SSG-like temporal patterns can emerge within the responsivity framework without any explicit grouped-calling rule. The necessary condition was a behavioural sequence containing intermittent redistribution of calls across multiple objects, within which the bat temporarily concentrated successive calls on one anchor. In that sense, the grouped pattern reflects a local episode of anchor commitment embedded in a more variable multi-object sampling regime. The implication is that the SSG phenomenon can be explained as an emergent consequence of anchor selection and closed-loop timing, rather than as direct evidence for a dedicated grouping-specific motor programme. The broader interpretive implications of this result, especially in relation to earlier behavioural studies, are developed further in the Discussion.

Taken together, the analytical, simulation, and field-based results support a common conclusion: call timing during prey approach is well described as the outcome of a closed-loop interaction between acoustic acquisition, behavioural interval, and approach kinematics, rather than as a set of separate special-case behaviours. Across conditions, the responsivity framework recovered the expected call-rate–distance structure, showed how echo-window choice shifts IPI and call-duration trajectories, explained how terminal buzz structure emerges from the interaction between responsivity and closing velocity, identified a buzz-readiness state preceding full buzz expression, and reproduced sonar strobe group-like temporal patterns under multi-target anchor switching. Application to field recordings recovered the broad empirical movement– timing structure, isolated a residual timing difference after velocity matching, and used conditional parameter profiling to localise the assumptions capable of producing it. Together, these results support responsivity as a compact framework for linking spatial geometry, acoustic timing, and behavioural organisation across approach, interception, and target-selection contexts.

## 4 DISCUSSION

The temporal patterning of echolocation calls has previously been understood through overlapping vocal–motor and perceptual accounts. Call emission is often coordinated with respiration and wingbeat, while changes in repetition rate, stable interpulse-interval groups, and terminal calling have been interpreted as adaptive organisation of sensory sampling (Moss and Surlykke, 2001; Schnitzler and Henson, 1980; Suthers et al., 1972; Wong and Waters, 2001). Moss and Surlykke (2001) further showed that bats can integrate changing echo delays across calls and proposed that stable repetition may assist the organisation of echoes within a dynamic auditory scene. These accounts identify important physiological constraints and possible perceptual functions, but they do not specify a general event-level rule determining when one call should be followed by the next.

Apparent wingbeat–call synchrony also need not imply a dedicated coupling mechanism. It can arise when the timing imposed by delayed sensory feedback is compatible with an independently generated wingbeat rhythm; as interpulse intervals contract, that compatibility disappears and phase locking becomes infeasible (Umadi, 2026b). Differences in motor capacity may therefore shift the boundary across species, while the compressed timing of late approach and the terminal buzz makes decoupling especially likely (Stidsholt et al., 2021; Xia et al., 2025). Against this background, the present study identifies the timing opportunity within which these vocal, perceptual and motor processes operate. The responsivity framework decomposes each interpulse interval into acoustic acquisition and behavioural components, providing a common basis for examining call-rate progression, terminal call density and duration, nested response timescales, information-window and anchor selection, and the emergence of sonar strobe groups.

### 4.1 Call timing is jointly physical and behavioural

The event decomposition clarifies what an IPI represents. The interval between calls *n* and *n*+ 1 is initiated by call *n*: the first call generates the echo stream from which behaviourally relevant information is acquired, and the second marks completion of the resulting call cycle. Consequently, the range associated with an observed IPI is the range to the effective acoustic anchor at the onset of call *n*, not the range reached by the time call *n* +1 is emitted (Figure 1). This distinction becomes consequential at high closing velocities, because bat and target can move appreciably during a single interval. Equation 12 shows that an IPI contains a physically constrained acquisition interval and a biologically regulated remainder; Equation 19 further shows that the former depends on anchor distance, radial closing velocity, call duration, and the selected echo window.

The proportional relation *T*_*b*_ = *k*_*r*_*T*_*a*_ in Equation 25 supplies the missing link between those components. Once substituted into the IPI, it yields Equation 28 and the general call-rate identity in Equation 31. The resulting inverse-distance tendency is therefore not a descriptive hyperbola fitted after the fact; it is the observable footprint expected when successive calls close a delayed active-sensing loop. This interpretation is consistent with the simulations in Figure 6: *T*_*a*_ was largely insensitive to *k*_*r*_, whereas *T*_*b*_ changed strongly because *k*_*r*_ governs the biological allocation made after acoustic acquisition. Conversely, changing the echo-window fraction shifted both intervals, while changing velocity altered how quickly the system moved through the range-dependent timing trajectory. These separations are the main analytical advantage of the framework: several processes that can produce similar IPIs are represented by distinct terms rather than attributed indiscriminately to “call-rate adaptation”.

The term *responsivity coefficient* is deliberate. Although *k*_*r*_ is mathematically a proportional factor, calling it a gain would imply an amplifier-like transformation and obscure the biological content compressed into *T*_*b*_. Equation 24 makes clear that *T*_*b*_ may contain afferent transmission, sensory selection and integration, decision-related activity, vocal-motor preparation, and optional lag, with some components operating partly in parallel. The coefficient is therefore an effective systems parameter: it describes how much behavioural interval is allocated per unit acquisition time, not the gain of a known neuron or circuit. Delay-tuned neurons, pulse-rate-dependent changes in delay sensitivity, and the dynamic cortical representation of echo-acoustic flow demonstrate that neural processing is sensitive to echo delay, repetition rate, and approach history (Bartenstein et al., 2014; Suga and Horikawa, 1986; Wong et al., 1992). These findings make a scalable timing relation biologically plausible, but they do not identify *k*_*r*_ with any single neural latency or establish its circuit implementation.

A possible explanation for the approximate constancy of *k*_*r*_ over an approach is regulation of the information entering the loop. As calls shorten, the signal length and echo-overlap depth in Equations 20 and 21 contract. If this contraction reduces the quantity of scene information admitted for immediate processing, a shorter *T*_*b*_ need not imply that every constituent neural process has accelerated by the same factor. Instead, the bat may be processing a progressively restricted information set. Such restriction becomes increasingly valuable as relative motion makes older scene information stale. On this interpretation, proportionality is an emergent consequence of coordinated emission, acquisition, and selection. It is a hypothesis rather than a demonstrated mechanism, but it generates a direct test: manipulate call-defined information content or scene complexity while holding echo delay and relative motion constant, and ask whether the effective *k*_*r*_ departs systematically from its baseline value.

The field comparison supports this timing organisation but also shows why agreement in one observable plane is insufficient to identify its cause. Field and simulated calls occupied virtually identical velocity distributions, followed the same broad IPI–displacement structure, and shared a central reciprocal duration–rate organisation (Figure 8a). Yet field IPIs remained longer when calls were compared within the same velocity bins (Figure 8bB). The simulator therefore recovered the movement-dependent organisation without reproducing the complete absolute timing under its initial parameterisation. This residual is informative rather than merely unexplained: it has the direction expected if acoustic acquisition lasted longer, responsivity were greater, or calls were anchored to a different distribution of scene elements. Conditional profiling showed how much the assumed acquisition-window contribution would need to change under fixed *k*_*r*_ and geometry, thereby illustrating the framework’s diagnostic use. It did not determine which of those biological alternatives was responsible.

### 4.2 Responsivity and closing velocity define the terminal opportunity

The terminal buzz is often treated as a discrete behavioural phase because it is acoustically conspicuous. The present results instead place it at the short-range limit of the same timing relation that operates earlier in approach. Equation 45 shows that the terminal call-rate ceiling depends mathematically on *k*_*r*_, radial closing velocity, and the effective stop distance, but these variables do not exert comparable effects. The final spatial interval defined by Equation 47 is primarily a geometric quantity, whereas Equation 48 converts it into a temporal opportunity through *v*_*r*_. Equation 49 then gives the number of calls that fit into that opportunity. In the simulations, this division was clear: *k*_*r*_ strongly altered terminal call density, velocity strongly altered terminal duration, and late deceleration recovered near-target time that a fast constant-velocity approach would otherwise lose (Figure 7A–D). The resulting scale overlaps classic field measurements in *M. daubentonii*, for which the terminal phase averaged 156 ms and 21 calls (*n* = 37), with observed durations of 110–262 ms and sequences of 17–30 calls (Kalko and Schnitzler, 1989). Across three European pipistrelles, terminal phases likewise averaged 151–168 ms, spanned 90–356 ms, and culminated in Buzz II intervals of approximately 5.4–5.5 ms (Kalko, 1995). Correspondence in both terminal duration and update density does not identify the underlying *k*_*r*_ or velocity profile, but shows that the model-derived terminal time and call budgets occupy ranges measured across natural pursuits.

The asymmetry follows directly from the scales in the equations. At a fixed distance and *k*_*r*_, increasing *v*_*r*_ raises the first-arrival call rate only by the factor 1+ *v*_*r*_ /*c*; because biological closing velocities are much smaller than the speed of sound, this is a small correction rather than a proportional velocity control of call rate. By contrast, the time spent in a fixed terminal zone varies approximately as 1/ *v*_*r*_. Distance and responsivity therefore principally set the instantaneous call-rate trajectory, while closing velocity determines how quickly that trajectory is traversed and how long the bat remains near its terminal ceiling. Faster closure can raise instantaneous call rate slightly, but its much larger effect is to shorten the available terminal time and thereby reduce the number of calls that can be emitted; deceleration has the converse effect. The pronounced slowing documented during natural pipistrelle and bulldog-bat captures provides the required biological opportunity: by reducing radial closure, a late manoeuvre can extend the terminal sequence even when the local responsivity relation is unchanged (Kalko, 1995; Kalko et al., 1998).

This distinction resolves several apparently conflicting observations. Longer buzzes have been reported for more complex interception tasks and moving prey, leading to the interpretation that buzz duration indexes task difficulty or attention (Hulgard and Ratcliffe, 2016). Evasive prey can also elicit longer buzzes and rapid vocal adjustments (Foskolos et al., 2025), while prey evasiveness and masking noise can strongly increase terminal call rate in some species and foraging contexts (Ma et al., 2025). These effects are real, but they do not imply that task difficulty directly writes buzz duration into the vocal programme. A moving or evasive target can prolong pursuit, reduce or reverse radial closure, induce the bat to decelerate, or delay interception; each change increases the time spent in a high-update region. Masking or uncertainty can additionally alter the selected information window or the effective responsivity policy. Thus, ecological demand can shape the buzz through kinematics, anchor selection, and behavioural timing while remaining consistent with the physical opportunity described here.

The same reasoning helps interpret classic observations from clutter. In big brown bats, mean buzz duration declined from approximately 211 ms for a moving target in open space to 123 ms in the nearest-clutter condition, while mean buzz-onset distance shifted from approximately 62–65 cm in open space to 34 cm near clutter and distance travelled during the buzz declined from approximately 87 to 36 cm. Buzz-offset distance, by contrast, did not differ significantly among conditions (Moss and Surlykke, 2001). This combination is consistent with variable entry into and traversal of a terminal opportunity around a more constrained endpoint, although it is not a direct test of the model-defined stopping boundary. The pattern was attributed to limiting the mixing of echoes from successive emissions (Moss et al., 2006; Moss and Surlykke, 2001). The responsivity framework adds two non-exclusive possibilities. First, clutter may change which return terminates *T*_*a*_, preventing the earliest echo from being equated automatically with the prey echo. Second, shortening the call and selected echo window may restrict the effective acoustic depth being processed. Neither mechanism predicts that clutter must always shorten a buzz: the outcome also depends on trajectory, target–clutter geometry, and whether the bat prolongs pursuit. The framework therefore treats “clutter” as a set of changes to acquisition and control variables rather than as a unitary cause.

The buzz is consequently neither guaranteed nor universally necessary for interception. Fringe-lipped bats can capture airborne prey while reducing their call intervals without producing the high-resolution buzz characteristic of many vespertilionids (Mortensen et al., 2026). Even among aerial-hawking vespertilionids, species with different call designs and buzz structures partly converge when behaviour is considered relative to flight velocity, distance travelled between calls, and time to interception (Jakobsen et al., 2025). These observations fit the present separation: a bat may intercept successfully with a different *k*_*r*_, minimum acquisition interval, stop geometry, or kinematic trajectory, and therefore with a different terminal call density. The framework does not claim that all buzz features are passive consequences of propagation. Frequency shifts, beam broadening, respiratory coordination, and task-specific actions may all contribute. It claims more narrowly that any proposed terminal programme must operate within the update opportunity created by acquisition time and relative motion.

Physiological ceilings remain important within that opportunity. Superfast laryngeal muscles constrain the rate at which calls can be produced (Elemans et al., 2011), and species differ in their recovery from acoustic forward masking at very short intervals (Capshaw et al., 2024). These constraints can be represented by the attainable *C*_*r*,max_ and by speciesor context-dependent *k*_*r*_, but the present simulations do not derive either quantity from physiology. The equation-defined ceiling is therefore a system-level limit conditional on the selected stop distance and anchoring rule, not a substitute for vocal-motor or auditory measurements.

### 4.3 Buzz readiness, feedback, and behavioural adaptation occupy different timescales

The buzz-readiness analysis distinguishes entry into terminal organisation from the appearance of a complete, definition-based buzz. The readiness point defined by Equations 53–55 shifted with both *k*_*r*_ and starting velocity, whereas its associated interval, *δt*^∗^, remained comparatively stable across the velocity sweep (Figure 9A–D). Readiness also occurred well before the stop boundary (Figure 10D). It should therefore be interpreted as a state in which the call sequence has acquired terminal organisation, not as evidence that capture is imminent at a fixed time or distance. This distinction is especially useful for approaches that are aborted, redirected, or completed without a conventional buzz.

Classic field reconstructions provide an ordered set of behavioural landmarks against which this state could be tested. In European pipistrelles, presumed detection occurred at 1.14–2.20 m, the first vocal reaction at 0.80– 1.70 m, conventional terminal onset at 0.30–0.70 m, Buzz II onset at 0.09–0.41 m, and final emission cessation at 0.035–0.10 m (Kalko, 1995). Detection was assigned to the call preceding the first timing response, whereas reaction marked the beginning of the approach phase; neither quantity is therefore definitionally equivalent to buzz readiness. The sequence nevertheless supplies measurable candidate landmarks for determining whether the model-derived readiness point corresponds to an early vocal response, a later transition into terminal calling, or another trajectory-dependent event.

The distinction also prevents *T*_*b*_ from being misread as the time required for an entire observable response. In sudden prey-removal experiments, median acoustic and capture-behaviour reaction times during aerial interception were 87 and 82 ms, respectively, whereas the corresponding values during trawling were 123 and 178 ms (Geberl et al., 2015). These intervals were measured from physical prey removal to the first observable interruption of the buzz or modification of capture behaviour. They therefore include an unknown wait until a call first supplied evidence of prey absence, echo propagation, any sensory accumulation and decision process, and the initiation of a measurable motor change. The analysis did not identify the first informative call–echo pair and consequently cannot establish whether the bat updated its target estimate after one missing echo or only after evidence accumulated across several calls. These values are neither minimum auditory latencies nor direct measurements of *T*_*b*_. The longer trawling latencies also need not indicate slower acoustic processing. They may instead reflect differences in motor commitment, task geometry, physical manoeuvres, and the multimodal feedback required for trawling and aerial interception. The results nevertheless show that an observable vocal or capture adjustment can span several call cycles. Echospace analyses have likewise found that reconstructed echo direction can precede measurable flight turning by several call cycles, although such lagged correlations do not by themselves establish the causal contribution of an individual echo (Teshima et al., 2022).

This timescale separation can begin well before the final buzz. The call-rate–distance identity (Figure 4) implies that, for *k*_*r*_ = 5 in the first-arrival case, IPIs of 80–100 ms correspond to anchor distances of approximately 2.3–2.9 m under the stationary-delay reference. Including a closing velocity of 5 m s^−1^ changes these values only slightly, to approximately 2.32–2.90 m. Calls can therefore recur more rapidly than the median latency of an observable prey-removal response well before the conventional terminal phase.

The comparison requires precision about what is meant by closed-loop control. In its general control-theoretic sense, a system is closed-loop when measurements of its present or past state influence subsequent control inputs; the feedback may be delayed and need not produce a completed response before another measurement is acquired (Oppenheim and Verghese, 2017). Short IPIs therefore do not make the echolocation system open-loop. What becomes untenable is the stronger interpretation that the echo from every call must produce a completed vocal or capture response during the immediately following call cycle. Conversely, a call–echo–call sequence demonstrates only that feedback was temporally possible, not that the received echo causally governed the next emission. Calls generated by an internal schedule or kinematic policy could receive echoes “in time” without their timing being determined by those echoes, producing the buffered pseudo–closed-loop regime described previously (Umadi, 2026b).

Delayed, multi-cycle, or hierarchically nested feedback is therefore one possible interpretation of how responsivity-defined timing opportunities are used, but such an architecture is not itself part of the responsivity framework. The framework specifies the cadence and decomposition of local update opportunities; it does not specify the state estimator, decision threshold, memory process, causal feedback lag, or flight controller operating across them. Equation 24 treats *T*_*b*_ as the composite post-acquisition portion of one call cycle, not as the time required to complete every higher-order consequence of the acquired information. Echo information could update a sensory representation over successive calls and influence later vocal or capture actions, consistent with behavioural evidence for accumulation across acoustic snapshots (Salles et al., 2020). Experiments with a fishing bat further showed that capture could be directed towards the extrapolated position of a moving target after it became acoustically inaccessible, indicating that overt action can be guided by a target-state estimate retained from earlier echoes (Campbell and Suthers, 1988). During the buzz, an 80–100 ms overt response (Geberl et al., 2015) may encompass many calls even though the IPI is only a few milliseconds. Information from the first informative echo could therefore update an internal target-state estimate while the ongoing vocal or capture programme remains outwardly unchanged; alternatively, several echoes may be required before sufficient evidence accumulates to initiate a response. The terminal region may consequently be increasingly constrained by feedback at the level of sensory acquisition and vocal timing while remaining multi-cycle at the level of flight and capture adaptation. Neither the absence of an immediate behavioural deviation nor the later appearance of a response establishes which intervening echoes were used.

This interpretation also reframes the “point of no return” near interception (Geberl et al., 2015). It need not be a single physiological instant at which sensory processing stops, and it may differ among behavioural outputs. Following late prey removal, bats could retain a nearly complete vocal sequence while aborting aerial capture movements, whereas trawling movements persisted longer when water contact continued to provide somatosensory feedback. These dissociations indicate output-specific limits imposed by decision thresholds, motor commitment, task geometry, and available feedback. A point of no return may therefore mark the stage at which the remaining terminal opportunity is too short for newly acquired information to alter a particular motor outcome, even though sensory acquisition and other behavioural channels continue to update. Responsivity curves provide a trajectory-specific means of locating the preceding transition into compressed control, whereas prey-removal and other perturbation experiments probe how many subsequent updates are required before each form of overt response becomes measurable. Combining these approaches could separate the timescale of local timing control from the timescales of sensory integration, decision formation, and task-specific behavioural execution.

### 4.3 Information sufficiency, call duration, and the acoustic anchor

A longstanding open question in bat echolocation is which object, echo feature, or portion of the returning echo stream governs the timing of the next call. This question is central to understanding echolocation control policies because it concerns both how a bat selects behaviourally relevant information from a multi-object acoustic scene and when that information is considered sufficient to support the next sensory update. Natural calls return cascades of echoes from prey, vegetation, ground, and other objects, and a single emission can produce multiple spatially distributed echo-incidence points whose directions need not coincide with the emitted beam (Teshima et al., 2022). Reflectors separated by less than the overlap depth in Equation 21 contribute to a composite waveform rather than to completely isolated echo packets. Equation 10 therefore defines *T*_*a*_ by the end of behaviourally selected acoustic information, and Equations 17 and 19 permit that endpoint to lie at first arrival or later within the returning stream. The strong displacement of IPI with *ϕ* in Figure 5 shows that this choice should leave a measurable signature. It also shows why *k*_*r*_ and *ϕ* can be confounded: extending the acquisition window increases the IPI and its derived *T*_*b*_ even when the responsivity coefficient is unchanged.

The field diagnostic provides an empirical illustration of this confounding. After velocity matching, both the anchor-distance and *T*_*b*_ profiles improved as *ϕ* was extended beyond the 0–1 range used in the original comparison simulations (Figure 8bC). Under fixed *k*_*r*_ = 5 and the assumed geometry, the field timing therefore required a larger effective acquisition-window contribution than those initial simulations supplied. This is not evidence that the bats literally used one common *ϕ* > 1. A later information-sufficiency event, a more distant or changing anchor, a larger effective *k*_*r*_, an omitted lag, or heterogeneity among species and pursuit geometries could all lengthen the observed IPI. The fact that the distance and *T*_*b*_ diagnostics shifted in the same direction but reached different optima makes the central point: timing data can constrain plausible control policies, but cannot uniquely identify the selected acoustic information without independent spatial and echo measurements.

Neural evidence provides plausible candidate rules but does not yet identify the behavioural anchor. Cortical neurons in passively stimulated bats predominantly represented the first echo of a cascade, even when later echoes were more intense, with later responses suppressed unless the leading echo was too weak to drive the neuron (Beetz et al., 2026). Related work shows that cortical representations of multiple-object echo streams depend on natural sequence context and the changing echo-acoustic flow (Greiter and Firzlaff, 2017). These findings support a strong first-arrival bias at the cortical population level. However, the first-echo experiments do not show that an actively behaving bat must time every subsequent call from the nearest object. Bats can initiate distance-appropriate approaches despite closer background objects (Amichai and Yovel, 2017), sequentially shift acoustic gaze between an obstacle and prey (Surlykke et al., 2009), and distribute sonar attention across current and future prey during consecutive captures (Fujioka et al., 2016; Sumiya et al., 2017). In the obstacle–prey experiment, the mean shift of beam aim towards the prey occurred 220 ± 140 ms before net crossing, while the call-duration increase that admitted echoes beyond the net occurred 219± 79 ms before crossing; the timing of the two changes was correlated (*R*^2^ = 0.59) (Surlykke et al., 2009). Echo perception, cortical response dominance, and the echo that governs call timing should therefore be treated as related but distinct questions.

Call duration offers a potential means of regulating this distinction. Equations 20 and 21 show that shorter calls reduce the spatial extent of the emitted waveform and the range separation over which echoes overlap. This gives call shortening a possible role beyond avoiding overlap between the outgoing call and the first returning echo: it may narrow the acoustic information aperture from which an anchor is selected. Field reconstructions show that *M. daubentonii* and European pipistrelles progressively shortened calls with target range and largely avoided outgoing-call overlap with prey echoes; pipistrelles additionally ceased emission several centimetres before contact (Kalko, 1995; Kalko and Schnitzler, 1989). These observations strongly support the physical no-overlap constraint, while not by themselves demonstrating that duration controls which later information is selected. Longer calls, conversely, deliver more energy and may maintain detection range, so their persistence at long range is not paradoxical. The useful trade-off may be between acquiring a broad, energetic view of the scene and restricting the information packet when precise target-centred updating becomes urgent. Behavioural evidence that bats shorten calls in clutter and coordinate call duration with shifts in range-axis acoustic gaze is consistent with this interpretation (Surlykke et al., 2009; Wilkinson et al., 2025), but it does not yet demonstrate active control of an integration window.

The gated time-window account developed from distance-discrimination experiments in *Noctilio albiventris* is especially relevant here. Interfering signals disrupted distance judgements only when they occurred within a call-related temporal window, and the relative timing of constant-frequency (CF) and frequency-modulated (FM) components was more consequential than CF duration alone (Roverud and Grinnell, 1985). A threshold-sensitive closure of such a window would provide one possible implementation of the selected endpoint in *T*_*a*_: the earliest sufficiently informative event could close the interval even while later echoes remain physically present. The responsivity framework does not require this particular mechanism, but it converts the broad question “when does the bat have enough information?” into a measurable timing problem.

Several experiments follow. Echo delay and relative motion could be held constant while emitted-call duration, target–clutter separation, echo amplitude, or the order of competing echoes is manipulated. If call duration merely prevents pulse–echo overlap, timing should follow the first relevant delay once overlap is avoided. If it regulates information admission, the effective *ϕ* and inferred *k*_*r*_ should change with scene content even at the same delay.

A second design could place a rewarding target behind a nearer, non-rewarding reflector and independently vary which echo is first, strongest, or most predictable. Simultaneous beam reconstruction, head and pinna tracking, and call-resolved reconstruction of the echoes reaching the bat (Teshima et al., 2022) would then test whether the next IPI is anchored retrospectively to an acquired echo, prospectively to an expected target return, or to a dynamically selected portion of the composite stream.

Physical-target paradigms cannot, however, provide direct and independent control over the echo waveform received by a freely flying bat, because altering a reflector generally changes several acoustic and geometric properties together. Addressing this limitation calls for a lightweight, low-latency, programmable *ad aures* (at-the-ears) augmented-reality system that attenuates uncontrolled natural returns and delivers call-triggered synthetic or filtered echoes, thereby controlling the auditory information available to the bat during free flight. Such a system would extend current recording and playback approaches towards call-by-call manipulation of return delay, level, spectrum, temporal order, and admitted echo-window content. By placing the auditory scene received by the bat under programmable control during free flight, this approach could move the study of echolocation timing from correlational inference towards causal tests of the information and control variables that organise behaviour.

### 4.2 Sonar strobe groups as transient organisation within a multi-anchor scene

SSGs have been interpreted as purposeful increases in spatial or temporal sampling during difficult tasks (Kothari et al., 2014; Moss et al., 2006; Moss and Surlykke, 2001). In big brown bats navigating obstacle arrays, dense clutter produced shorter IPIs and call sequences dominated by doublets and triplets; the alternation between short within-group and longer between-group intervals was interpreted as balancing rapid close-range obstacle updating against longer-range path sampling (Petrites et al., 2009). SSGs also occur during cluttered interception and the tracking of unpredictable targets, and may appear without an increase in the total number of calls, indicating temporal rearrangement rather than simple call addition (Kothari et al., 2018). This association has now been reproduced in a controlled target-motion paradigm: the median proportion of calls assigned to SSGs increased significantly from 4% during predictable motion to 5% during unpredictable motion, but the respective ranges, 1–9% and 3–13%, overlapped substantially and SSGs occurred under both conditions (Cadigan et al., 2026). The accumulated experiments therefore establish that uncertainty and acoustic complexity modulate SSG prevalence, but neither unpredictability nor clutter is a necessary event-generating condition. The literature has converged more strongly on the circumstances that elicit additional grouping than on the call-resolved process that creates the groups themselves. Hulgard and Ratcliffe (2016) consequently proposed that the terminal buzz can be viewed as an extended, call-rich SSG. The present simulations suggest the complementary possibility that brief SSGs can be transient, quasi-terminal episodes of local anchor commitment embedded within a broader, less regular sampling sequence.

The crucial result is one of sufficiency, not exclusivity. SSG-like islands occurred only when two anchors were available and the simulation permitted random call-to-call anchor selection; one-target and nearest-anchor conditions produced none (Figure 11A). Once an island formed, its calls almost always remained on the same local anchor (Figure 11D), even though switching occurred in the surrounding sequence. Thus, no specialised group generator needs to be assigned to real bats on the basis of this result. Random switching was an intentionally simple implementation used to demonstrate that intermittent redistribution plus temporary commitment is sufficient to satisfy the temporal island criterion.

This matters because the criterion identifies a pattern, not a unique behavioural computation. It requires similar within-group intervals bounded by longer intervals, so it preferentially detects conspicuous local islands and may miss comparable changes in anchor allocation that lack both flanks. Conversely, a sequence can satisfy the criterion through different processes: repeated sampling of one anchor amid alternation, compensation between velocity and range, or recovery following a violated echo expectation. The predominance of doublets and rarity of longer groups in the simulations underline this ambiguity (Figure 11B). Under independent random allocation, the probability of an uninterrupted run on one of several anchors declines as the run length increases; the observed group-size distribution additionally reflects the changing geometry and the requirements for stable within-group timing and two longer flanks. A single additional call on the same anchor can therefore turn a zig-zagging multi-object sequence into a detected doublet without changing the underlying timing law, whereas progressively longer stable islands become increasingly uncommon.

Kinematics can help discriminate among these explanations. Under continued radial closure, repeated sampling of the same object should normally shorten the acquisition interval from one call to the next. Nearly equal intervals are therefore difficult to attribute to an unchanged moving anchor unless relative motion is negligible or changes in velocity, geometry, or the selected information window compensate for the range change. An earlier-than-expected return can shorten the ensuing acquisition interval, whereas a later return from a receding target should lengthen it if the responsivity relationship and anchor remain unchanged. Either event can displace an individual IPI, but neither alone explains a run of two or three nearly equal IPIs bounded on both sides by longer intervals. This observation does not identify an alternative anchor by itself, but it makes stable intervals a diagnostic prompt: the analyst should ask which object and motion state would make those intervals compatible with Equation 29. The anchor-distance and within-group consistency patterns in Figure 11C–D demonstrate how this calculation can be applied when the simulated anchors are known.

The prediction-recovery interpretation proposed for unpredictable motion is compatible with this account at a higher level, but it answers a different question. Cadigan et al. argue that target reversals conflict with an internal model of approaching prey motion, prompting the bat to collect additional information and delaying its transition from tracking to capture (Cadigan et al., 2026). Their “across-timescales” result places a local change in vocal organisation alongside later changes in capture-related calling and pinna position; it does not specify how a violated expectation generates the stable intervals and longer boundaries of an individual SSG. Complementarily, their SSGs occurred over a broad range of target distances under both motion conditions, and the simulated events were distributed across broad portions of the normalised sequences (Figure 11A). Empirical and simulated grouping is therefore not confined to a single trajectory phase. Prediction error may help determine when sampling is reorganised, whereas responsivity and anchor allocation provide a candidate account of the temporal form that reorganisation takes.

Previous behavioural results are compatible with, but do not prove, the anchor-allocation interpretation. Big brown bats redirect their acoustic gaze sequentially across nearby objects (Surlykke et al., 2009); target tracking near clutter is accompanied by shorter calls, more SSGs, and greater head movement (Wilkinson et al., 2025). Likewise, the rewarding target in unpredictable-motion experiments is usually assumed to remain the anchor throughout the trial, but the object governing each IPI was not measured (Cadigan et al., 2026; Kothari et al., 2018). Increased uncertainty could promote repeated verification of the prey, redistribution among scene elements, or recovery from a violated target-state prediction. These alternatives operate above the responsivity rule: responsivity constrains when another update can occur, while acoustic gaze and anchor selection determine where that update is allocated.

A decisive experiment would therefore identify the anchor of each call interval rather than ask which target the SSG as a whole “follows”. A bat could track a rewarded target whose motion alternates between predictable and erratic trajectories while a non-rewarding distractor is presented concurrently; the distractor’s predictability, echo arrival order, and echo level could be varied independently of reward. Reconstructed candidate echoes could then be compared with each observed next-call onset, while call-resolved beam direction and head orientation establish how acoustic gaze was allocated. The anchor-allocation account makes a local prediction: calls in the surrounding sequence may sample different objects, but the stable intervals forming an SSG should represent a temporary run governed by one effective anchor, with changes in anchor or acoustic gaze occurring preferentially near the longer flanking intervals. A target-state-recovery account instead predicts that the onset of grouping should follow unexpected changes in the rewarded target’s trajectory and that this target’s echo should remain the best predictor of within-group call timing, even when the distractor provides an earlier, stronger, or more predictable return. This design would distinguish temporary anchor commitment from target-specific recovery without treating the operationally identified group as a behavioural entity or assuming a formal switching probability.

### 4.6 Comparative scope, inference, and limitations

Species differences provide an important test of whether *k*_*r*_ is biologically informative. Temporal resolution differs at early auditory stages between bats with different echolocation behaviours (Capshaw et al., 2024), and species with markedly different call designs and flight speeds reorganise their approach sequences in relation to their locomotor dynamics (Jakobsen et al., 2025). Within the framework, variation in maximum call rate can arise from differences in the minimum attainable *T*_*a*_, stop geometry, radial velocity, or *k*_*r*_ rather than from the presence or absence of a categorically distinct buzz programme. This accommodates high-rate Buzz II sequences, shorter buzzes, and successful aerial captures without a prominent high-resolution buzz (Mortensen et al., 2026). It also makes *k*_*r*_ a candidate comparative trait for species, guilds, or tasks, although establishing that status requires repeated estimates under matched geometry and behavioural context.

The terminal region is likely to be the most informative phase for such estimation because call duration and the selected echo-window contribution are small, while anchor distance and interception geometry can in principle be measured. Rearranging Equation 45 yields *k*_*r*_ only if *d*_stop_, *v*_*r*_, and a genuinely attained terminal ceiling are known. If the observed maximum call rate falls below the physiological or behavioural ceiling, it provides only a lower bound on that ceiling and the corresponding estimate can overstate *k*_*r*_. A worked reanalysis of published *Noctilio leporinus* sequences illustrates the approach: terminal call-rate anchoring yielded similar estimates for two individuals (*k*_*r*_ ≈ 3.6–3.7), after which independently digitised IPIs implied cue distances substantially shorter than the reported prey distances over much of the approach (Hartley et al., 1989; Umadi, 2026a). This example is diagnostic rather than definitive, because the original sequences did not directly identify the echo source. Mid-approach estimates face the additional confounding shown by Equation 32: increasing *ϕ* and increasing *k*_*r*_ can both lengthen an IPI. Comparative work should therefore fit entire call-resolved trajectories, include call duration and relative motion, and test whether a common *k*_*r*_ remains adequate before assigning species- or individual-level values.

The same caution applies to state inference. Equations 39 and 40 allow anchor range or radial velocity to be recovered when the other quantities and *k*_*r*_ are known, but these are conditional identities under first-arrival anchoring. They cannot turn call timing alone into an unambiguous measurement of prey distance if the effective anchor changes, if *ϕT*_*c*_ is non-negligible, or if the bat is sampling several objects. Their strongest use may therefore be diagnostic: candidate anchors can be tested for consistency with the observed timing, known trajectory, and plausible *k*_*r*_, rather than accepting the experimentally designated target by default.

Extension beyond low-duty-cycle FM echolocation requires redefining the information anchor, not necessarily abandoning the timing principle. High-duty-cycle CF and CF–FM bats experience call–echo overlap as a normal operating condition and obtain important velocity and flutter information spectrally (Fenton et al., 2012; Neuweiler, 2003). Their relevant *T*_*a*_ may therefore end at a finite information-accumulation event rather than at the end of a discrete echo packet. FM components can still provide temporal registration for range, as indicated by the gated distance-discrimination results in *Noctilio* (Roverud and Grinnell, 1985), and fish-catching *Noctilio* substantially modify signal design during terminal capture (Hartley et al., 1989). The present equations may thus remain useful if the anchor is defined by sufficient information rather than by complete echo reception – a prospective generalisation.

Several further limitations bound the present claims. The simulator reduces biological processing to an effective coefficient and does not model auditory thresholds, beam and pinna transfer functions, echo intensity, explicit attention, learning, state estimation, respiratory coupling, or vocal-motor physiology. The SSG implementation uses random anchor selection as a sufficiency test and is not a behavioural model of how bats choose objects. The field sample combines unidentified *Pipistrellus* and *Myotis* sequences and lacks prey trajectories and call-resolved echo anchors. Its conditional profiles also impose *k*_*r*_ = 5 and an assumed initial geometry: *k*_*r*_, *ϕ*, initial distance, anchor identity, and unmodelled lag can partially compensate for one another. The *d*_0_ sensitivity analysis showed that the default 5 m starting distance was not solely responsible for the best-fitting region, but the fitted *ϕ* nevertheless changed across candidate starting distances. Moreover, field-inferred distance and *T*_*b*_ are transformations of IPI under these assumptions rather than independent measurements. The profile results should therefore be read as constraints on candidate explanations, not as recovered behavioural parameters. Buzz readiness depends on a model-defined call-rate ceiling and should be validated against independent perturbations rather than treated as a discovered physiological threshold. Finally, an approximate proportional relation does not require *k*_*r*_ to remain constant throughout an entire pursuit. Systematic changes in task, information content, call design, or behavioural state may produce phase-specific coefficients, and such departures are potentially informative rather than failures of the framework.

The immediate value of responsivity theory is therefore methodological as much as explanatory. It specifies what must be measured to distinguish competing accounts: the distance and identity of the effective anchor at call onset, radial relative velocity, emitted duration, candidate echo-window endpoint, and the next call time. With these variables, experiments can ask whether a longer buzz reflects additional terminal time or denser updating; whether clutter changes *k*_*r*_, *ϕ*, or the anchor; whether an SSG is a local run on one object or a distinct timing mode; and how many local updates precede an observable motor decision. These questions move the analysis from naming temporal patterns to testing the event structure that generated them.

## 5 CONCLUSIONS

The principal contribution of the responsivity framework is to shift the unit of explanation in echolocation from named call patterns to the call-to-call sensorimotor cycle that generates them. Within this cycle, acoustic acquisition determines when relevant information can become available, responsivity determines the temporal allocation preceding the next update, and movement determines how many such updates remain possible. The terminal buzz and transient call groups can therefore be understood as different observable organisations of a common timing process rather than as mechanisms in themselves.

This distinction is also methodologically important. Similar call sequences may arise from different combinations of anchor selection, information-window closure, responsivity and approach kinematics; temporal pattern alone is therefore insufficient to identify the underlying control policy. By expressing these alternatives as separable and measurable components, responsivity provides a basis for testing which information governs each call interval and how that information is converted into subsequent action. More broadly, the framework suggests that adaptive sensing is organised not only around what information is present, but around when selected information becomes sufficient to support another behavioural update.

Responsivity theory therefore provides a route from observed temporal patterns to experimentally discriminable hypotheses. By identifying the quantities that must be measured or manipulated, the framework enables candidate control policies to be evaluated directly. Its broader value lies in providing a general basis for investigating how active-sensing behaviour is organised in time.

## REVISION SUMMARY

This version adopts the title *A responsivity theory of call timing patterns in bat echolocation*, which more directly summarises the scope of the framework. Earlier versions were titled *Biosonar Responsivity Sets the Stage for the Terminal Buzz* and *The Temporal Logic of Bat Echolocation: A Responsivity Theory of Call Timing Patterns*; these are available through the bioRxiv version history.

Relative to the preceding version, the manuscript clarifies the empirical motivation and conditional scope of the proportional timing relation in Equation 25, distinguishing an effective approximation within a behavioural state or approach segment from its conditional distance-scaling predictions. The equations and analyses remain unchanged. The title, abstract and presentation have been refined, and supplementary material is included at the end of this document.

## ABBREVIATIONS

AC: alternating current
AD: analogue-to-digital
CF: constant frequency
DA: digital-to-analogue
DC: direct current
FM: frequency modulated
IPI: interpulse interval
IQR: interquartile range
MAD: median absolute deviation
SSG: sonar strobe group.

## 6 DECLARATIONS

### Ethics Approval and Consent to Participate

Not applicable. The field data were obtained through passive acoustic recordings of free-flying wild bats. No animals were captured, handled, or experimentally manipulated, and no human participants were involved.

### Consent for Publication

Not applicable.

### Data Availability

The analysis code, processed field dataset, and supporting simulation and analysis outputs are available in the Zenodo archive (Umadi, 2026c). The archived release is version V1.1.0, DOI:https://doi.org/10.5281/zenodo.21834638. The processed dataset provides the reproducible starting point for the reported field analyses; the release does not provide a fully automated reconstruction of the manually curated dataset from the original multichannel recordings.

### Authors’ Contributions

**RU:** Conceptualisation, Methodology, Software, Validation, Formal Analysis, Investigation, Data Curation, Writing – Original Draft, Writing – Review & Editing, Visualisation, Project Administration.

**UF:** Supervision, Validation, Resources, Writing – Review & Editing, Funding Acquisition.

All authors read and approved the final manuscript.

## Acknowledgments

We thank our colleagues at the Technical University of Munich (TUM) for their valuable discussions and feedback, and Christian Fink for his technical assistance in constructing the field array. We are grateful to all reviewers who engaged with this and previous versions of the manuscript. Their critical feedback and constructive engagement helped reshape the work into a comprehensive theoretical model. We also thank the editors for their valuable judgements and careful handling of the manuscript throughout the review process.

## Abbreviations

CF: constant frequency
FM: frequency modulated
IPI: interpulse interval
IQR: interquartile range
MAD: median absolute deviation
SD: standard deviation
SSG: sonar strobe group
TTC: time to contact.

## Supplementary Information

References to equations, figures, and sections without an S prefix refer to the accompanying main manuscript.

The supplementary material first provides a notation glossary and a conditional analytical reference for applying the responsivity framework. These are followed by selected derivations and the complete secondary numerical results supporting the streamlined accounts in the main text. Quantities described as recoverable below are conditional on the listed inputs and assumptions; none of the inverse identities resolves several unknown latent variables from call timing alone.

## S1 NOTATION AND PARAMETER GLOSSARY

**Table S1.**
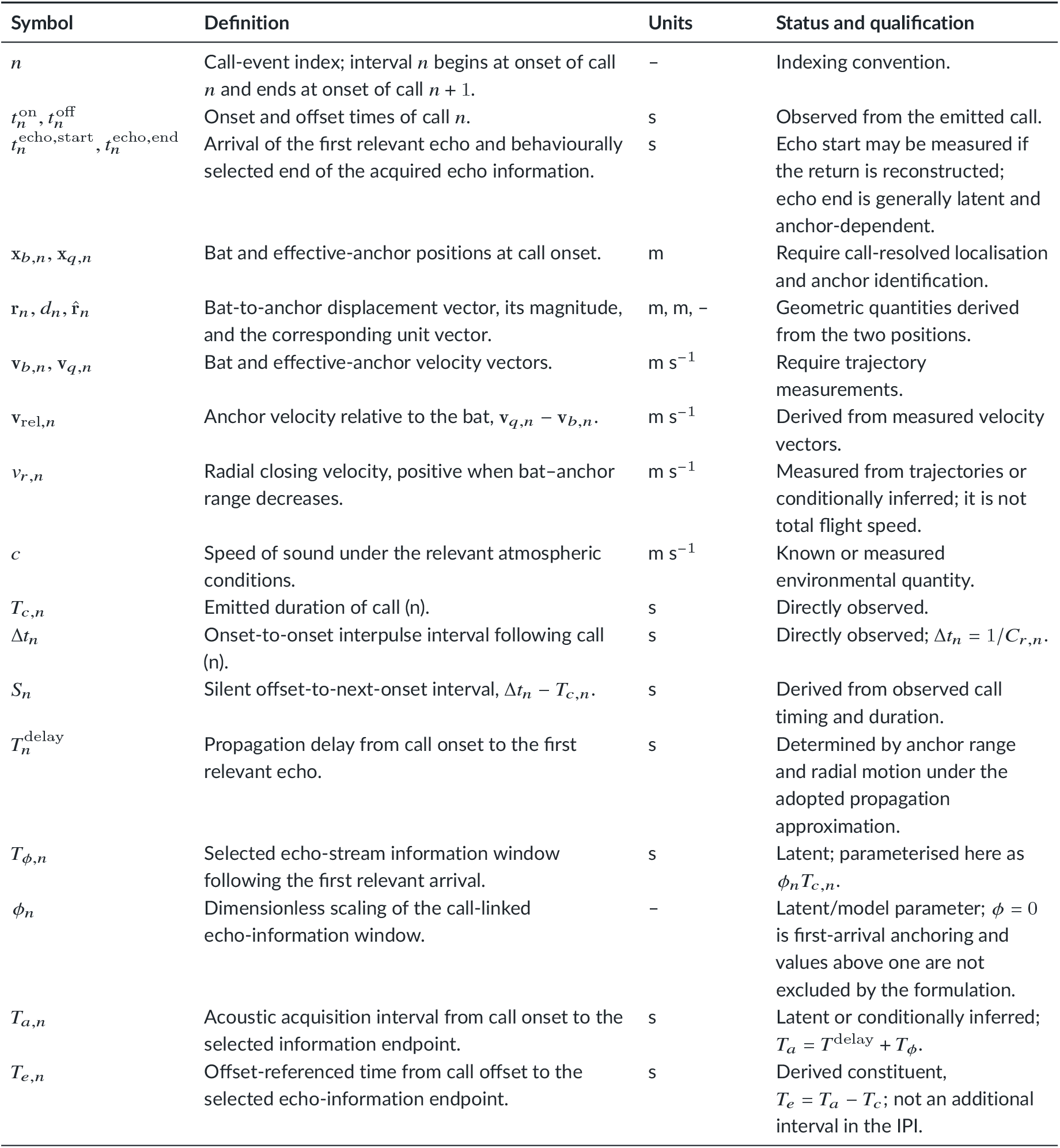
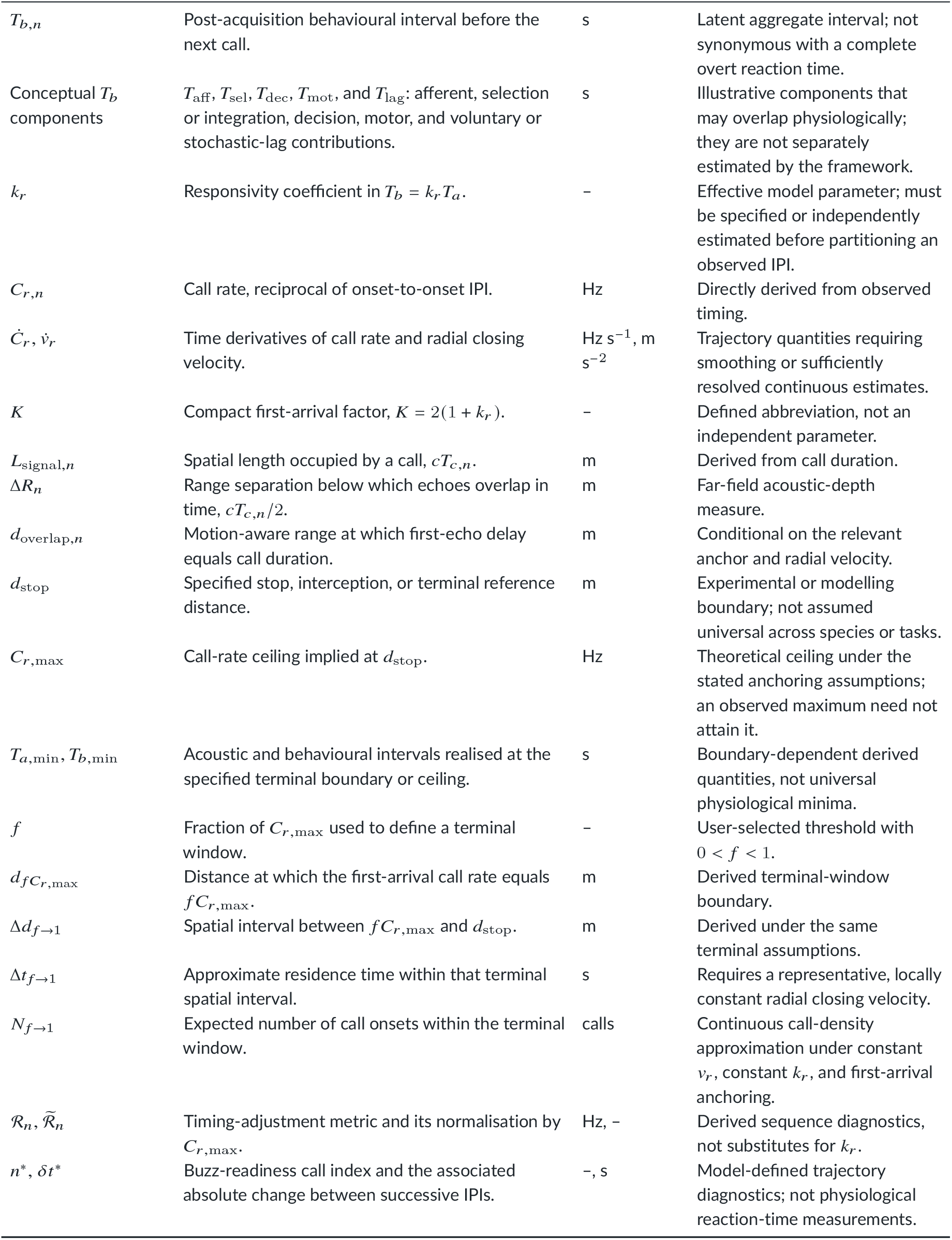
Core notation of the responsivity framework. “Observed” denotes quantities available from call timing or emitted signals; “measured” denotes quantities requiring spatial or kinematic observations; “latent” denotes quantities that can be inferred only after the required model parameters and anchor have been specified. All event-level quantities may vary with call index (n).

| Symbol | Definition | Units | Status and qualification |
| --- | --- | --- | --- |
| $n$ | Call-event index; interval $n$ begins at onset of call $n$ and ends at onset of call $n + 1$. | – | Indexing convention. |
| $t_n^{\text{on}}, t_n^{\text{off}}$ | Onset and offset times of call $n$. | s | Observed from the emitted call. |
| $t_n^{\text{echo,start}}, t_n^{\text{echo,end}}$ | Arrival of the first relevant echo and behaviourally selected end of the acquired echo information. | s | Echo start may be measured if the return is reconstructed; echo end is generally latent and anchor-dependent. |
| $\mathbf{x}_{b,n}, \mathbf{x}_{q,n}$ | Bat and effective-anchor positions at call onset. | m | Require call-resolved localisation and anchor identification. |
| $\mathbf{r}_n, d_n, \hat{\mathbf{r}}_n$ | Bat-to-anchor displacement vector, its magnitude, and the corresponding unit vector. | m, m, – | Geometric quantities derived from the two positions. |
| $\mathbf{v}_{b,n}, \mathbf{v}_{q,n}$ | Bat and effective-anchor velocity vectors. | $\text{m s}^{-1}$ | Require trajectory measurements. |
| $\mathbf{v}_{\text{rel},n}$ | Anchor velocity relative to the bat, $\mathbf{v}_{q,n} - \mathbf{v}_{b,n}$. | $\text{m s}^{-1}$ | Derived from measured velocity vectors. |
| $v_{r,n}$ | Radial closing velocity, positive when bat–anchor range decreases. | $\text{m s}^{-1}$ | Measured from trajectories or conditionally inferred; it is not total flight speed. |
| $c$ | Speed of sound under the relevant atmospheric conditions. | $\text{m s}^{-1}$ | Known or measured environmental quantity. |
| $T_{c,n}$ | Emitted duration of call ($n$). | s | Directly observed. |
| $\Delta t_n$ | Onset-to-onset interpulse interval following call ($n$). | s | Directly observed; $\Delta t_n = 1/C_{r,n}$. |
| $S_n$ | Silent offset-to-next-onset interval, $\Delta t_n - T_{c,n}$. | s | Derived from observed call timing and duration. |
| $T_n^{\text{delay}}$ | Propagation delay from call onset to the first relevant echo. | s | Determined by anchor range and radial motion under the adopted propagation approximation. |
| $T_{\phi,n}$ | Selected echo-stream information window following the first relevant arrival. | s | Latent; parameterised here as $\phi_n T_{c,n}$. |
| $\phi_n$ | Dimensionless scaling of the call-linked echo-information window. | – | Latent/model parameter; $\phi = 0$ is first-arrival anchoring and values above one are not excluded by the formulation. |
| $T_{a,n}$ | Acoustic acquisition interval from call onset to the selected information endpoint. | s | Latent or conditionally inferred; $T_a = T^{\text{delay}} + T_{\phi}$. |
| $T_{e,n}$ | Offset-referenced time from call offset to the selected echo-information endpoint. | s | Derived constituent, $T_e = T_a - T_c$; not an additional interval in the IPI. |

Supplementary Table S1 continued
| Symbol | Definition | Units | Status and qualification |
| --- | --- | --- | --- |
| $T_{b,n}$ | Post-acquisition behavioural interval before the next call. | s | Latent aggregate interval; not synonymous with a complete overt reaction time. |
| Conceptual $T_b$ components | $T_{\text{aff}}$, $T_{\text{sel}}$, $T_{\text{dec}}$, $T_{\text{mot}}$, and $T_{\text{lag}}$: afferent, selection or integration, decision, motor, and voluntary or stochastic-lag contributions. | s | Illustrative components that may overlap physiologically; they are not separately estimated by the framework. |
| $k_r$ | Responsivity coefficient in $T_b = k_r T_a$. | – | Effective model parameter; must be specified or independently estimated before partitioning an observed IPI. |
| $C_{r,n}$ | Call rate, reciprocal of onset-to-onset IPI. | Hz | Directly derived from observed timing. |
| $\dot{C}_r, \dot{v}_r$ | Time derivatives of call rate and radial closing velocity. | Hz s<sup>-1</sup>, m s<sup>-2</sup> | Trajectory quantities requiring smoothing or sufficiently resolved continuous estimates. |
| $K$ | Compact first-arrival factor, $K = 2(1 + k_r)$. | – | Defined abbreviation, not an independent parameter. |
| $L_{\text{signal},n}$ | Spatial length occupied by a call, $cT_{c,n}$. | m | Derived from call duration. |
| $\Delta R_n$ | Range separation below which echoes overlap in time, $cT_{c,n}/2$. | m | Far-field acoustic-depth measure. |
| $d_{\text{overlap},n}$ | Motion-aware range at which first-echo delay equals call duration. | m | Conditional on the relevant anchor and radial velocity. |
| $d_{\text{stop}}$ | Specified stop, interception, or terminal reference distance. | m | Experimental or modelling boundary; not assumed universal across species or tasks. |
| $C_{r,\text{max}}$ | Call-rate ceiling implied at $d_{\text{stop}}$. | Hz | Theoretical ceiling under the stated anchoring assumptions; an observed maximum need not attain it. |
| $T_{a,\text{min}}, T_{b,\text{min}}$ | Acoustic and behavioural intervals realised at the specified terminal boundary or ceiling. | s | Boundary-dependent derived quantities, not universal physiological minima. |
| $f$ | Fraction of $C_{r,\text{max}}$ used to define a terminal window. | – | User-selected threshold with $0 < f < 1$. |
| $d_{fC_{r,\text{max}}}$ | Distance at which the first-arrival call rate equals $fC_{r,\text{max}}$. | m | Derived terminal-window boundary. |
| $\Delta d_{f \rightarrow 1}$ | Spatial interval between $fC_{r,\text{max}}$ and $d_{\text{stop}}$. | m | Derived under the same terminal assumptions. |
| $\Delta t_{f \rightarrow 1}$ | Approximate residence time within that terminal spatial interval. | s | Requires a representative, locally constant radial closing velocity. |
| $N_{f \rightarrow 1}$ | Expected number of call onsets within the terminal window. | calls | Continuous call-density approximation under constant $v_r$, constant $k_r$, and first-arrival anchoring. |
| $\mathcal{R}_n, \tilde{\mathcal{R}}_n$ | Timing-adjustment metric and its normalisation by $C_{r,\text{max}}$. | Hz, – | Derived sequence diagnostics, not substitutes for $k_r$. |
| $n^*, \delta t^*$ | Buzz-readiness call index and the associated absolute change between successive IPIs. | –, s | Model-defined trajectory diagnostics; not physiological reaction-time measurements. |

**Table S2.** Simulation and variability notation. These quantities implement optional numerical constraints or stochastic variability and are not additional components of the core responsivity law.

| Symbol | Definition | Units | Role |
| --- | --- | --- | --- |
| $v_{r,n}^{\text{tim}}$ | Non-negative, subsonic radial velocity used only in the simulated acoustic-delay calculation. | m s<sup>-1</sup> | Numerical timing state; signed $v_r$ is retained for kinematics. |
| $\varepsilon_c$ | Small margin keeping $v_r^{\text{tim}} < c$. | m s<sup>-1</sup> | Numerical bound. |

Supplementary Table S2 continued
| Symbol | Definition | Units | Role |
| --- | --- | --- | --- |
| $d_0, v_0$ | Initial anchor distance and initial bat or approach speed specified for a simulated sequence. | m, m s<sup>-1</sup> | Initial-condition parameters; $v_0$ need not equal eventwise radial closing velocity. |
| $T_b^*$ | Optional minimum behavioural interval. | s | Response-floor parameter. |
| $T_{b,n}^{\text{core}}$ | Core interval after application of the optional response floor. | s | Equals $\max(k_r T_a, T_b^*)$. |
| $T_{b,n}^{\text{eff}}$ | Realised post-acquisition interval after optional caller-side lag. | s | Enters the emitted IPI. |
| $\delta_n, \bar{\delta}_n$ | Realised non-negative caller-side lag and its conditional mean. | s | Optional scheduling variability. |
| $f_\delta, f_{\delta, \max}$ | Mean and maximum lag as fractions of $T_b^{\text{core}}$. | – | Lag-scale parameters. |
| $\epsilon_n$ | Autoregressive deviation of caller-side lag from its mean. | s | Stochastic state. |
| $\rho_\delta$ | Lag-one autocorrelation of $\epsilon_n$. | – | Temporal persistence parameter. |
| $\sigma_{\delta, n}, s_\delta$ | Call-specific innovation standard deviation and its scale relative to mean lag. | s, – | Innovation-magnitude parameters. |
| $\eta_n$ | Independent standard-normal innovation driving caller-side lag. | – | Random variate. |
| $\Delta s_n^{\text{pred}}$ | Timing-predicted radial displacement, $v_r^{\text{tim}} \Delta t_n$. | m | Diagnostic, distinct from realised vector displacement. |
| $T_{c,0}, T_{c, \min}$ | Initial or maximum and minimum permitted call durations. | s | Duration bounds. |
| $g_c$ | Gain linking the next deterministic call duration to the preceding acquisition interval. | – | Causal duration-controller parameter. |
| $T_{c,n}^{\text{det}}, T_{c,n}^{\text{raw}}$ | Deterministic duration before jitter and unconstrained duration after jitter. | s | Intermediate simulation states. |
| $\sigma_c, p_c$ | Additive duration-jitter standard deviation and proportional jitter magnitude. | s, – | Duration-variability parameters. |
| $z_n$ | Standardised additive duration-jitter state. | – | Independent or autoregressive stochastic state. |
| $\rho_c$ | Lag-one autocorrelation coefficient of $z_n$. | – | Duration-jitter persistence. |
| $\xi_n, \zeta_n$ | Independent standard-normal innovations for correlated additive and separate proportional duration jitter. | – | Random variates. |

## S2 CONDITIONAL ANALYTICAL REFERENCE

For compactness, define the propagation-delay component implied by an observed interval under a specified *k*_*r*_ and *ϕ* as

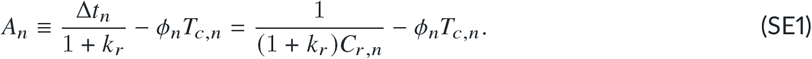

Under the motion-aware acquisition model, *A*_*n*_ = 2*d*_*n*_/(*c* + *v*_*r,n*_). A physically admissible positive-range reconstruction therefore requires *A*_*n*_ > 0. If *A*_*n*_ ≤ 0, the selected combination of *k*_*r*_, *ϕ*, timing, and call duration is incompatible with a positive propagation delay; assigning zero distance, as in the field diagnostic, is a computational boundary rather than a measurement of zero range.

**Table S3.**
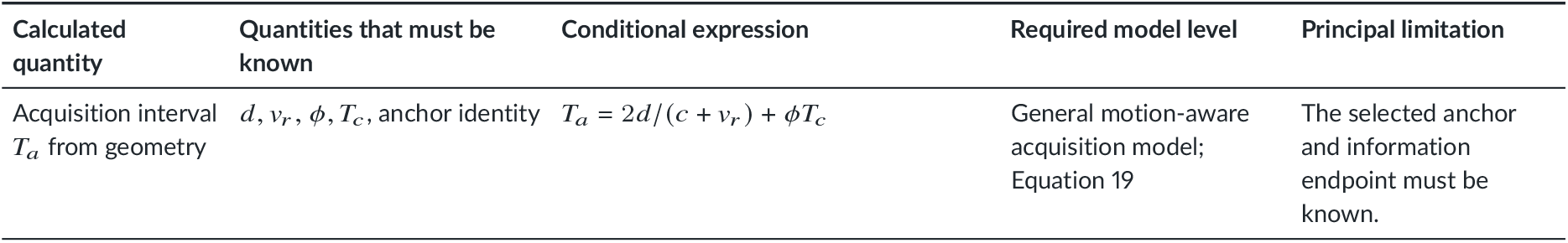
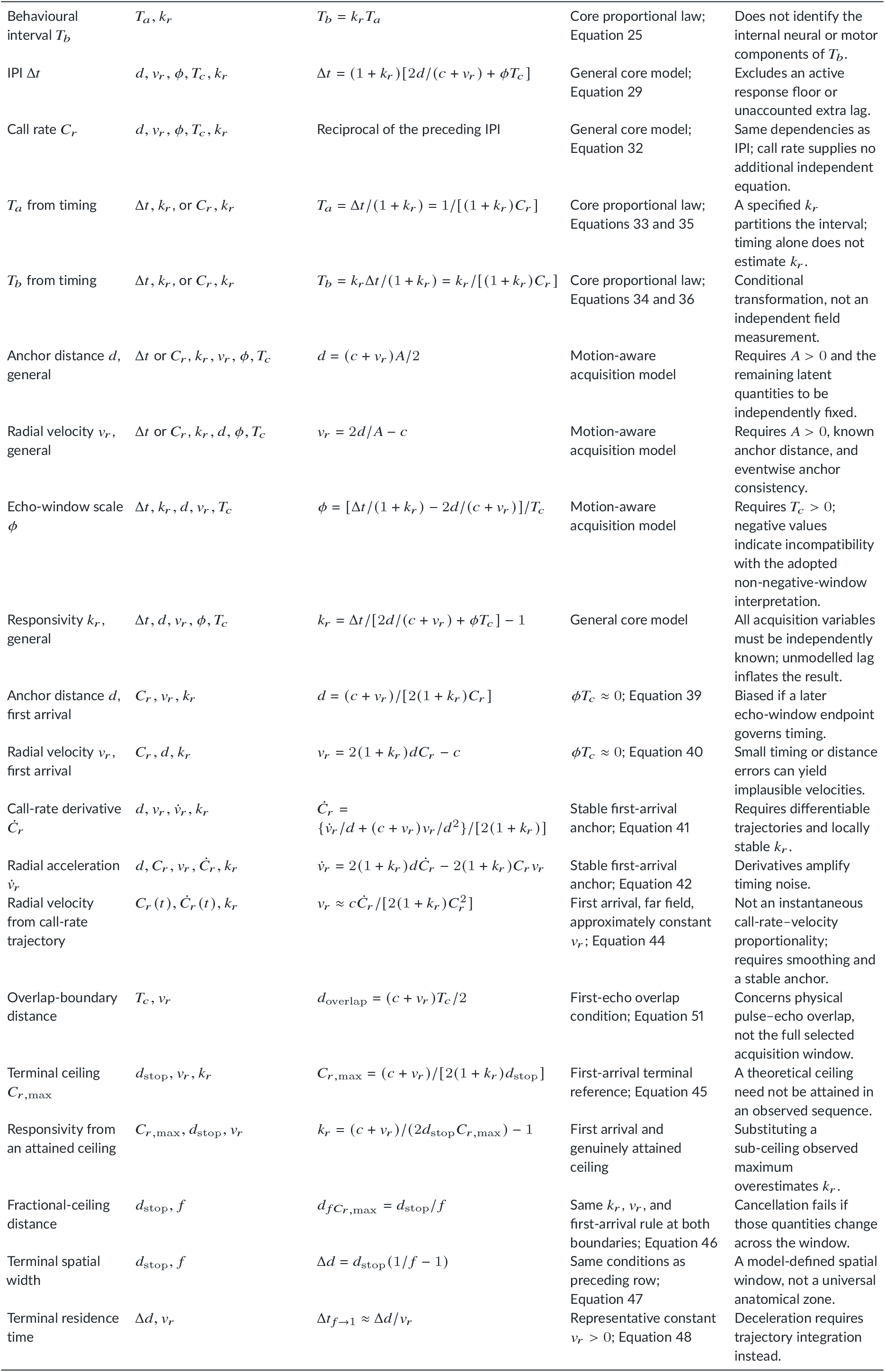
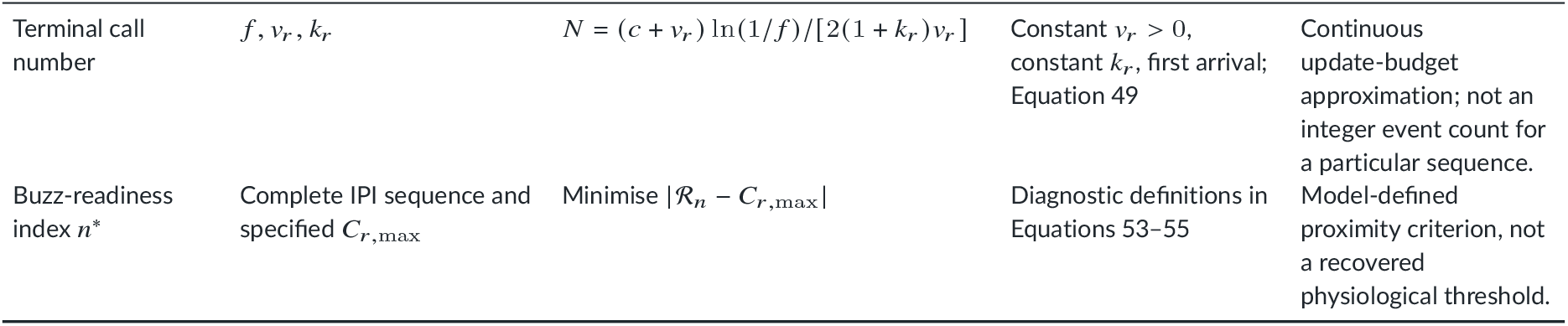
Analytical lookup table: what can be calculated from which known quantities. Each row is a conditional calculation, not a claim that the required inputs are ordinarily observable. Equivalent IPI and call-rate forms use Δ*t* = 1/*C*_*r*_. Main-text equation references identify the governing derivation.

**Table S4.**
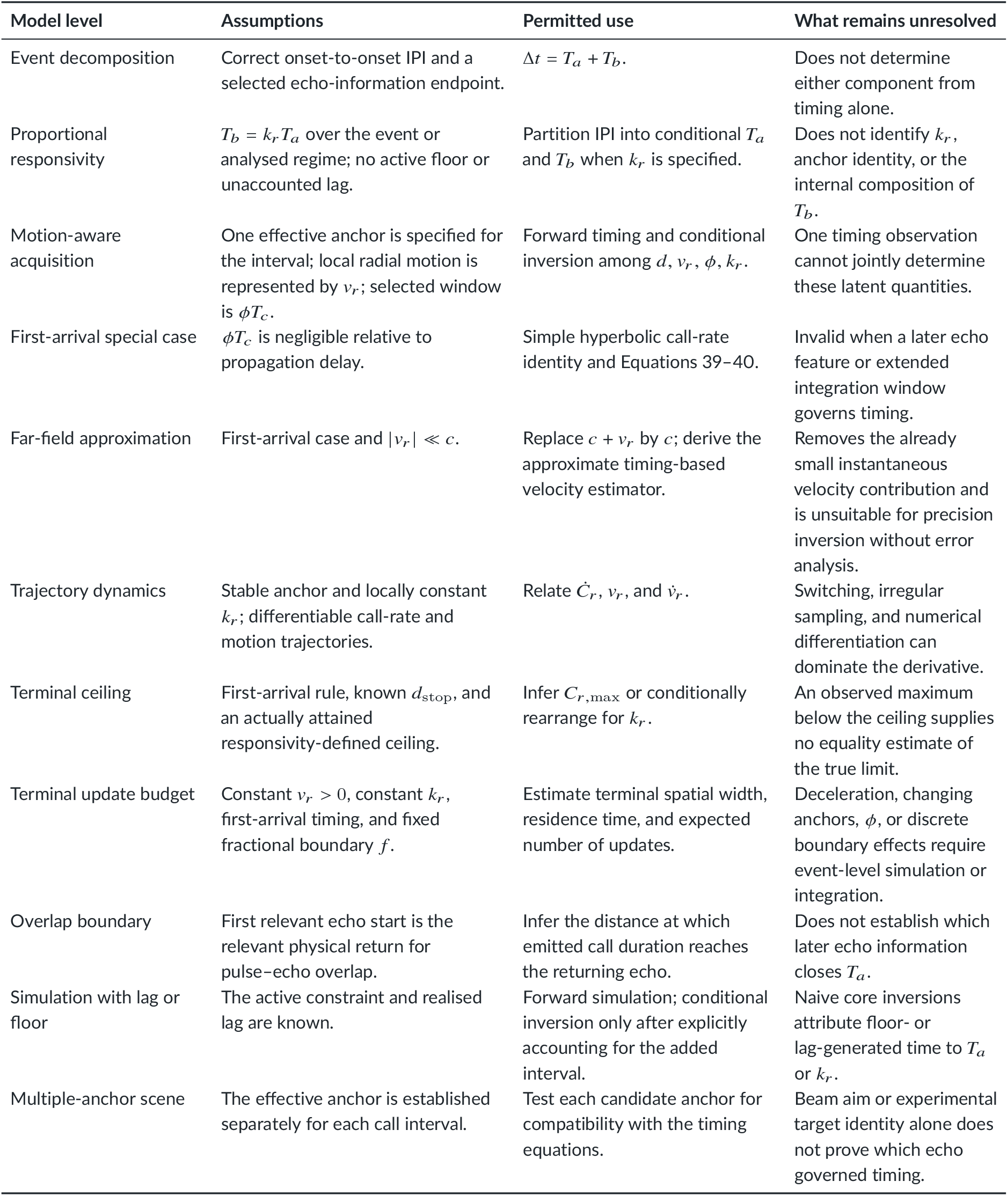
Assumption and identifiability map for the analytical equations. The rows identify which simplifications permit each family of calculations and what cannot be concluded from them.

## S3 SELECTED DERIVATIONS AND SAFEGUARDS

### S3.1 General inverse family

Combining Equations 29 and SE1 gives *A*_*n*_ = 2*d*_*n*_/(*c* + *v*_*r,n*_). Consequently, any one of *d*_*n*_, *v*_*r,n*_, *ϕ*_*n*_, or *k*_*r*_ can be calculated only when the other acquisition variables are independently specified:

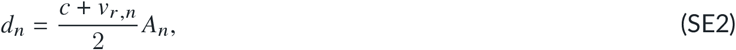

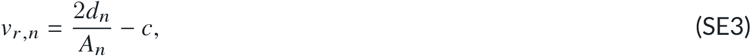

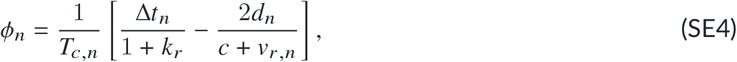

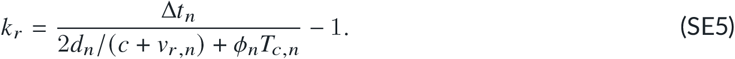

These rearrangements are algebraically exact within the core motion-aware model, but they are not jointly identifying. One observed IPI supplies one timing constraint. If several of the anchor distance, radial velocity, echo-window endpoint, and responsivity coefficient are unknown, infinitely many combinations can explain the same interval. Moreover, if a known additional lag δ_*n*_ is present, Δ*t*_*n*_ − δ_*n*_, rather than Δ*t*_*n*_, enters the proportional core inversion. When the response floor is active, *T*_*b*_ ≠ *k*_*r*_*T*_*a*_ and this inverse family cannot recover *k*_*r*_ from that event.

### S3.2 First-arrival and terminal rearrangements

When *ϕT*_*c*_ is negligible, *A* = Δ*t*/(1 + *k*_*r*_) and the general inverse family reduces to Equations 39 and 40. At a known stop distance, rearranging Equation 45 gives

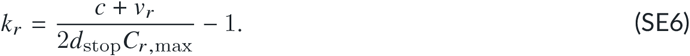

This is an equality only when the observed rate is the attained terminal ceiling under the assumed anchor and first-arrival rule. If the largest recorded call rate lies below the attainable ceiling, substituting it into Equation SE6 overestimates *k*_*r*_ under otherwise correct assumptions.

For constant radial velocity, the terminal call budget follows by integrating call density per unit distance over the interval from *d*_stop_ to *d*_stop_/ *f* :

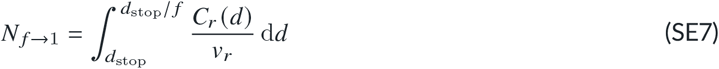

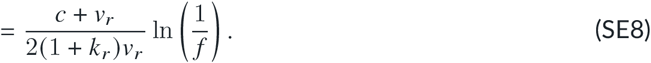

The cancellation of *d*_stop_ in this expression does not make the stopping geometry irrelevant: that distance fixes the physical placement and width of the terminal window. The closed form applies only when *v*_*r*_, *k*_*r*_, and the anchoring rule remain constant across it; otherwise the event-level trajectory must be integrated directly.

### S3.3 Trajectory derivative

For the first-arrival identity *C*_*r*_ = (*c* + *v*_*r*_)/(*Kd*), differentiation with *d* = −*v*_*r*_ gives

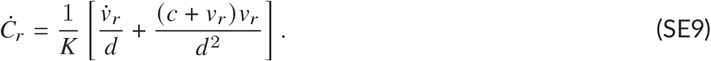

If 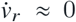, substituting *d* ≈ *c*/(*KC*_*r*_) in the far field yields Equation 44. Thus, velocity is related to the trajectory derivative of call rate, not directly proportional to instantaneous call rate. Anchor changes or noisy numerical differentiation can violate the conditions needed for this estimator even when individual calls remain compatible with the general timing equation.

### S4 SECONDARY NUMERICAL RESULTS

**Table S5.**
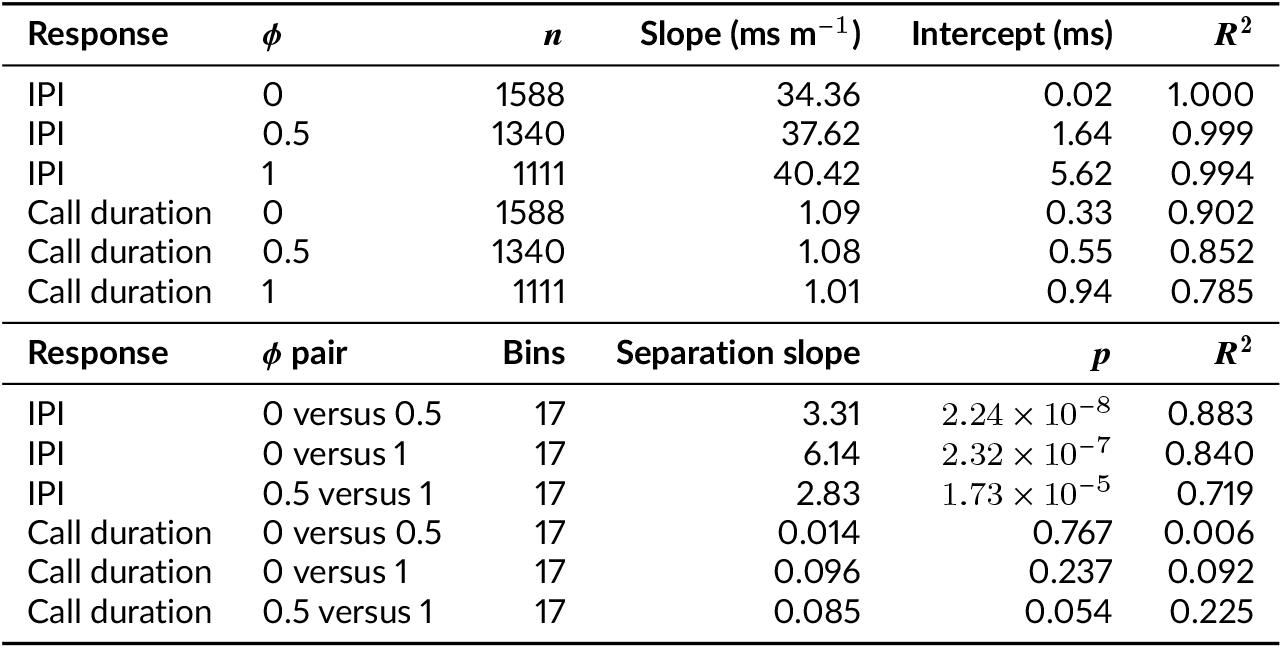
Echo-window effects on IPI, call duration, and call-rate spread. Slopes and intercepts are from per-condition fits against anchor distance. Pairwise slopes describe the change in absolute separation between binned median curves per metre; positive values indicate convergence towards the target. For IPI, the omnibus effects of distance, *ϕ*, and their interaction were all *p* ≈ 0 (*R*^2^ = 0.997); for call duration, the corresponding model had *R*^2^ = 0.855. In the call-rate bend window, mean IQRs were 8.80, 7.81, and 7.28 Hz for *ϕ* = 0, 0.5, and 1; neither *ϕ* (*p* = 0.890) nor its interaction with distance (*p* = 0.955) explained spread.

**Table S6.** Interval–distance fits for the responsivity, echo-window, and velocity sweeps. Slopes and intercepts are given separately for *T*_*a*_ and *T*_*b*_. The nested interaction tests detected distance, condition, and distance-by-condition effects in every sweep (*p*≈ 0); the near-identical *T*_*a*_ slopes across *k*_*r*_, and the small changes across velocity, show why effect magnitude rather than significance is decisive here. For the *ϕ* = 0 versus 1 curves, binned separation increased with distance at 0.48 ms m^−1^ for *T*_*a*_ and 2.38 ms m^−1^ for *T*_*b*_ (*p* < 0.001).

| Sweep | Level | $n$ | $T_a$ slope | $T_a$ intercept | $T_b$ slope | $T_b$ intercept | $R^2$ |
| --- | --- | --- | --- | --- | --- | --- | --- |
| $k_r$ | 3 | 1867 | 5.99 | 0.65 | 17.96 | 1.94 | 0.998 |
| $k_r$ | 5 | 1283 | 6.05 | 0.37 | 30.23 | 1.85 | 0.999 |
| $k_r$ | 7 | 955 | 6.05 | 0.25 | 42.38 | 1.74 | 0.999 |
| $k_r$ | 9 | 759 | 6.05 | 0.19 | 54.43 | 1.73 | 0.999 |
| $\phi$ | 0 | 1433 | 5.74 | 0.00 | 28.72 | 0.01 | 1.000 |
| $\phi$ | 0.5 | 1274 | 6.04 | 0.37 | 30.22 | 1.85 | 0.999 |
| $\phi$ | 1 | 1106 | 6.29 | 0.97 | 31.44 | 4.83 | 0.995 |
| $v_0$ (m s<sup>-1</sup>) | 2 | 3292 | 6.13 | 0.29 | 30.64 | 1.46 | 0.999 |
| $v_0$ (m s<sup>-1</sup>) | 4 | 1624 | 6.07 | 0.36 | 30.35 | 1.78 | 0.999 |
| $v_0$ (m s<sup>-1</sup>) | 6 | 1042 | 6.02 | 0.39 | 30.08 | 1.96 | 0.999 |
| $v_0$ (m s<sup>-1</sup>) | 8 | 762 | 5.96 | 0.49 | 29.79 | 2.44 | 0.999 |
| $v_0$ (m s<sup>-1</sup>) | 10 | 602 | 5.91 | 0.53 | 29.54 | 2.65 | 0.999 |

**Table S7.** Secondary statistics for the field–simulation comparison and conditional profiles. The first three rows compare measured quantities; subsequent rows are conditional transformations or profile diagnostics under fixed *k*_*r*_ = 5 and the stated simulation geometry. They must not be interpreted as independent measurements of target distance, *T*_*b*_, or *ϕ*. The initial-distance sensitivity sweep covered *d*_0_ = 2–8 m; both diagnostics retained the global minimum at *d*_0_ = 5 m and *ϕ* = 1.5 on the coarser grid, although the preferred *ϕ* varied with *d*_0_.

| Quantity | Field | Simulation | Comparison or profile result |
| --- | --- | --- | --- |
| Effective velocity ($\text{m s}^{-1}$) | 6.461 (5.239–7.593) | 6.460 (5.209–7.606) | Velocity-band occupancy:<br>$\chi^2 = 0.011$, $df = 4$,<br>$p = 0.999985$ |
| IPI interaction model | 4425 calls | 4427 calls | Dataset-dependent terms $p \approx 0$;<br>$R^2 = 0.969$ |
| Duration-envelope compatibility | 20 seeded ensembles |  | Median 94.1% (IQR 93.1–95.0%) |
| Distance profile | 1.668 m (1.074–1.946) | 1.259 m (0.727–1.983) | Minimum at $\phi = 1.35$; mean<br>quantile gap 0.362 m; 1.79%<br>clipped at zero |
| $T_b$ profile | 68.59 ms (50.05–77.03) | 49.99 ms (32.09–74.83) | Minimum at $\phi = 1.20$; mean<br>quantile gap 16.83 ms |
| Rank-matched required $\phi$ | Median 1.76 (IQR 1.05–2.22) | | Conditional diagnostic, not<br>parameter recovery |
| Initial-distance sensitivity | Candidate $d_0 = 2\text{--}8$ m | | Global minimum retained at<br>$d_0 = 5$ m; best $\phi = 1.5$ on<br>coarse grid |

**Table S8.** Buzz-readiness summaries across responsivity, velocity, and initial call duration. Values are grouped medians; parentheses give the interquartile range for readiness distance. Each *k*_*r*_ level contains 400 sequences, each velocity level 320 sequences, and each duration level 400 sequences.

| Sweep | Level | Readiness distance (m) | $\delta t^*$ (ms) | Calls per sequence |
| --- | --- | --- | --- | --- |
| $k_r$ | 3 | 1.98 (1.43–3.16) | 3.52 | 47.5 |
| $k_r$ | 5 | 1.37 (1.01–2.00) | 5.43 | 32 |
| $k_r$ | 7 | 1.02 (0.70–1.45) | 6.78 | 23 |
| $k_r$ | 9 | 0.80 (0.52–1.19) | 8.72 | 19 |
| $v_0$ ($\text{m s}^{-1}$) | 1 | 3.55 (2.75–4.54) | 6.10 | 85 |
| $v_0$ ($\text{m s}^{-1}$) | 2 | 1.72 (1.32–2.49) | 6.00 | 42 |
| $v_0$ ($\text{m s}^{-1}$) | 3 | 1.19 (0.91–1.67) | 6.37 | 27.5 |
| $v_0$ ($\text{m s}^{-1}$) | 4 | 0.85 (0.61–1.22) | 6.12 | 20 |
| $v_0$ ($\text{m s}^{-1}$) | 5 | 0.66 (0.52–0.98) | 5.96 | 16 |
| $T_{c,0}$ (ms) | 3 | 1.29 (0.80–2.25) | 6.01 | 30 |
| $T_{c,0}$ (ms) | 5 | 1.28 (0.80–2.24) | 5.98 | 30 |
| $T_{c,0}$ (ms) | 7 | 1.28 (0.80–2.24) | 5.97 | 30 |
| $T_{c,0}$ (ms) | 10 | 1.28 (0.79–2.24) | 5.96 | 30 |

**Table S9.**
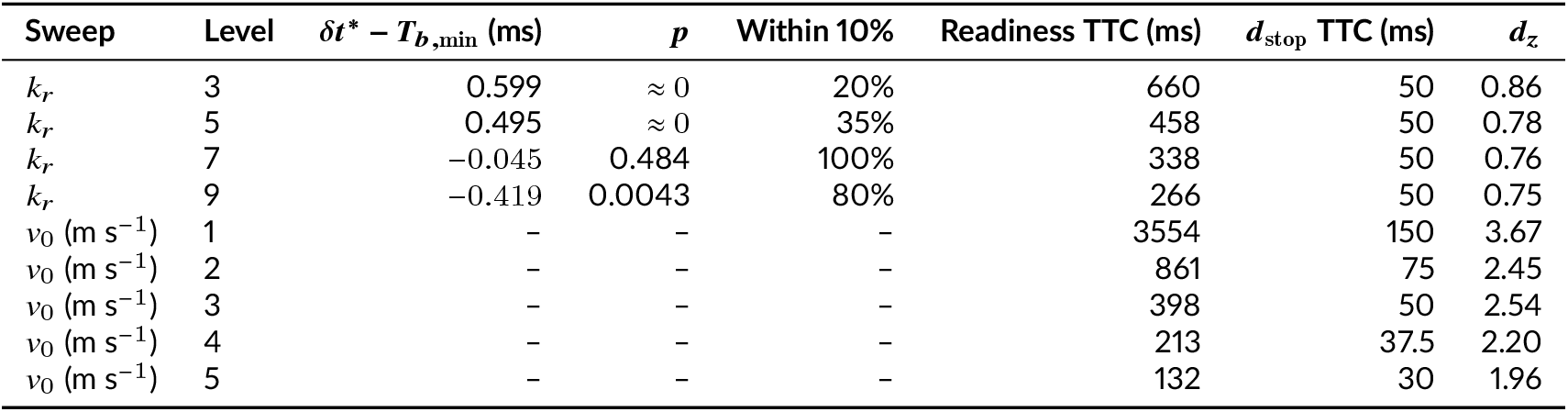
Proxy and time-to-contact (TTC) comparisons for buzz readiness. The *δt*^∗^ −*T*_*b*,min_ columns apply to the *k*_*r*_ sweep. TTC columns compare the median remaining time from readiness with the time from *d*_stop_. Paired tests for all TTC comparisons gave *p* ≈ 0 and 0% agreement within 10%.

**Table S10.** Structure of SSG-like events in the productive multi-target conditions. Stable IPI and anchor distance are means with SD and interquartile range. Group-size counts are given as doublets/triplets/quadruplets.

| Condition | Events | Group sizes | Stable IPI (ms) | Anchor distance (m) | Same anchor |
| --- | --- | --- | --- | --- | --- |
| C3: stationary, random anchor | 78 | 77/1/0 | 10.30 (5.30; 7.27–11.80) | 0.296 (0.152; 0.209–0.339) | 100% |
| C6: moving, random anchor | 438 | 405/32/1 | 13.14 (7.14; 8.78–14.89) | 0.378 (0.205; 0.252–0.427) | 98.4% |

